# Genome-Wide DNA Methylation Profiling of the Failing Human Heart with Mechanical Unloading Identifies *LINC00881* as an Essential Regulator of Calcium Handling in the Cardiomyocyte

**DOI:** 10.1101/2022.03.01.482163

**Authors:** Xianghai Liao, Peter J. Kennel, Bohao Liu, Trevor R. Nash, Richard Zhuang, Amandine F. Godier-Furnemont, Chenyi Xue, Rong Lu, Paolo C. Colombo, Nir Uriel, Muredach P. Reilly, Steven O. Marx, Gordana Vunjak-Novakovic, Veli K. Topkara

**Author notes:** **Corresponding Author:** Veli K. Topkara, M.D., M.Sc., Assistant Professor of Medicine, Center for Advanced Cardiac Care, Columbia University Medical Center – New York Presbyterian, 622 West 168^th^ St, PH10-203A. These authors contributed equally to this work (Xianghai Liao, Peter J. Kennel). **Industry Relationships:** All other authors have reported that they have no relationships relevant to the contents of this paper to disclose.

## Abstract

**Background:** Human heart failure is characterized by global alterations in the myocardial DNA methylation profile, yet little is known about epigenetic regulation of non-coding transcripts and potential reversibility of DNA methylation with left ventricular assist device (LVAD) support.

**Method:** Genome-wide mapping of myocardial DNA methylation was performed in 36 patients with end-stage heart failure at the time of LVAD implant, 8 patients at the time of LVAD explant, and 7 non-failing controls using high-density bead array platform. Transcriptomic and functional studies were performed in human induced pluripotent stem cell derived cardiomyocytes (iPSCs).

**Results:** Etiology-specific analysis revealed 2079 differentially methylated positions (DMPs) in ischemic cardiomyopathy (ICM) and 261 DMPs in non-ischemic cardiomyopathy (NICM). 192 DMPs were common to ICM and NICM. Analysis of paired samples before and after LVAD support demonstrated reverse methylation of only 3.2% of HF-specific DMPs. Methylation-expression correlation analysis yielded several protein-coding genes that are hypomethylated and upregulated (*HTRA1, FAM65A, FBXO16, EFCAB13, AKAP13, RPTOR*) or hypermethylated and downregulated (*TBX3*) in ICM and NICM patients. A novel cardiac-specific super-enhancer lncRNA (*LINC00881*) is hypermethylated and downregulated in the failing human heart. *LINC00881* is an upstream regulator of sarcomere and calcium channel gene expression including *MYH6, CACNA1C*, and *RYR2. LINC00881* knockdown significantly reduced peak calcium amplitude in the beating human iPSCs.

**Conclusions:** Failing human heart exhibits etiology-specific changes in DNA methylation including coding and non-coding regions, which are minimally reversible with mechanical unloading. Epigenetic reprogramming may be necessary to achieve transcriptional normalization and sustained clinical recovery from heart failure.

## Introduction

Heart failure (HF) is a major public health problem with more than 6 million patients affected in the United States alone.^1^ Despite the tremendous progress achieved in diagnosis and treatment of HF, the majority of patients progress to advanced stages of the disease leading to unacceptably high rates of morbidity and mortality, even exceeding most cancers. Pharmacological management of HF has traditionally focused on targeting endogenous neurohormonal signaling cascades that are associated with disease progression as well as diuretic therapy for symptom relief.^2^ Yet, our limited mechanistic understanding of the complex HF pathophysiology does not fully explain the clinical heterogeneity and wide variation in the treatment response observed in this population, highlighting the need for novel molecular diagnostics as well as therapeutic strategies.

The failing human heart undergoes structural and functional remodeling that is accompanied by profound alterations in the myocardial transcriptome including recapitulation of the fetal gene expression program and downregulation of the genes involved in the oxidative phosphorylation pathway.^3, 4^ While some of these changes are common to all forms of heart failure, gene expression could be etiology-specific and help distinguish patients with different types of HF.^5, 6^ Transcriptional profiling of paired myocardial samples obtained from HF patients before and after left ventricular assist device (LVAD) support demonstrate that only a small percentage of genes that are dysregulated in HF normalize with mechanical unloading of the failing human heart, suggesting that normalization of dysregulated HF transcriptional program could be necessary to achieve sustained clinical recovery.^7-9^ Several transcription factors (TFs) including GATA-4, NFAT, MEF2, and Nkx-2.5 have been implicated in maladaptive cardiac remodeling, however emerging evidence suggest that epigenetic regulation may play important roles in transcriptional reprogramming by altering gene accessibility and TF binding to gene promoters or enhancers.^10, 11^

DNA methylation is an essential epigenetic modification involving transfer of a methyl group onto the fifth position of the cytosine catalyzed by a family of DNA methyltransferases (DNMTs) and generally signals for transcription repression at the promoter sites.^12^ Several pathological conditions, in particular malignant transformation, have been associated with hyper-methylation of specific target gene promoters as well as global hypomethylation leading to genomic instability.^13^ DNA methylation assays have been FDA approved for early detection of cancer and have the potential to predict disease course and response to therapy.^14, 15^ Moreover, DNA demethylation agents have become standard of care therapy for high-risk patients with myelodysplastic syndrome through epigenetic re-activation of tumor suppressor genes.^16^ While the data is lagging for HF, several studies to date investigated genome-wide DNA methylation in the failing human heart using targeted bisulphite sequencing or chip-based approaches and identified differentially methylated regions in the genome.^17-23^ However, none of these studies have examined whether and if so to what extent aberrant DNA methylation is reversible in the failing human myocardium. Moreover, the relationship between DNA methylation and the transcription of non-coding genomic elements with regulatory function including long non-coding RNAs (lncRNAs) remains largely unknown. Hence, in the current study we characterize the myocardial DNA methylation profile of patients with Ischemic (ICM) and Non-Ischemic (NICM) HF, and the impact of mechanical unloading on DNA methylation profile in paired myocardial tissue obtained before and after LVAD support. Furthermore, we identify a novel cardiac-specific and super-enhancer associated long-noncoding RNA (*LINC00881*), that is hypermethylated and transcriptionally downregulated in the failing human heart, as a key regulator of calcium handling in the cardiomyocyte.

## Material and Methods

### Patient Enrollment

Myocardial DNA methylation profiling was performed 36 patients with end-stage heart failure (12 Ischemic [ICM] and 24 Non-Ischemic [NICM] dilated cardiomyopathy, n=36 pre-LVAD) who underwent LVAD implantation at Columbia University Irving Medical Center including 8 patients with paired cardiac tissue at the time of heart transplantation (n= 8 post-LVAD) and non-failing control cardiac tissue (n=7, non-failing) obtained from the National Disease Research Interchange (NDRI) in Philadelphia, PA, USA. Informed consent was obtained for the procurement of discarded apical core myocardial tissue at the time of LVAD implantation and cardiac transplantation. The study was approved by Columbia University Irving Medical Center Institutional Review Board.

### Genome-wide DNA Methylation Profiling

Apical core myocardial samples were washed in ice-cold saline (0.9% NaCl) and stored in liquid nitrogen until DNA and RNA were extracted. DNA was extracted using a DNA Kit (Qiagen). Purity and concentration were determined using a Bioanalyzer (Agilent Technologies, CA). For DNA methylation profiling, DNA was bisulfite converted using a commercially available kit (Zymo Research, Orange, CA). Illumina Infinium Human Methylation 450K Bead Chip and EPIC Bead Chip were used for genome-wide profiling of DNA methylation.^24^ The EPIC chip measures over 850 000 methylation sites, with high reproducibility in comparison to the previous 450k chip.^25^ After whole genome DNA amplification, the samples were applied following the Illumina Infinium Human Methylation DNA chip manufacturer protocols (Illumina, San Diego, CA). Chips were analyzed using the Illumina Hi-Scan system at the McDonnell Genome Institute (MGI) at Washington University.

### Bioinformatics Analysis

Integrated analysis of the 450K and EPIC Bead Array data was conducted using Bioconductor package ChAMP in R.^26^ The analysis pipeline has been summarized in **Fig. S1**. Briefly, raw methylation data was imported into R and normalized using BMIQ. 450K and EPIC data was merged on common probes and corrected for batch effects using combat function. Additional filtering was performed for common SNPs. Differentially methylated positions (DMP) were identified using a cut-off of 10% change in methylation and q value <0.05. The first analysis focused on HF etiology and identified DMPs associated with ICM (ICM vs. NF) and NICM (NICM vs. NF) using data from pre-LVAD tissue obtained from 36 patients and 7 non-failing controls. The second analysis focused on the impact of mechanical unloading, and identified DMPs that are associated with HF (pre-LVAD vs. NF) and reverse remodeling (post-LVAD vs. pre-LVAD) in 8 paired cardiac tissue samples. The variance in global DNA methylation between subjects was assessed using principal component analysis (PCA) plots using the first three principal components. Hierarchical clustering of subjects was performed using the complete linkage method. DMPs common to ICM and NICM were screened for reciprocal changes in gene expression. RNA-sequencing data from left ventricular samples of 50 heart failure patients (13 with Ischemic and 37 with Non-Ischemic cardiomyopathy) and 14 non-failing controls was obtained from GEO (Accession number: 116250) and analyzed using the limma package in R. 36,755 transcripts with RPKM level of 0 in more than 50% of patients were filtered out. Data was log transformed using (RPKM + 1). Differential expression analysis was performed using linear model ANOVA. Differentially expressed genes (DEGs) were defined as genes with a *p*-value adjusted for Benjamini-Hochberg false discovery rate (FDR) ≤ 0.05 between heart failure and control samples. The locations of human cardiac super-enhancers were acquired from previous publications.^27, 28^ ChIP-seq data is available through the Gene Expression Omnibus (GEO) using the following accession numbers: adult human heart H3K27ac (GSE101345), adult human heart H3K4me1 (GSE101156), adult human heart H3K27me3 (GSE101387), adult human heart CTCF (GSE 127553). DMPs located within intergenic regions were mapped to lnc-RNAs using GENCODE annotation database.^29^

### Cell Culture and Human Cardiomyocyte Differentiation

Induced pluripotent stem cells were obtained through material transfer agreements with Dr. Vunjak-Novakovic from B. Conklin, Gladstone Institute (WTC-11, healthy). Cells were maintained on 1:20 diluted growth factor reduced Matrigel (Corning, Corning, NY) in mTeSR-plus medium (StemCell Technologies, Vancouver, CA) supplemented with 1% penicillin/streptomycin (ThermoFisher Scientific, Waltham, MA) at 37°C, 21% O_2_. iPS cells were passaged at 30–50% confluence using 0.5 mM EDTA (ThermoFisher Scientific, Waltham, MA) and cultured for 24 hours in iPS media supplemented with 5 μM Y-27632 (Tocris Biosciences, Bristol, UK) prior to maintenance in iPS media. Cells were used between passages 40 and 70.

Cardiac differentiation of human iPS cells was performed using a stage-based protocol in RPMI-1640 (ThermoFisher Scientific, Waltham, MA) supplemented with 0.5 mg/mL recombinant human albumin (Sigma-Aldrich, St. Louis, MO), 213 μg/mL L-ascorbic acid 2-phosphate (Sigma-Aldrich, St. Louis, MO), and 1% penicillin/streptomycin (CM media). iPS cells were grown to 80– 90% confluence and changed into CM Media supplemented with 3 μM CHIR99021 (Tocris Biosciences, Bristol, UK) for 2 days. Media was then changed to CM Media supplemented with 2 μM Wnt-C59 (Tocris Biosciences, Bristol, UK) for 2 days prior to switching to CM Media without any supplements. CM Media is changed every 48 hours until contracting cells were noted by around day 10 following the initiation of differentiation, at which time the medium was changed to RPMI 1640 supplemented with B27 (50X; Gibco). Experiments were performed using cells at day 15 – 35.

### LINC00881 Overexpression and Knockdown in Human iPS cell derived Cardiomyocytes

Beating human iPSCs were switched to Opti-MEM™ Reduced Serum Medium (ThermoFisher Scientific, Waltham, MA). *LINC000881* knockdown was achieved by lipofectamine based transfection of antisense LNA GapmeRs designed against *LINC00881* versus scrambled oligonucleotide (QIAGEN, Germantown, MD) using RNAiMAX reagent (ThermoFisher Scientific, Waltham, MA) (**Table S14**). iPSCs were harvested at 48-72 hours following transfection for functional and gene expression studies. Knockdown of LINC00881 was confirmed by qPCR. For LINC00881 overexpression experiment, full length human LINC00881 was cloned into p3XFLAG-CMV-7 (Millipore Sigma, St. Louis, MO, USA) vector and amplified in DH5α strain of *Escherichia coli* (*E coli*) (Life Technologies, Grand Island NY). After amplification, plasmids were extracted through QIAGEN Plasmid Midi Kit (QIAGEN, Germantown, MD) and stored at −80 °C until use. Beating human iPSCs were switched to Opti-MEM™ Reduced Serum Medium (ThermoFisher Scientific, Waltham, MA). *LINC000881* overexpression was achieved by lipofectamine based transfection of *LINC00881* plasmid versus bacterial alkaline phosphatase control plasmid using RNAiMAX reagent (ThermoFisher Scientific, Waltham, MA). iPSCs were harvested at 48-72 hours following transfection for functional and gene expression studies. Overexpression of LINC00881 was confirmed by qPCR.

### Quantitative Real-time PCR

Total RNA was isolated from LV apical core tissue and from beating human iPS cell derived cardiomyocytes using Quick-RNA Miniprep Plus (Zymo Research, Irvine, CA, USA). The qRT-PCR was performed using SYBR mix (Thermofisher) on PicoReal96 Real-time PCR Systems (Thermoscintific). Transcript quantification for mRNAs were performed using delta-delta method using forward and reverse primers designed specifically for each of the target mRNAs and lncRNAs. The primer sequences used for independent validation are listed in **Table S13**. 18S was used as internal control.

### RNA sequencing and Data Analysis

RNA concentration and integrity were assessed using a 2100 BioAnalyzer (Agilent, Santa Clara CA). Sequencing libraries were constructed using the TruSeq Stranded Total RNA Library Prep Gold mRNA (Illumina, San Diego CA) with an input of 1000 ng and 11 cycles final amplification. Final libraries were quantified using High Sensitivity D1000 ScreenTape on a 2200 TapeStation (Agilent, Santa Clara CA) and Qubit 1x dsDNA HS Assay Kit (Invitrogen, Waltham MA). Samples were pooled equimolar with sequencing performed on an Illumina NovaSeq6000 SP 300 Cycle Flow Cell v1.5 as Paired-end 151 reads. Sequences from fastq files were aligned to reference human genome (Gencode v38) using the splice-aware aligner STAR v.2.7.3a. Differential gene expression analysis with LINC00881 knockdown or overexpression was performed using the R package DESeq2 (v1.34.0) from unnormalized count data.

### Human iPS Cell derived Cardiomyocyte Calcium Imaging

Human iPS cell derived cardiomyocyte monolayers were loaded with 5μM Calbryte-590 in RPMI + B27 medium at 37°C for 45 minutes. Loading media was washed and replaced with Tyrode’s salt solution. Calcium transience videos were acquired at 50 frames per second using a sCMOS camera (Zyla 4.2, Andor Technology) connected to an inverted fluorescence microscope (IX-81, Olympus) with cells placed in a live-cell chamber (STX Temp & CO_2_ Stage Top Incubator, Tokai Hit). Calcium signal analysis was performed using custom Python script as previously described.^30^

### Statistical Analysis

Statistical analyses were performed using R (R Core Team, 2021). Continuous data are presented as mean ± standard deviation. For comparisons between two groups, a two tailed unpaired t-test was used. p value less than 0.05 was considered statistically significant in all analyses.

## Results

### Myocardial DNA Methylation in Ischemic and Non-Ischemic Cardiomyopathy

36 patients with end-stage heart failure (12 Ischemic [ICM] and 24 Non-Ischemic [NICM] dilated cardiomyopathy) who underwent LVAD implantation at Columbia University Irving Medical Center and 7 non-failing controls were included for genome-wide DNA methylation analysis. Patients with myocarditis, amyloid cardiomyopathy, restrictive/hypertrophic cardiomyopathy, and previous cardiac transplantation were excluded. Post-LVAD cardiac samples were obtained from 8 patients who were bridged to heart transplantation. Clinical information obtained at the time of tissue procurement and during LVAD support was summarized in **Table S1**.

Genome-wide DNA methylation profiling demonstrated 2079 differentially methylated positions (DMPs) in myocardium of patients with ischemic cardiomyopathy (ICM vs. NF, q <0.05, **Table S2**). Of those 625 DMPs were hypermethylated and 1454 DMPs were hypomethylated. 261 DMPs were differentially methylated in myocardium of patients with non-ischemic cardiomyopathy (NICM vs. NF, q <0.05, **Table S3**). Of those, 117 DMPs were hypermethylated and 144 DMPs were hypomethylated in NICM. 192 DMPs (**Table S4**) were common to both ICM and NICM patients (**Fig. 1A**). All of these “common HF DMPs” were either concordantly hypomethylated (n=125) or concordantly hypermethylated (n=67) in ICM and NICM.

**Figure 1.**
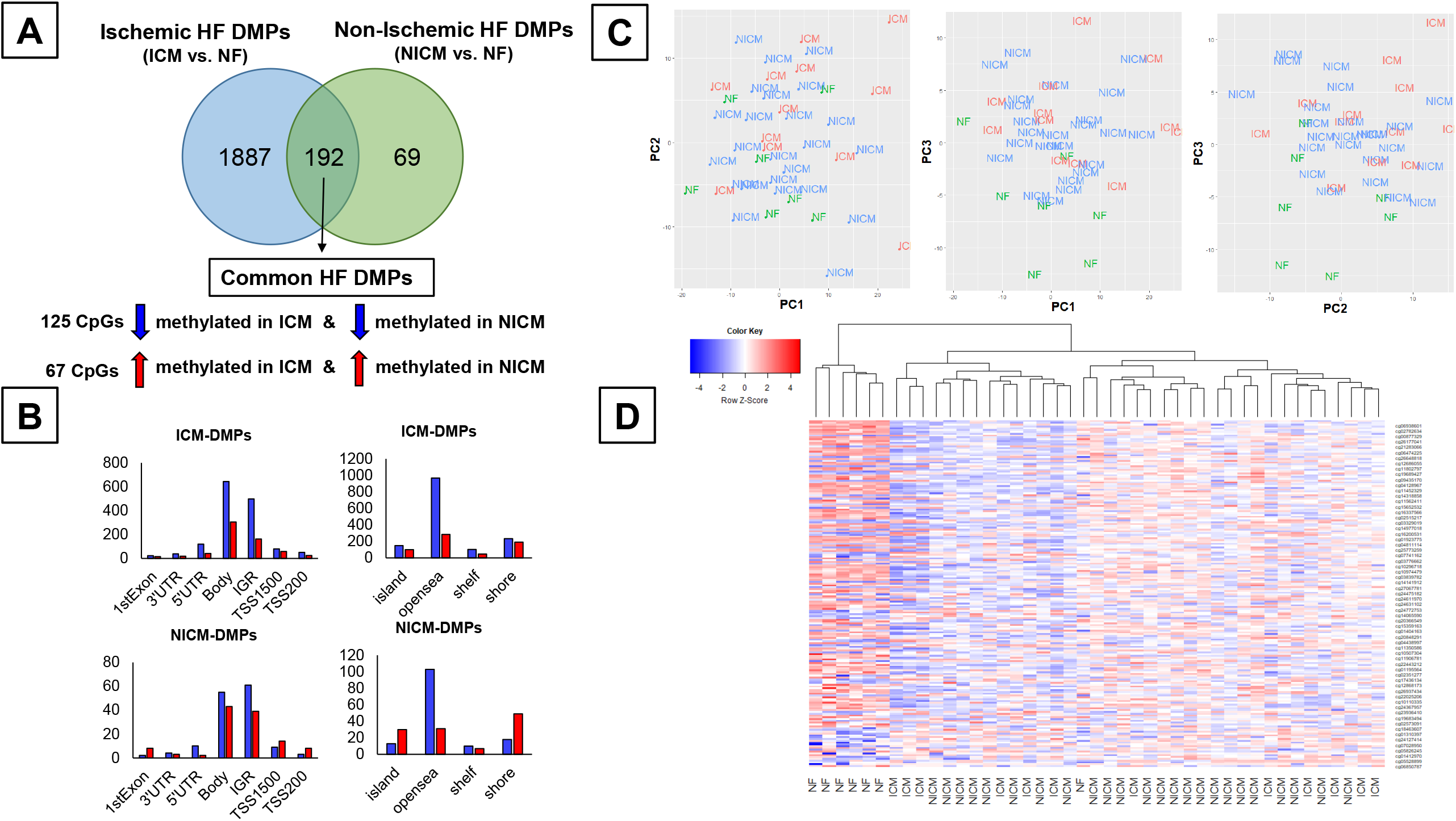
Genome-wide Changes in Myocardial DNA Methylation Based on HF Etiology. A) Venn diagram depicting number of differentially methylated positions (DMPs) in patients with Ischemic (ICM) versus Non-ischemic cardiomyopathy, B) Significant ICM and NICM DMPs by genomic location and feature, C) Principal component analysis of myocardial DNA methylation grouped by etiology of HF (NF: non-failing [green], ICM: ischemic cardiomyopathy [red], and NICM: non-ischemic cardiomyopathy [blue]), D) Heatmap clustering of subjects by methylation profile of 192 Common HF DMPs.

Characterizing the location of DMPs within gene regions, for ICM and NICM similarly, the majority of DMPs were mostly located in gene bodies or intergenic regions (IGR) (**Fig. 1B**). The majority of differentially methylated sites were in open sea regions, with a greater degree of hypomethylation for both ICM and NICM. Hypermethylation in CpG islands and associated shores was more common in the NICM subgroup (**Fig. 1B, bottom panel**). 5.5% of hypomethylated DMPs mapped to protomer regions compared to 9.3% of hypermethylated DMPs in ICM patients. Similarly, 6.3% of hypomethylated DMPs mapped to protomer regions compared to 12.0% of hypermethylated DMPs in NICM patients. Principal component analysis (PCA) of genome-wide methylation levels demonstrates clustering of the non-failing versus failing samples, however, did not clearly separate between ischemic vs non-ischemic etiology (**Fig. 1C**). Heatmap of methylation level Z-scores of 192 common HF DMPs across non-failing healthy controls and HF samples with unbiased hierarchical heatmap clustering separated the non-failing controls from the failing heart DMPs but did not separate ICM from NICM (**Fig. 1D**).

### Minimal Reversibility of Myocardial DNA Methylation with LVAD Support

Patterns of myocardial DNA methylation were analyzed in paired myocardial samples obtained from 8 patients before and after LVAD support. 1075 CpG sites were differentially methylated in the failing myocardium compared to non-failing (pre-LVAD vs NF, q <0.05, **Table S5**). In contrast, only 130 CpG sites were differentially methylated with LVAD support (post-LVAD vs. pre-LVAD, q<0.05, **Table S6**). Only 35 CpG sites (**Table S7**) were common in heart failure and reverse remodeling (**Fig. 2A**), all of which were methylated in opposite directions.

**Figure 2.**
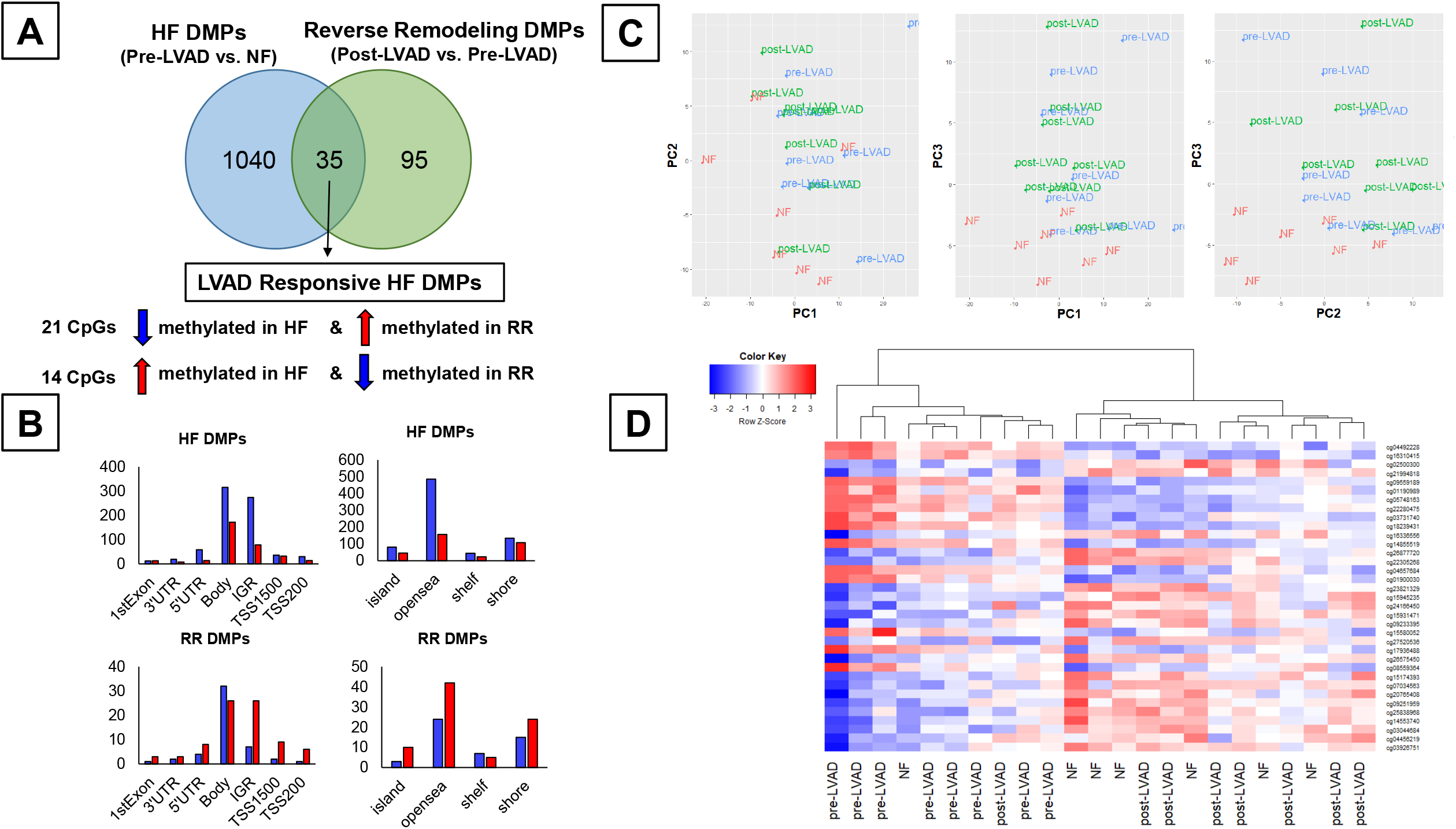
Impact of Mechanical Unloading on myocardial DNA methylation Profile. A) Venn diagram depicting number of differentially methylated positions (DMPs) in Heart Failure (pre-LVAD vs. Non-Failing) versus Reverse Remodeling (post-LVAD vs. pre-LVAD), B) Significant HF and reverse remodeling DMPs by genomic location and feature, C) Principal component analysis of myocardial DNA methylation before and after LVAD support (NF: non-failing [red], pre-LVAD [blue], and post-LVAD [green]), D) Heatmap clustering of subjects by 35 LVAD Responsive HF DMPs.

Characterizing the location of DMPs within gene regions for HF and Reverse Remodeling (RR), the majority of DMPs were mostly located in gene bodies or intergenic regions (IGR) (**Fig. 2B**). The majority of HF DMPs including IGR and transcription start sites (TSSs) were hypomethylated, in contrast the majority of DMPs including IGR and TSSs were hypermethylated in RR. (**Fig. 2B, bottom panel**). PCA of genome-wide methylation levels demonstrated clustering of the non-failing versus pre-LVAD samples, but not pre-vs post-VAD samples, suggesting only minor changes in global DNA methylation with mechanical unloading (**Fig.2C**). Heatmap of methylation level Z-scores of 35 LVAD-responsive HF DMPs across non-failing and paired LVAD samples with unbiased hierarchical clustering demonstrated grouping of post-LVAD samples with 8 out of 9 non-failing control samples (**Fig. 2D**).

### Integrated Analysis of DNA Methylation with Gene Expression in the Failing Human Heart

The classical paradigm of promoter DNA methylation as a transcriptional silencing mechanism has recently been challenged with growing lines of evidence suggesting that DNA hypermethylation could also result in transcriptional activation.^31-33^ Accordingly, we assessed the relationship between differentially methylated CpG sites and changes in gene expression levels using a large publicly available transcriptional dataset obtained from heart failure patients. Out of 192 common HF DMPs, 121 CpG sites map to 93 protein-coding genes (**Fig. 3A**). Out of those 74 genes were expressed in the myocardium (RPKM >1), and 34 of those were differentially expressed in human heart failure. When changes in DNA methylation were correlated with the changes in gene expression, 14 genes were hypomethylated and transcriptionally upregulated, 13 genes were hypomethylated and transcriptionally downregulated, 6 genes were hypermethylated and transcriptionally upregulated, and 1 gene was hypermethylated and transcriptionally downregulated (**Table S8**). Using independent cardiac tissue samples obtained from patients with end-stage ICM and NICM, we validated upregulation of *AKAP13* (fold change [FC] = 2.98), *HTRA1* (FC= 1.52), *EFCAB13* (FC= 2.86), and *FBXO16* (FC= 1.92) as well as downregulation of *TBX3* (FC= 0.39) in the failing human hearts by qPCR analysis, while changes in *RPTOR* and *HDAC9* transcripts did not reach statistical significance (**Fig. 3C**).

**Figure 3.**
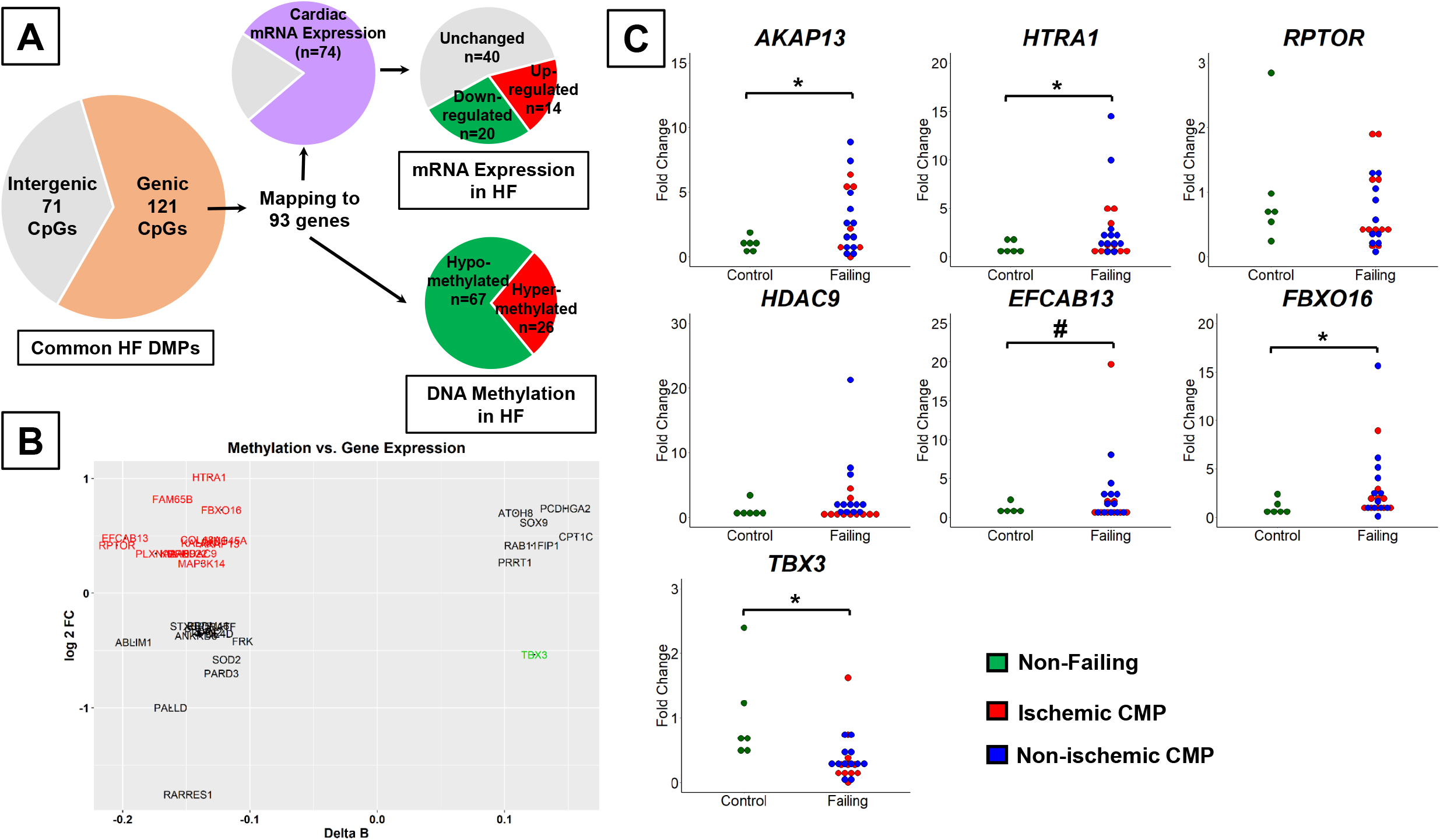
Correlation of DNA Methylation with Gene Expression in Human Heart Failure. A) Differential Methylation and Gene Expression of 192 Common Heart Failure DMPs, B) Methylation vs. Gene Expression Correlation Plot, C) qPCR validation of candidate genes that are epigenetically regulated in independent samples (* p <0.05, # p <0.10).

### Super-enhancer region associated LINC00881 is downregulated in the Failing Human Heart

Common HF and LVAD responsive HF DMPs located in intergenic regions were screened for the presence of overlapping non-coding RNAs using GENCODE and NONCODE datasets. We identified a novel long intergenic non-protein coding RNA (*LINC00881*) located∼9 kb upstream of a significantly hypermethylated CpG site (cg01535205) (**Table S9**). *LINC00881* is highly and exclusively expressed in human myocardium according to GTEx, which made it an interesting candidate for further exploration in the context of epigenetic modification of the failing heart (**Fig. S2**). Interestingly, *LINC00881* and cg01535205 are located within a cardiac super-enhancer region (**Fig. S3**). RNA-seq and CHIP-seq tracks confirms high expression of *LINC00881* in the adult non-failing human heart as well as presence of active chromatin marks in this region including H3K27ac (**Fig. 4A**). qPCR of *LINC00881* in independent HF samples confirmed that this intergenic RNA is downregulated in ICM and NICM compared to myocardium from healthy controls (**Fig. 4B**). *LINC00881* is largely restricted to the nuclear compartment as opposed to cytoplasmic in human induced pluripotent stem-cell derived cardiomyocytes (iPSCs) (**Fig. 4C**). *LINC00881* expression was significantly upregulated during differentiation of human iPSCs, in parallel with upregulation of transcription factors including *GATA4, HAND2*, and *TBX5*, confirming its role as a cardiomyocyte lineage specific super-enhancer lncRNA (**Fig. 4D**).

**Figure 4.**
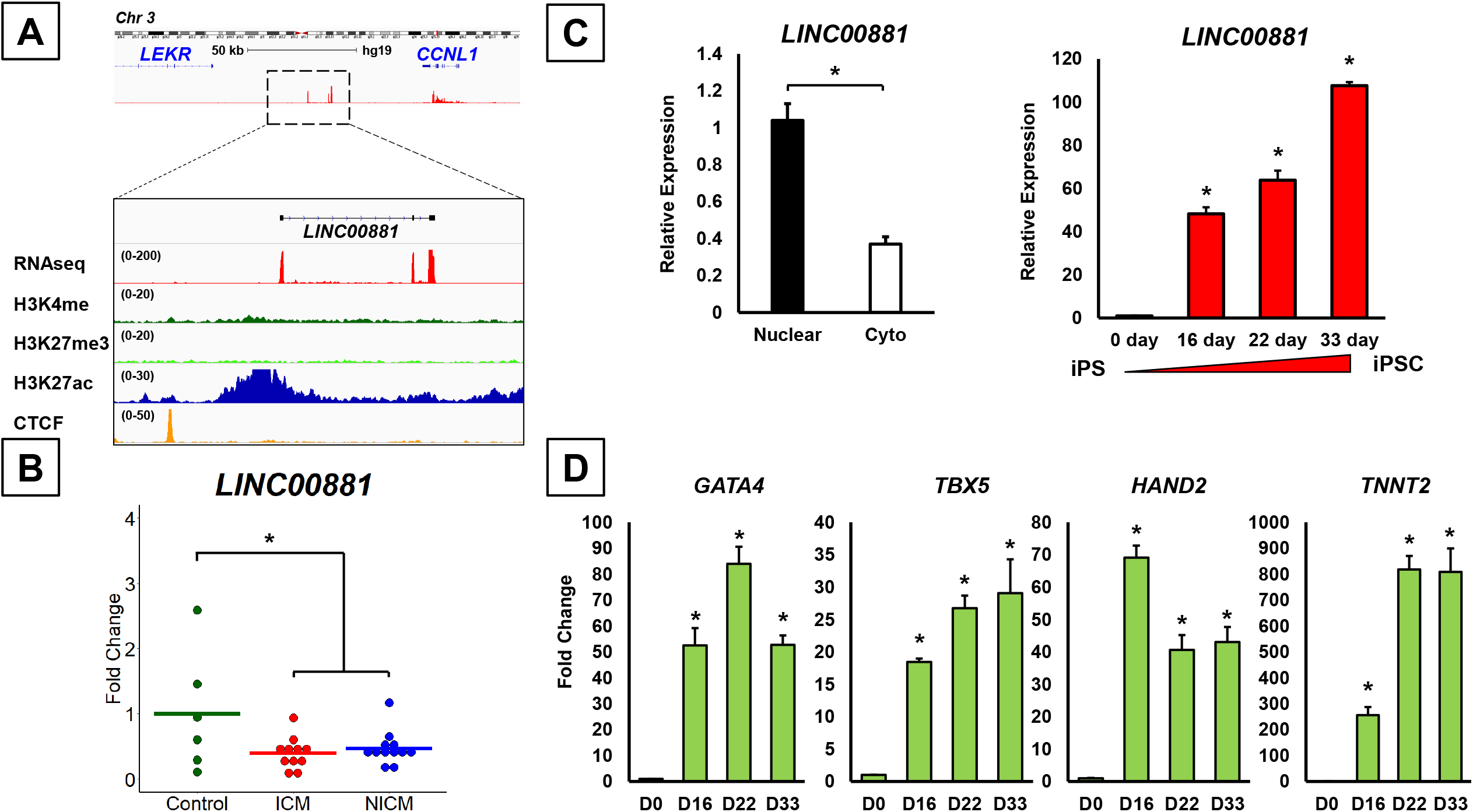
*LINC00881* is a cardiomyocyte lineage-specific super-enhancer lincRNA. A) Position, expression, and epigenetic regulation of the *LINC00881* locus in the non-failing human heart, B) Expression of *LINC00881* in patients with ischemic and non-ischemic (NICM) cardiomyopathy by qPCR (* p <0.05), C) Expression of *LINC00881* in the nuclear versus cytoplasmic fractions (* p <0.05) and in differentiating human iPSCs (* p <0.05 compared to Day 0), (D) Expression of *LINC00881* and cardiac transcription factors during human iPSC differentiation (* p <0.05 compared to Day 0).

To understand the mechanistic basis of *LINC00881* dysregulation in human heart failure, we used plasmid-mediated overexpression and GapmeR-based knockdown of *LINC00881* in the beating human iPSCs (**Fig. S4)**. RNA-sequencing identified 1545 genes that were differentially expressed with *LINC00881* overexpression (**Table S10**) and 2268 genes that were differentially expressed with *LINC00881* knockdown (p-value cut-off <0.05) (**Table S11**) (**Fig. S5**). Among 199 common genes that were differentially regulated in both *LINC00881* overexpression and *LINC00881* knockdown models, 174 (87.4%) were regulated in opposite directions including 73 genes that are positively regulated and 101 genes that are negatively regulated by *LINC00881* (**Table S12, Fig. 5A**). Gene Ontology analysis of transcripts that are positively regulated by *LINC00881* demonstrated sarcomere organization (*MYH6, MYOM2, LDB3, LFOD3, OSBCN*) and calcium ion transport as the top positively regulated pathways (*CACNA1C, CACNA1D, RYR2, CAMK2A, ANXA6)* (**Fig 5B)**. Gene Ontology analysis of transcripts that are negatively regulated by *LINC00881* showed regulation of transcription from RNA II polymerase promoter (*CEBPG, KLF5, KLF10, SMAD5, ATF1, ATF3*, and *ANKRD1)* and regulation of apoptotic process (*BCLAF1, DNAJA1, GNA13, MCL1, ANRKRD1, PHLDA1, SIRT1*) as the top negatively regulated pathways. *LINC00881* regulation of sarcomere and calcium channel genes were validated in human iPSCs by qPCR with or without *LINC00881* knockdown (**Fig. 5C**). To determine whether *LINC00881* regulation of sarcomere genes and calcium transport genes have functional relevance in the heart, we measured calcium transients in beating human iPSCs treated with GapmeRs targeting *LINC00881* versus scrambled control oligonucleotide. *LINC00881* knockdown resulted in significant reductions in the peak calcium amplitude in human iPSCs (**Fig. 5D**).

**Figure 5.**
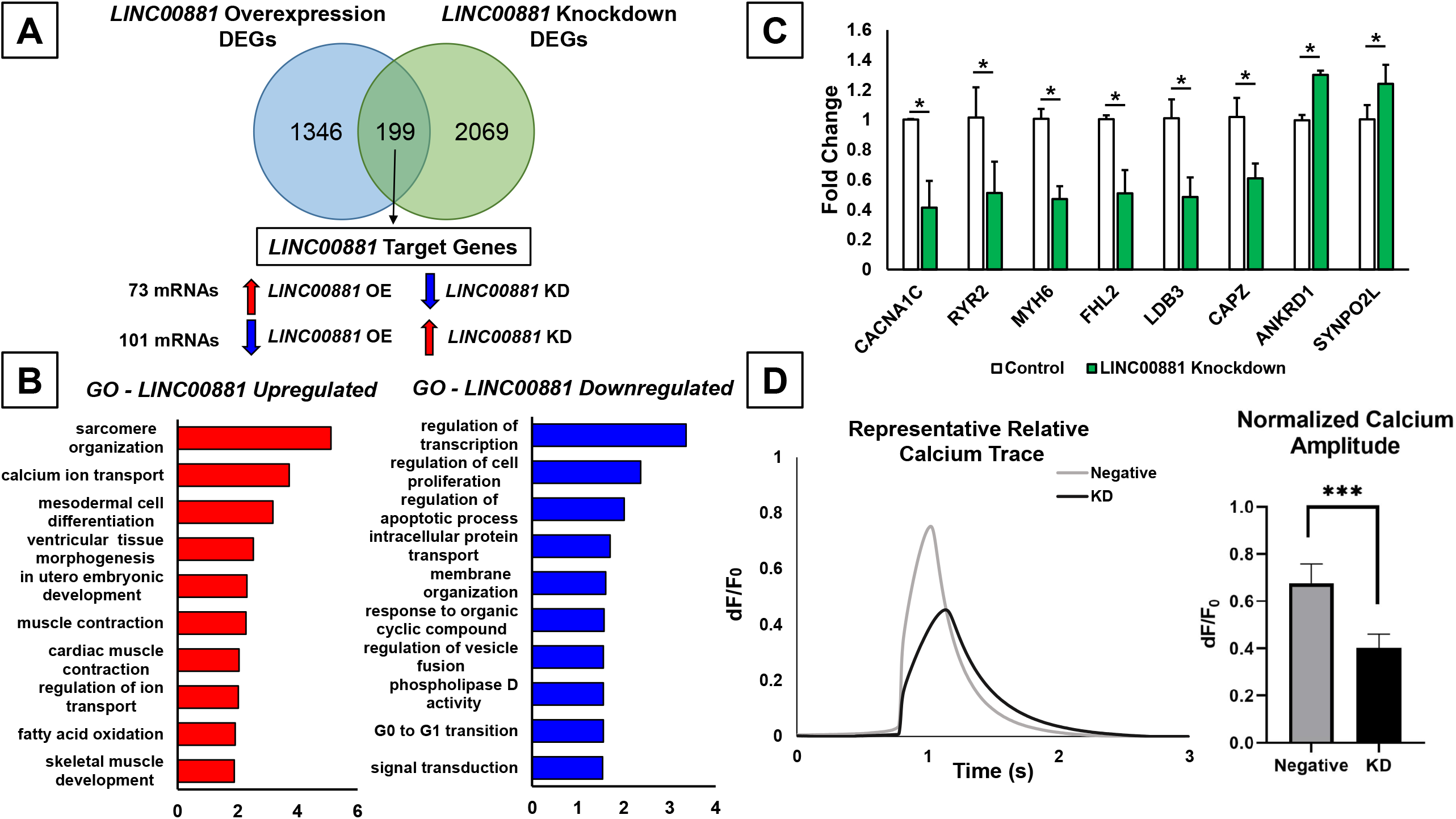
*LINC00881* is an essential regulator of cardiomyocyte calcium cycling. A) Venn-diagram depicting number of differentially expressed genes in human iPS derived cardiomyocytes with *LINC00881* plasmid-based overexpression or GapmeR-mediated knockdown, B) Gene Ontology analysis of gene targets that are positively or negatively regulated by *LINC00881* for Biological Process, C) qPCR validation of LINC00881 target genes (* p <0.05), D) Representative relative calcium traces in beating human iPSCs with *LINC00881* versus scrambled oligonucleotide knockdown with averaged normalized calcium amplitude.

## Discussion

The present study utilized bead array technology for high-density genome-wide mapping of DNA methylation in the failing human heart before and after LVAD support. Our analysis identified myocardial DNA methylation patterns that are associated with reciprocal regulation of gene expression in patients with ICM and NICM. In addition to providing the most comprehensive mapping of etiology-specific changes in myocardial DNA methylation to date, we show for the first time that mechanical unloading with LVAD is associated with an incomplete normalization of the myocardial DNA methylation profile, suggesting that HF-related epigenetic alterations could be persistent. Moreover, our analysis identified a novel cardiac-specific super-enhancer lncRNA gene (*LINC00881*), which is hypermethylated and downregulated in the failing human heart, as an essential regulator of cardiomyocyte calcium cycling and contractility (**Fig 6**).

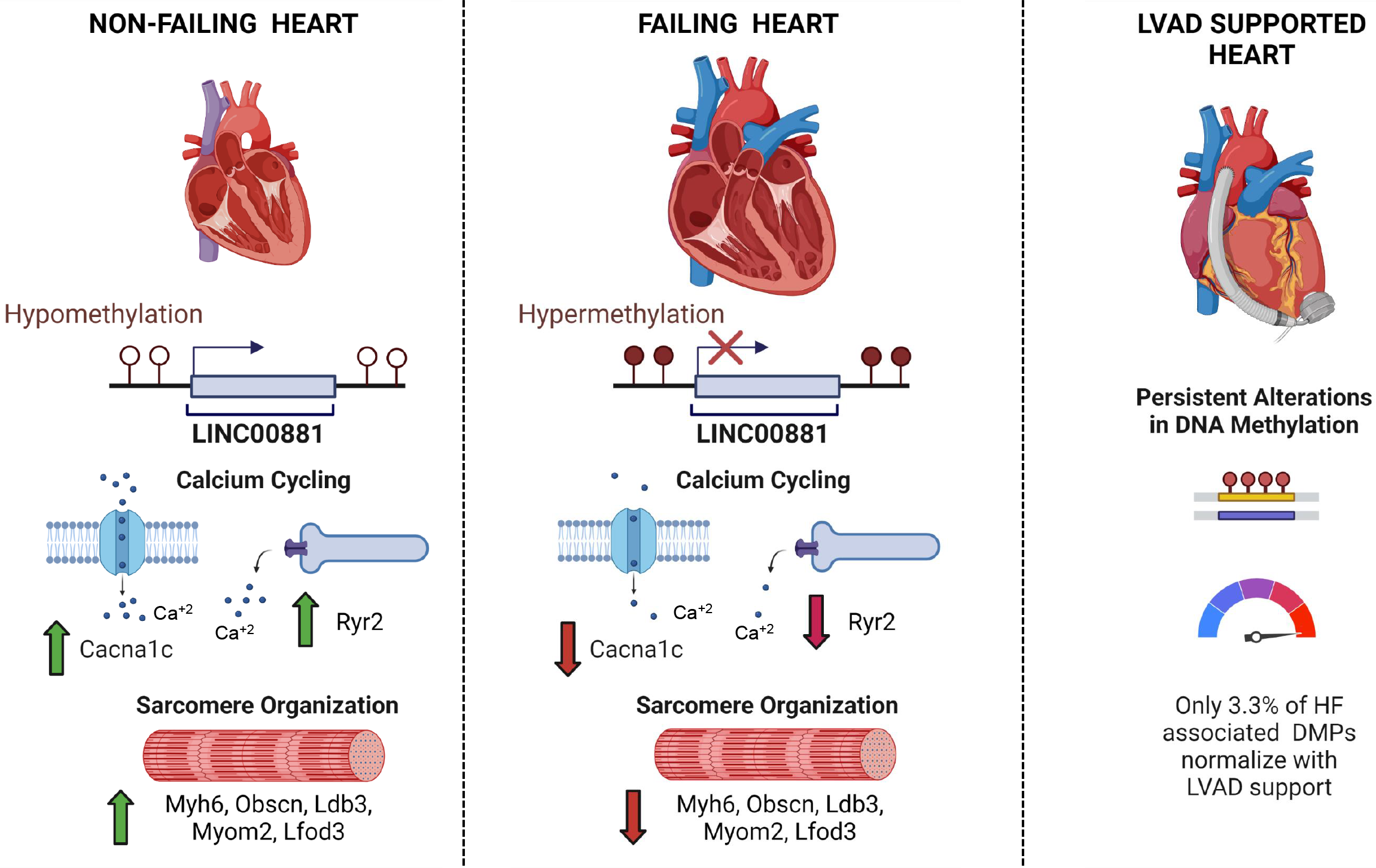

### DNA Methylation in Human Heart Failure

Consistent with previous reports, we found that the majority of differentially methylated positions in end-stage heart failure had reduction in DNA methylation levels, suggesting a global hypomethylation similar to what has been demonstrated in cancer biology.^17, 18, 20, 34^ Movassagh et al. demonstrated that HF-related differential methylation in CpG islands was predominantly located in gene promoters and gene bodies, but not in intergenic or 3’UTRs.^17^ Our analysis expands on this observation and demonstrates a large number of intergenic positions that are also differentially methylated, including a subset mapping to non-coding genome. Our etiology-specific analysis identified 192 DMPs that were common to patients with ICM (2079 DMPs) and NICM (261 DMPs), similar to a recent study by Glezeva et al. which identified 13 common HF DMPs from patients with ICM (51 DMPs) and NICM (118 DMPs).^20^ Overlapping DNA methylation profiles in patients with ICM and NICM support a convergent “final common pathway” of gene expression that are found in all forms of heart failure irrespective of the inciting event. Our analysis showed that hypomethylated CpG sites, particularly in ICM, were more likely to be within gene body or intergenic regions as opposed to transcription start sites, which is consistent with a prior study by Pepin et al. demonstrating a relative hypermethylation of promoter-associated CGIs in ICM patients.^35^ 25 out of 93 common HF DMPs mapping to a protein-coding gene in current study were also found to be differentially methylated in at least one previous study validating our analytical approach.^18, 22, 23^

### Effect of Mechanical Unloading with LVAD on Myocardial DNA Methylation Patterns

Mechanical unloading with LVAD is associated with favorable changes in the biology of the failing cardiomyocyte including regression of myocyte hypertrophy, improvement in excitation-contraction coupling, and down-regulation of the fetal gene expression, also termed as reverse remodeling, which results in normalization of cardiac function in a small number of patients allowing for LVAD explantation.^36-39^ Transcriptional studies of paired myocardial samples obtained from patients with end-stage heart failure before and after LVAD support demonstrate that only <5% of HF-related transcripts normalize with LVAD support.^7, 9^ Consistent with these observations, we found that only 3.2% of HF-related DMPs were reverse methylated with LVAD support, which may in part explain persistent transcriptional dysregulation and the low incidence of myocardial recovery in patients supported by LVAD. Since DNA methylation is a reversible phenomenon, these findings raise the possibility that epigenetic modulation may be necessary to normalize the HF transcriptome and achieve myocardial recovery. In support of this hypothesis, inhibition of DNA methylation using 5-aza-2′-deoxycytidine rescued a HF phenotype in a rat model of norepinephrine induced cardiac hypertrophy.^40^ Mice with cardiomyocyte-specific deletion of DNMT3b develop cardiomyopathy with sarcomeric disarray and interstitial fibrosis.^41^ Similarly, differences in myocardial DNA methylation among mouse strains has been shown to determine susceptibility to cardiac hypertrophy following isoproterenol treatment.^42^ Taken together, these findings suggest that epigenetic reprogramming may have therapeutic relevance in heart failure and additional research is necessary to elucidate the mechanistic basis of this approach.

### Common Heart Failure DMPs with Reciprocal Changes in Gene Expression

Our analysis identified a subset of differentially methylated protein-coding genes, which were transcriptionally regulated in the opposite direction of methylation in HF. Among upregulated genes were *HTRA1, FAM65B, UNC45A, KALRN, AKAP13, RPTOR*, and *HDAC9*, which have been previously implicated in maladaptive hypertrophy. *HTRA1* has been identified as part of a candidate gene signature correlated with cardiomyopathies in a gene correlation network analysis model and its mRNA expression is upregulated 6.9 fold in DCM.^43, 44^ *PINK1* dependent phosphorylation of *FAM65B* attenuates ischemia reperfusion injury by suppressing autophagy.^45^ The *UNC45A* gene has been characterized as potential de novo mosaic variant in sporadic cardiomyopathy.^46^ *KLRN* is a Rho Guanine Nucleotide Exchange Factor (*GEF*), which is downregulated in ICM and NICM hearts.^47^ A-Kinase Anchoring Protein 13 (*AKAP13*) promotes downstream hypertrophic gene expression, mediated at least in part via *HDAC5* phosphorylation and *MEF2*-mediated transcription in TAC model of cardiac hypertrophy.^48^ Genetic deficiency of *RPTOR*, regulatory associated protein of mTOR Complex 1, leads to reduction of mTORC1 activity and dilated cardiomyopathy in mice.^49^ Mice lacking *HDAC9* are sensitized to hypertrophic signals and exhibit stress-dependent cardiomegaly.^50^ *miR-21* levels are increased selectively in fibroblasts of the failing heart.^51^ In vivo silencing of *miR-21* has been shown to attenuate cardiac dysfunction.^51, 52^

*TBX3* was the only protein-coding gene common to ICM and NICM with DNA hypermethylation and transcriptional downregulation in our analysis. *TBX3* is located within the 12q24.21 locus, which harbors several other genes that have previously been linked to cardiomyopathies. Meder et al. identified this gene locus to be differentially methylated in NICM patients, validating our observation.^23^ Genetic variation in *TBX3* was associated with LV mass in health Japanese population, highlighting potential implication in cardiac hypertrophy, however precise mechanisms warrants further investigation.^53^ In addition, we identified and validated several genes that were hypomethylated and upregulated in the failing human heart with previously unknown link to HF including *FXBO16, EFCAB13, COL18A1, PLXNA2, BRE, KIAA0922*, and *MAP3K14*. Additional research is warranted to investigate the function of these genes in the HF pathophysiology.

### Role of LINC00881 in Human Heart Failure

Our genome-wide screening identified a novel lincRNA with epigenetic and transcriptional dysregulation in human heart failure. *LINC00881* is transcribed from a cardiac specific super-enhancer region with abundant expression levels in the adult human heart. We show that this super-enhancer region is hypermethylated which is associated with down-regulation of *LINC00881* gene expression in patients with ICM and NICM. To date, very little has been known regarding the role and function of *LINC00881* except that it is expressed in cardiomyocytes and regulated by a GATA-4 responsive super-enhancer element.^54, 55^ *LINC00881* (NR_034008) was listed among significantly down-regulated transcripts in the failing human heart in two independent transcriptomic publications validating our observation.^5, 56^ Our in-vitro work in beating human iPSCs expands on the role of LINC00881 in human HF and suggest that it is an essential regulator of cardiomyocyte calcium cycling and contractility by regulating expression of several key calcium channel and sarcomeric genes including *CACNA1C, RYR2*, and *MYH6*.

Potential limitations of the study include unknown confounders that may influence DNA methylation patterns in the human myocardium. Cardiac tissue is composed of multitude of cell types including cardiomyocytes, fibroblasts, endothelial cells, and immune cells, which likely have distinct methylation profiles. Since our study utilized myocardial DNA for profiling, it does not address cell-specific alterations in the DNA methylation profile.^19^ We used bead array technology for methylome profiling and were restricted to genomic sites that were included in the platform inferring a selection bias. Epigenetic modifications other than DNA methylation such as histone modifications, which may have an impact on transcriptional dysregulation in HF, were not assessed in this study.

In conclusion, heart failure is associated with common and distinct alterations in DNA methylation in ischemic versus non-ischemic failing myocardium. Mechanical unloading with LVAD fails to normalize the majority of HF-related DNA methylation, which remain persistently dysregulated. Differential DNA methylation regulates expression of both protein-coding and non-coding transcripts in the failing human heart with previously uncharacterized functions. Among these, *LINC00881* is a cardiac super-enhancer lncRNA, which is an essential regulator of cardiomyocyte calcium cycling. These findings suggest that epigenetic targeted therapies could be necessary to normalize the dysregulated transcriptome in the failing human myocardium and help achieve sustained clinical recovery from heart failure.

## Acknowledgements

Authors thank Dr. Benjamin Tycko and Dr. Catherine Do from Columbia University Epigenetics Core for their assistance with DNA methylation data analysis. X.L. performed majority of experiments with help from B.L, T.R.N, R.Z., A.G.F., and R.L. V.K.T. and X.L. designed the experiments. V.K.T., X.L., and P.J.K. wrote the manuscript with input from co-authors. P.C.C., N.U., S.O.M., M.P.R, and G.V.N. provided conceptual advice. V.K.T. coordinated and oversaw the whole project.

## Material Availability

All unique reagents generated in this study are available from the lead contact with a completed materials transfer agreement

