## Supplemental Materials for "Genome-Wide DNA Methylation Profiling of the Failing Human Heart with Mechanical Unloading Identifies *LINC00881* as an Essential Regulator of Calcium Handling in the Cardiomyocyte"

### CHAMP DNA Methylation
### Data Analysis Pipeline

#### Data Processing and Annotation

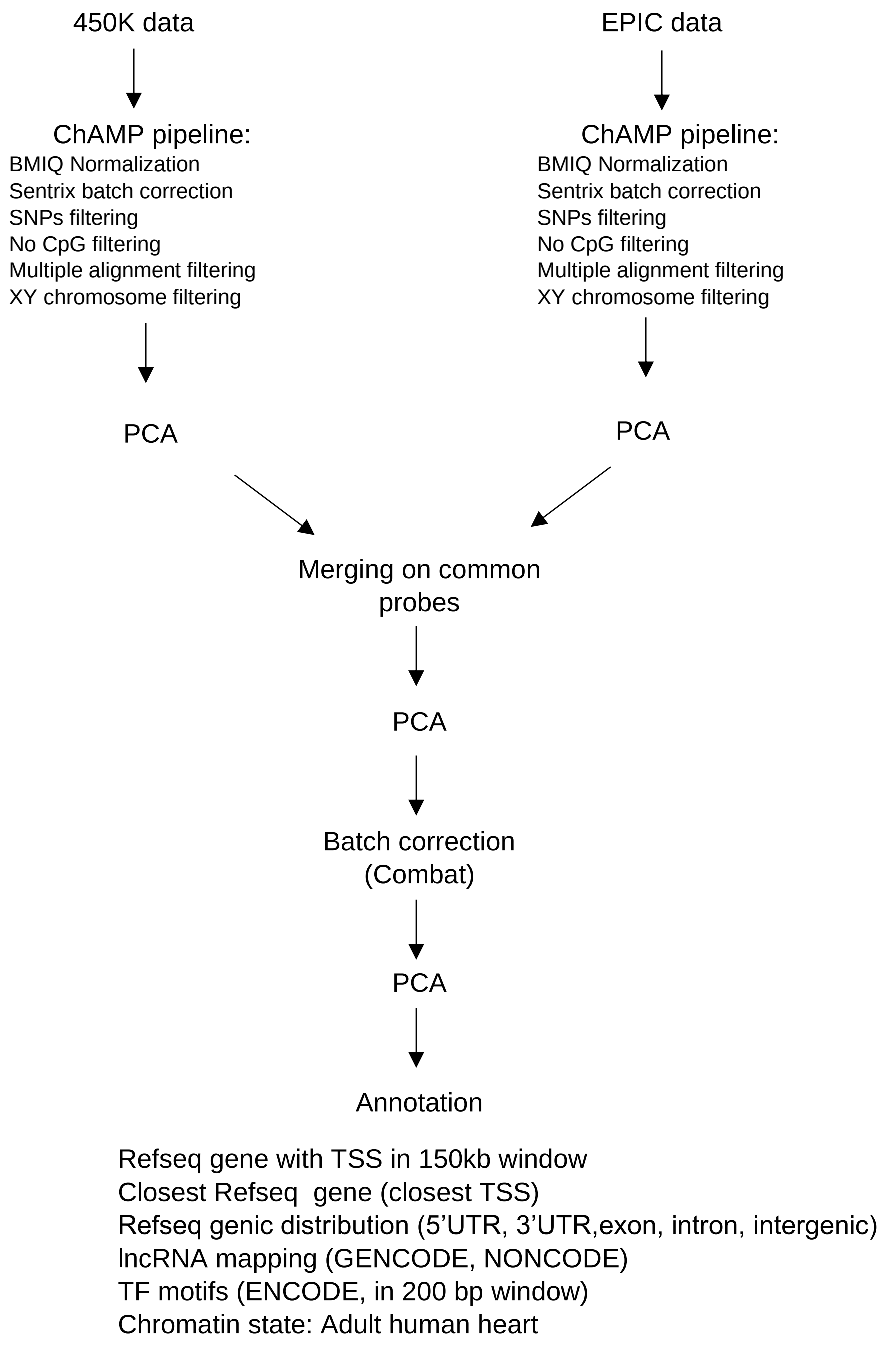


**Supplemental Figure 1A.** DNA Methylation Analysis pipeline using Bioconductor CHAMP Package from Illumina 450K and EPIC bead-array chip data.

#### Analysis 1: Etiology Specific HF DMPs

**DMP Associated with Heart Failure**

Annotated and filtered merged dataset

Case control:

Ischemic HF vs Non-Failing

DM: absolute Δ fract. Methyl>0.1

q-value <0.05

ISCHEMIC DMPs

Case control:

Non-Ischemic HF vs Non-Failing

DM: absolute Δ fract. Methyl>0.1

q-value <0.05

NON-ISCHEMIC DMPs

#### Analysis 2: LVAD Responsive HF DMPs

**DMP Associated with Reverse Remodeling on LVAD**

Annotated and filtered merged dataset

Case control:

Pre-LVAD vs Non-Failing

DM: absolute Δ fract. Methyl>0.1

q-value <0.05

HEART FAILURE DMPs

Paired Analysis:

Pre-LVAD vs Post-LVAD

DM: absolute Δ fract. Methyl>0.1

q-value <0.05

REVERSE REMODELING DMPs

**Supplemental Figure 1B.** Approach for human heart differential DNA Methylation analysis focusing on HF etiology specific (Analysis #1) and LVAD responsive (Analysis #2) differentially methylated positions.

**A**

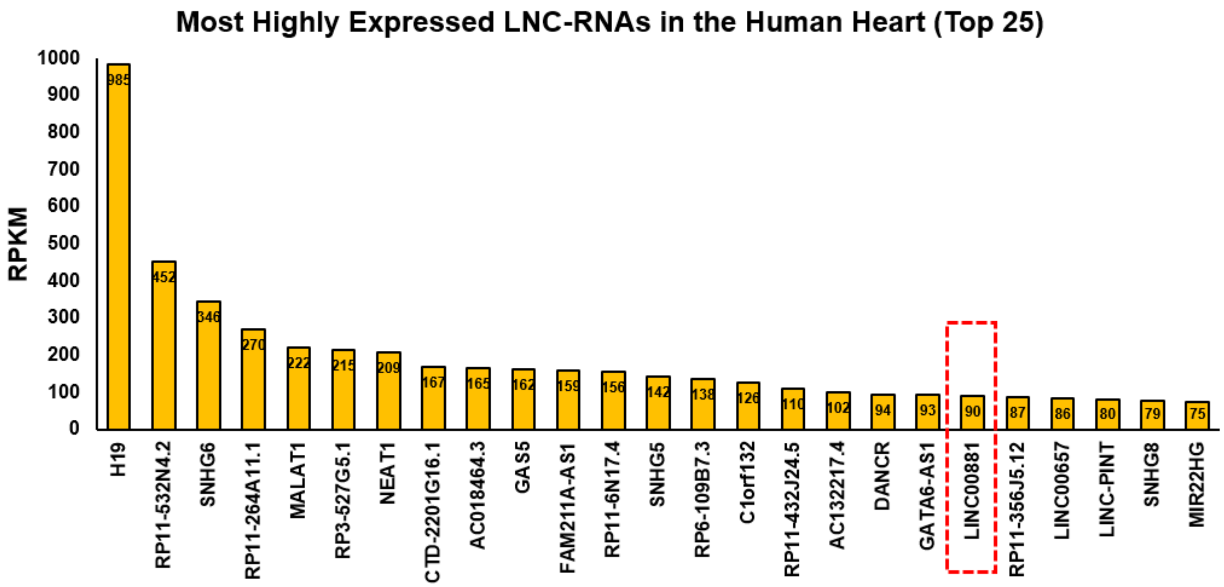


**B**

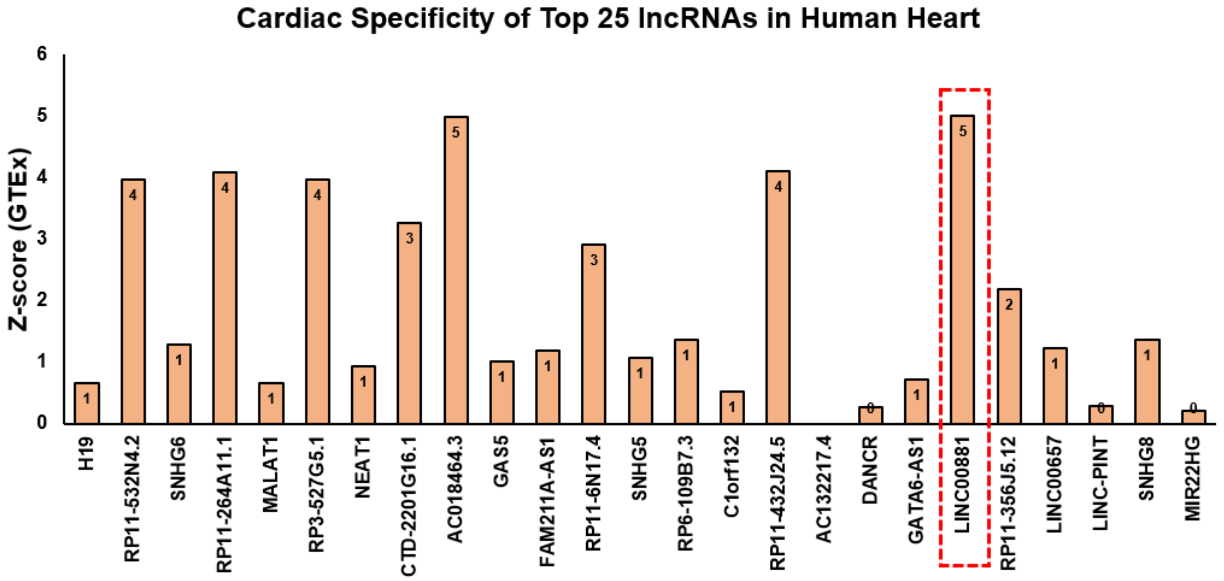


**Supplemental Figure 2. Top Lnc-RNAs in the Human Heart. (A)** Most highly expressed lnc-RNAs in the human heart in the descending order of RPKM (Top 25) using RNA-seq data obtained from GEO accession number 116250 **(B)** Cardiac-specificity of Top 25 cardiac lncRNAs in the GTEx database

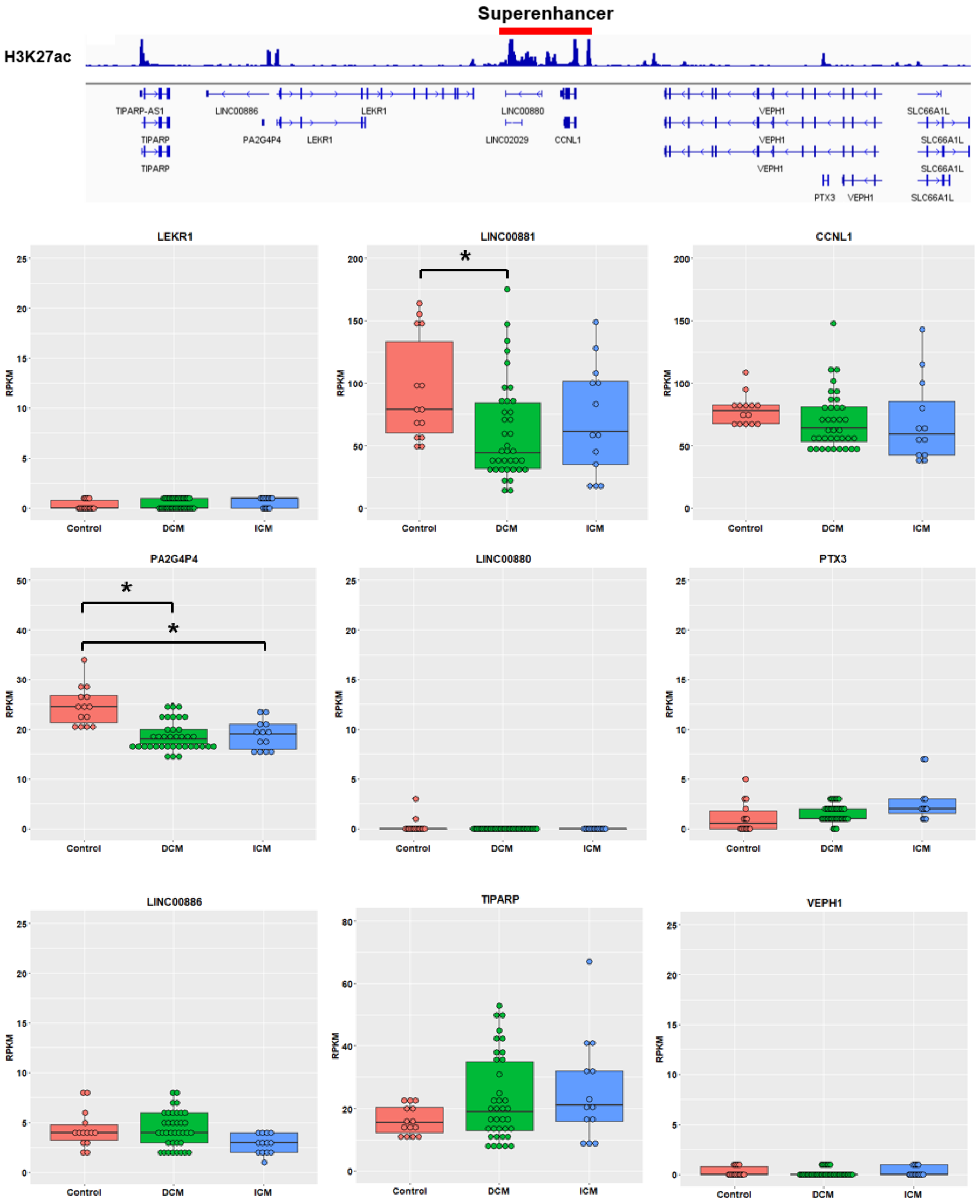


**Supplemental Figure 3. LINC00881 cardiac super-enhancer region**. Expression levels of LINC00881 and neighboring coding and non-coding transcripts in the non-failing (Control-red), non-ischemic (DCM-green), and ischemic (ICM-blue) human heart failure. RNA-seq data obtained from GEO accession number 116250. * FDR value <0.05

A

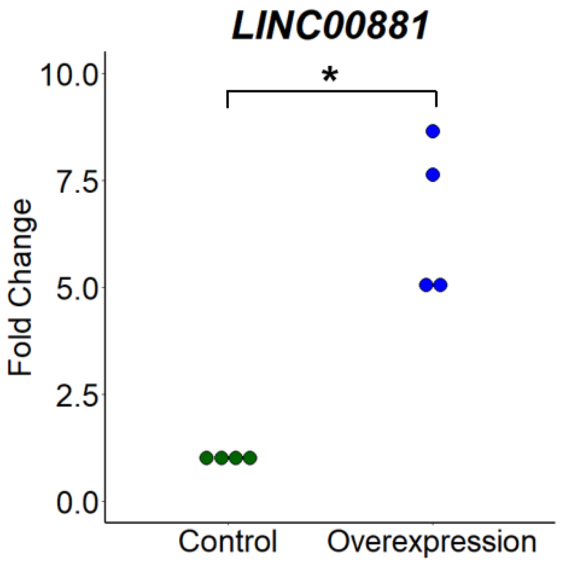


B

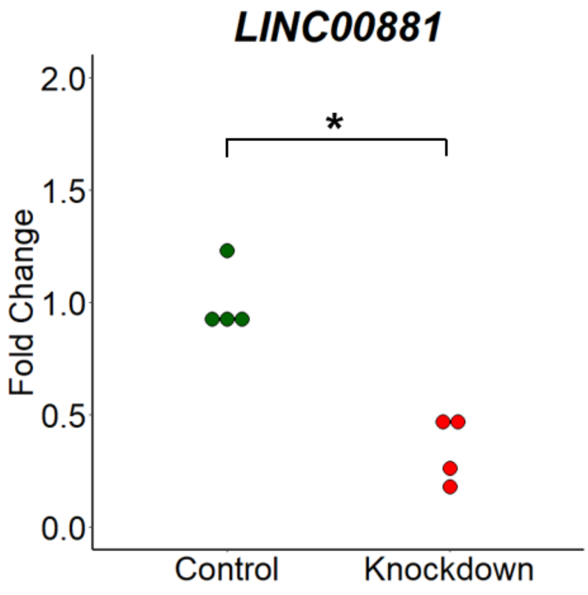


**Supplemental Figure 4. LINC00881 overexpression and knockdown in the beating human iPS cell derived cardiomyocytes**. (A) Plasmid-based overexpression of *LINC00881* validated by qPCR (B) GapmeR-mediated knockdown of *LINC00881* validated by qPCR * p-value <0.05

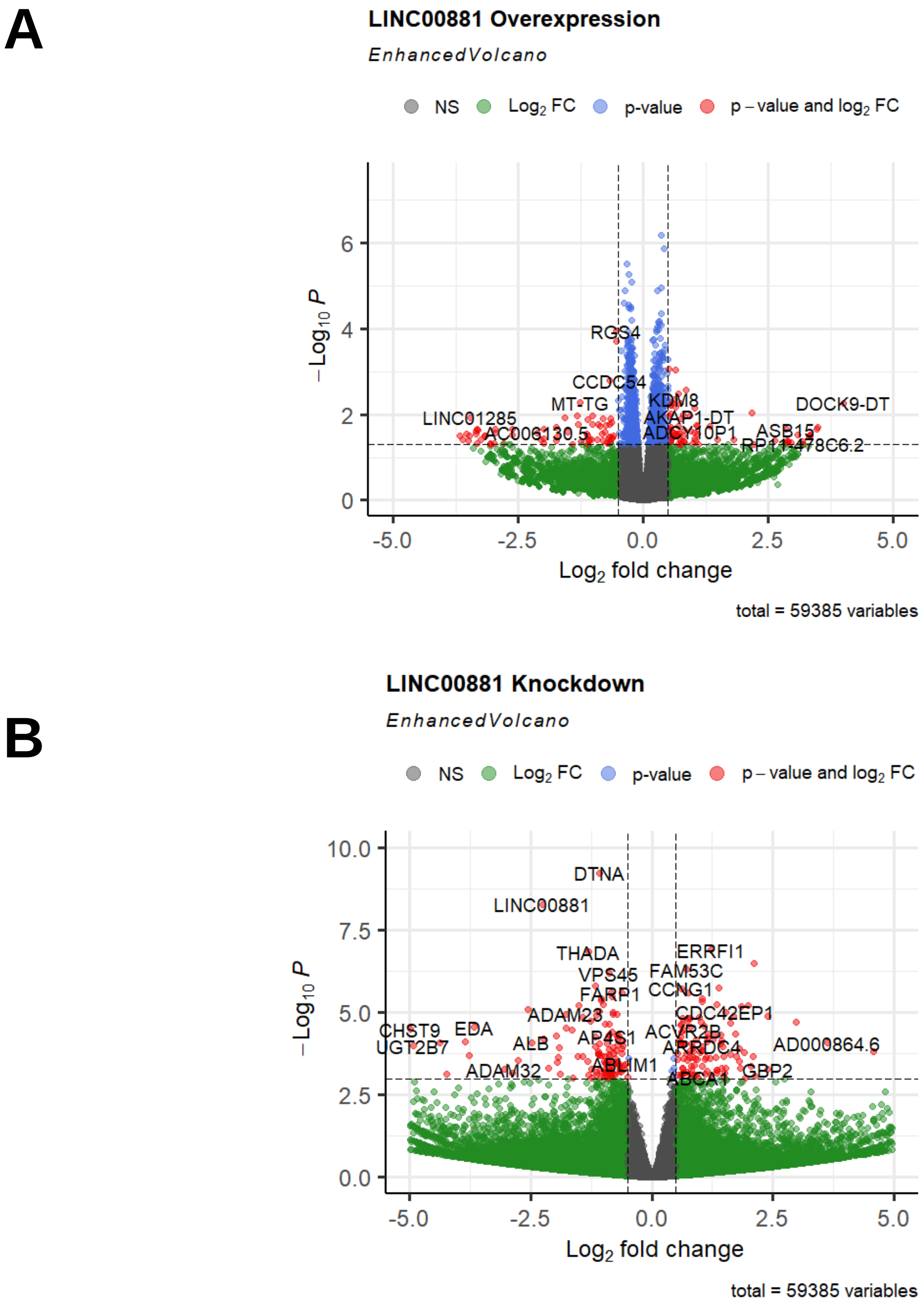


**Supplemental Figure 5. RNA-seq in the beating human iPS cell derived cardiomyocytes**.
(A) Volcano plot of differentially expressed transcripts with *LINC00881* plasmid overexpression
(B) Volcano plot of differentially expressed transcripts with *LINC00881* GapmeR knockdown

**Supplemental Table 1A.** Clinical Characteristics of Study Subjects with Heart Failure (n=36)

| Variables | Full Cohort (n=36) | Ischemic HF (n=12) | Non-Ischemic HF (n=24) | p-value |
|---|---|---|---|---|
| **Age** | 56.7 ± 13.3 | 64.7 ± 6.4 | 52.65 ± 14.2 | **0.008** |
| **Gender (M)** | 69% (25) | 67% (8) | 71% (17) | 0.999 |
| **Ethnicity (Non-white)** | 47% (17) | 17% (2) | 62% (15) | **0.014** |
| **LVAD Type** | | | | |
| Heartmate II | 83% (30) | 58% (7) | 96% (23) | **0.010** |
| Heartware LVAD | 17% (6) | 42% (5) | 4% (1) | |
| **Destination Therapy** | 44% | 42% | 46% | |
| **LVEF (%)** | 15 ± 4 | 17 ± 3 | 14 ± 3 | **0.023** |
| **LVEDD (cm)** | 7.1 ± 1 | 6.7 ± 1.1 | 7.3 ± 1 | 0.121 |
| **Serum Creatinine** | 1.41 ± 0.58 | 1.26 ± 0.32 | 1.49 ± 0.67 | 0.283 |
| **Serum BNP** | 1402 ± 1564 | 1061 ± 918 | 1597 ± 1828 | 0.353 |
| **Medication Use** | | | | |
| Beta-blocker | 83% (30) | 92% (11) | 80% (19) | 0.639 |
| NHB | 64% (23) | 58% (7) | 66% (16) | 0.719 |
| Diuretics | 47% (17) | 58% (7) | 42% (10) | 0.483 |
| Antiarrhythmics | 53% (19) | 58% (7) | 50% (12) | 0.732 |
| **Duration of support** | 577 ± 424 | 652 ± 522 | 627 ± 487 | 0.672 |
| **LVAD Outcome** | | | | |
| Death | 22% (8) | 25% (4) | 17% (4) | 0.397 |
| Transplant | 61% (22) | 50% (6) | 67% (16) | 0.471 |
| Ongoing | 9% (3) | 0% | 13% (3) | N/A |

**Supplemental Table 1B.** Echocardiographic and Laboratory Markers of 8 patients with paired pre- and post- LVAD cardiac tissue

| Variables | Pre-LVAD (n= 8) | Post-LVAD (n=8) | p-value |
|---|---|---|---|
| **LVEF (%)** | 16 ± 3 | 26 ± 14 | 0.072 |
| **LVEDD (cm)** | 7.3 ± 0.8 | 6.3 ± 0.96 | **0.05** |
| **Serum Creatinine** | 1.74 ± 1 | 1.86 ± 1.9 | 0.88 |
| **Serum BNP** | 1654 ± 1677 | 450 ± 619 | 0.08 |

**Supplemental Table 2.** Differentially methylated positions in Ischemic HF vs. Non-Failing (adjusted p-value <0.05, abs (delta beta) > 15%)

| CpG Site | log FC | Adj p-val | ICM | NF | Delta B | Gene | Feature | CGI |
|---|---|---|---|---|---|---|---|---|
| cg07167872 | -0.34 | 0.0076 | 0.58 | 0.24 | -0.34 | PM20D1 | TSS200 | shore |
| cg04118610 | -0.33 | 0.0034 | 0.76 | 0.43 | -0.33 | LPHN3 | Body | opensea |
| cg05528899 | -0.33 | 0.0432 | 0.67 | 0.35 | -0.33 | | IGR | island |
| cg14159672 | -0.32 | 0.0182 | 0.55 | 0.23 | -0.32 | PM20D1 | 1stExon | island |
| cg11965913 | -0.30 | 0.0178 | 0.39 | 0.09 | -0.30 | PM20D1 | TSS200 | shore |
| cg04811114 | -0.26 | 0.0001 | 0.47 | 0.21 | -0.26 | LGR6 | TSS200 | opensea |
| cg14893161 | -0.26 | 0.0206 | 0.39 | 0.14 | -0.26 | PM20D1 | 5'UTR | shore |
| cg26536949 | -0.25 | 0.0442 | 0.76 | 0.51 | -0.25 | | IGR | island |
| cg17178900 | -0.24 | 0.0262 | 0.63 | 0.39 | -0.24 | PM20D1 | Body | island |
| cg24503407 | -0.24 | 0.0182 | 0.64 | 0.40 | -0.24 | PM20D1 | TSS1500 | shore |
| cg12777520 | -0.20 | 0.0002 | 0.55 | 0.35 | -0.20 | LMX1B | Body | island |
| cg13283845 | -0.20 | 0.0266 | 0.77 | 0.57 | -0.20 | | IGR | shore |
| cg04160030 | -0.19 | 0.0002 | 0.63 | 0.43 | -0.19 | FUCA1 | TSS1500 | shore |
| cg24122364 | -0.19 | 0.0038 | 0.59 | 0.40 | -0.19 | DOCK9 | Body | opensea |
| cg14204784 | -0.18 | 0.0002 | 0.53 | 0.35 | -0.18 | LMX1B | Body | island |
| cg12563372 | -0.18 | 0.0001 | 0.50 | 0.32 | -0.18 | | IGR | shore |
| cg02928365 | -0.18 | 0.0000 | 0.44 | 0.26 | -0.18 | HLX | Body | shore |
| cg12930727 | -0.18 | 0.0023 | 0.47 | 0.30 | -0.18 | HYAL1 | TSS1500 | opensea |
| cg24382823 | -0.18 | 0.0005 | 0.44 | 0.26 | -0.18 | LMX1B | Body | island |
| cg17436134 | -0.17 | 0.0034 | 0.47 | 0.30 | -0.17 | | IGR | shore |
| cg12583076 | -0.17 | 0.0060 | 0.56 | 0.39 | -0.17 | RASSF3 | Body | opensea |
| cg10070864 | -0.17 | 0.0009 | 0.72 | 0.55 | -0.17 | | IGR | shelf |
| cg23066280 | -0.17 | 0.0007 | 0.54 | 0.37 | -0.17 | PTPRN2 | Body | opensea |
| cg14318858 | -0.17 | 0.0000 | 0.77 | 0.60 | -0.17 | CPT1C | Body | island |
| cg19657945 | -0.17 | 0.0452 | 0.92 | 0.75 | -0.17 | | IGR | shore |
| cg18918831 | -0.17 | 0.0418 | 0.66 | 0.50 | -0.17 | MUC4 | Body | island |
| cg11679455 | -0.16 | 0.0002 | 0.68 | 0.51 | -0.16 | GATA3 | Body | island |
| cg10902396 | -0.16 | 0.0008 | 0.52 | 0.36 | -0.16 | CHN2 | Body | opensea |
| cg02455346 | -0.16 | 0.0008 | 0.68 | 0.52 | -0.16 | HLX | Body | shore |
| cg23936410 | -0.16 | 0.0072 | 0.43 | 0.27 | -0.16 | | IGR | opensea |
| cg24838345 | -0.16 | 0.0020 | 0.69 | 0.53 | -0.16 | MTSS1 | Body | shelf |
| cg06410057 | -0.16 | 0.0014 | 0.35 | 0.19 | -0.16 | | IGR | shore |
| cg22077361 | -0.16 | 0.0064 | 0.42 | 0.27 | -0.16 | FUCA1 | TSS1500 | shore |
| cg16200531 | -0.16 | 0.0001 | 0.61 | 0.45 | -0.16 | | IGR | opensea |
| cg01525538 | -0.16 | 0.0003 | 0.45 | 0.30 | -0.16 | DUSP5P | Body | island |
| cg01107874 | -0.16 | 0.0003 | 0.73 | 0.58 | -0.16 | C10orf41 | Body | island |
| cg13752114 | -0.16 | 0.0271 | 0.84 | 0.68 | -0.16 | MUC4 | Body | island |
| cg03329019 | -0.15 | 0.0001 | 0.62 | 0.47 | -0.15 | | IGR | shore |
| cg10653297 | -0.15 | 0.0094 | 0.68 | 0.53 | -0.15 | LOC285033 | TSS200 | opensea |

| | | | | | | | | |
|---|---|---|---|---|---|---|---|---|
| cg25495534 | -0.15 | 0.0006 | 0.55 | 0.39 | -0.15 | | IGR | opensea |
| cg21283066 | -0.15 | 0.0000 | 0.50 | 0.35 | -0.15 | | IGR | opensea |
| cg23500537 | -0.15 | 0.0001 | 0.49 | 0.33 | -0.15 | | IGR | opensea |
| cg13375589 | -0.15 | 0.0161 | 0.54 | 0.39 | -0.15 | SMTNL2 | TSS200 | shore |
| cg15131258 | -0.15 | 0.0286 | 0.60 | 0.45 | -0.15 | C20orf71 | 5'UTR | opensea |
| cg02723558 | -0.15 | 0.0039 | 0.55 | 0.40 | -0.15 | BDNFOS | Body | opensea |
| cg16732787 | -0.15 | 0.0003 | 0.56 | 0.41 | -0.15 | DUSP5P | Body | island |
| cg06013117 | -0.15 | 0.0000 | 0.78 | 0.63 | -0.15 | MSX2 | Body | shore |
| cg07964219 | 0.15 | 0.0015 | 0.37 | 0.52 | 0.15 | COL18A1 | Body | shore |
| cg17552333 | 0.15 | 0.0016 | 0.26 | 0.41 | 0.15 | | IGR | opensea |
| cg07008591 | 0.15 | 0.0069 | 0.67 | 0.82 | 0.15 | TEAD1 | Body | opensea |
| cg21708130 | 0.15 | 0.0107 | 0.29 | 0.44 | 0.15 | LRRFIP1 | Body | shelf |
| cg15633390 | 0.15 | 0.0042 | 0.39 | 0.54 | 0.15 | EIF4E | 5'UTR | shore |
| cg25123566 | 0.15 | 0.0080 | 0.69 | 0.84 | 0.15 | FAM113B | 5'UTR | opensea |
| cg20848291 | 0.15 | 0.0017 | 0.24 | 0.39 | 0.15 | ZAN | Body | opensea |
| cg12427162 | 0.15 | 0.0000 | 0.57 | 0.72 | 0.15 | SFT2D2 | Body | shelf |
| cg18992848 | 0.15 | 0.0007 | 0.27 | 0.42 | 0.15 | TPK1 | Body | shore |
| cg06853894 | 0.15 | 0.0157 | 0.59 | 0.74 | 0.15 | TNFRSF8 | 3'UTR | opensea |
| cg01649611 | 0.15 | 0.0201 | 0.55 | 0.71 | 0.15 | THADA | Body | opensea |
| cg14753094 | 0.15 | 0.0017 | 0.69 | 0.84 | 0.15 | HSD17B12 | Body | opensea |
| cg25467833 | 0.15 | 0.0004 | 0.54 | 0.69 | 0.15 | | IGR | opensea |
| cg08928408 | 0.15 | 0.0048 | 0.69 | 0.84 | 0.15 | | IGR | opensea |
| cg05349016 | 0.15 | 0.0017 | 0.38 | 0.53 | 0.15 | NMT1 | Body | opensea |
| cg02832512 | 0.15 | 0.0057 | 0.60 | 0.76 | 0.15 | FLJ22536 | Body | opensea |
| cg08913523 | 0.15 | 0.0005 | 0.45 | 0.60 | 0.15 | | IGR | opensea |
| cg05762671 | 0.15 | 0.0055 | 0.58 | 0.74 | 0.15 | KCTD19 | Body | opensea |
| cg13052638 | 0.15 | 0.0059 | 0.30 | 0.46 | 0.15 | | IGR | shelf |
| cg23295647 | 0.15 | 0.0207 | 0.40 | 0.56 | 0.15 | NPAS3 | Body | island |
| cg04642300 | 0.15 | 0.0009 | 0.43 | 0.58 | 0.15 | ARMC2 | Body | opensea |
| cg13327911 | 0.15 | 0.0266 | 0.67 | 0.82 | 0.15 | COL21A1 | Body | opensea |
| cg26197915 | 0.15 | 0.0005 | 0.20 | 0.35 | 0.15 | PTPRJ | Body | opensea |
| cg03354554 | 0.15 | 0.0067 | 0.29 | 0.44 | 0.15 | | IGR | shore |
| cg02573091 | 0.15 | 0.0096 | 0.26 | 0.42 | 0.15 | | IGR | shore |
| cg21369466 | 0.15 | 0.0073 | 0.65 | 0.81 | 0.15 | LANCL2 | Body | shelf |
| cg06570967 | 0.15 | 0.0034 | 0.76 | 0.91 | 0.15 | | IGR | opensea |
| cg06161600 | 0.15 | 0.0036 | 0.32 | 0.48 | 0.15 | BAIAP3 | Body | island |
| cg26177041 | 0.15 | 0.0000 | 0.20 | 0.35 | 0.15 | CAMK2D | Body | opensea |
| cg22473770 | 0.15 | 0.0063 | 0.53 | 0.68 | 0.15 | EVI2A | 5'UTR | opensea |
| cg13419330 | 0.15 | 0.0019 | 0.56 | 0.71 | 0.15 | IRAK2 | Body | opensea |
| cg07664000 | 0.16 | 0.0053 | 0.53 | 0.69 | 0.16 | TMIGD1 | 5'UTR | opensea |
| cg09228833 | 0.16 | 0.0201 | 0.48 | 0.64 | 0.16 | ZNF217 | TSS200 | shore |
| cg00993830 | 0.16 | 0.0010 | 0.38 | 0.53 | 0.16 | UBE2H | Body | opensea |
| cg25298319 | 0.16 | 0.0048 | 0.23 | 0.39 | 0.16 | | IGR | island |

| | | | | | | | | |
|---|---|---|---|---|---|---|---|---|
| cg01993169 | 0.16 | 0.0006 | 0.73 | 0.89 | 0.16 | | IGR | opensea |
| cg08939850 | 0.16 | 0.0054 | 0.59 | 0.74 | 0.16 | RPTOR | Body | shore |
| cg06478886 | 0.16 | 0.0324 | 0.32 | 0.48 | 0.16 | | IGR | shore |
| cg20968743 | 0.16 | 0.0079 | 0.31 | 0.46 | 0.16 | TSPAN18 | 5'UTR | opensea |
| cg00223245 | 0.16 | 0.0014 | 0.41 | 0.57 | 0.16 | | IGR | opensea |
| cg25616869 | 0.16 | 0.0017 | 0.54 | 0.70 | 0.16 | | IGR | opensea |
| cg06691616 | 0.16 | 0.0006 | 0.51 | 0.67 | 0.16 | | IGR | opensea |
| cg18156592 | 0.16 | 0.0018 | 0.27 | 0.43 | 0.16 | ARL6IP5 | Body | opensea |
| cg13017929 | 0.16 | 0.0039 | 0.64 | 0.80 | 0.16 | | IGR | opensea |
| cg18456803 | 0.16 | 0.0004 | 0.40 | 0.56 | 0.16 | ELF1 | TSS200 | opensea |
| cg00377497 | 0.16 | 0.0001 | 0.55 | 0.71 | 0.16 | TRIM35 | Body | shore |
| cg01116477 | 0.16 | 0.0000 | 0.59 | 0.75 | 0.16 | | IGR | opensea |
| cg05227773 | 0.16 | 0.0003 | 0.62 | 0.78 | 0.16 | ZFHX3 | 3'UTR | shelf |
| cg24736734 | 0.16 | 0.0005 | 0.26 | 0.42 | 0.16 | | IGR | opensea |
| cg10732871 | 0.16 | 0.0016 | 0.29 | 0.45 | 0.16 | GPX4 | TSS1500 | shore |
| cg03987648 | 0.16 | 0.0046 | 0.66 | 0.82 | 0.16 | MAML3 | Body | opensea |
| cg07285237 | 0.16 | 0.0178 | 0.69 | 0.85 | 0.16 | | IGR | opensea |
| cg10110335 | 0.16 | 0.0050 | 0.53 | 0.69 | 0.16 | SYN2 | Body | shelf |
| cg10557907 | 0.16 | 0.0013 | 0.43 | 0.59 | 0.16 | PRDM16 | Body | opensea |
| cg27310092 | 0.16 | 0.0042 | 0.30 | 0.46 | 0.16 | | IGR | opensea |
| cg07813142 | 0.16 | 0.0022 | 0.10 | 0.26 | 0.16 | SP5 | Body | island |
| cg14701867 | 0.16 | 0.0445 | 0.45 | 0.61 | 0.16 | ZNF365 | Body | opensea |
| cg13001142 | 0.16 | 0.0000 | 0.62 | 0.78 | 0.16 | STXBP5 | Body | shelf |
| cg09450153 | 0.16 | 0.0428 | 0.56 | 0.72 | 0.16 | CREB5 | Body | opensea |
| cg00754989 | 0.16 | 0.0281 | 0.53 | 0.69 | 0.16 | | IGR | opensea |
| cg24508426 | 0.16 | 0.0016 | 0.23 | 0.39 | 0.16 | | IGR | island |
| cg15852787 | 0.16 | 0.0000 | 0.68 | 0.84 | 0.16 | FRMD6 | 5'UTR | opensea |
| cg01409343 | 0.16 | 0.0008 | 0.38 | 0.54 | 0.16 | TMEM49 | Body | opensea |
| cg05373263 | 0.16 | 0.0251 | 0.56 | 0.72 | 0.16 | | IGR | shore |
| cg26284735 | 0.16 | 0.0006 | 0.46 | 0.63 | 0.16 | | IGR | shelf |
| cg01412970 | 0.16 | 0.0335 | 0.28 | 0.44 | 0.16 | PLD6 | 1stExon | island |
| cg06769820 | 0.16 | 0.0212 | 0.40 | 0.56 | 0.16 | | IGR | opensea |
| cg13315147 | 0.16 | 0.0343 | 0.13 | 0.30 | 0.16 | CYP2E1 | Body | island |
| cg02351277 | 0.16 | 0.0031 | 0.27 | 0.43 | 0.16 | | IGR | opensea |
| cg24314564 | 0.16 | 0.0312 | 0.56 | 0.72 | 0.16 | | IGR | shelf |
| cg09435170 | 0.16 | 0.0000 | 0.14 | 0.30 | 0.16 | | IGR | opensea |
| cg24475182 | 0.16 | 0.0007 | 0.22 | 0.38 | 0.16 | | IGR | opensea |
| cg20899781 | 0.16 | 0.0000 | 0.35 | 0.51 | 0.16 | | IGR | opensea |
| cg10763234 | 0.16 | 0.0011 | 0.22 | 0.38 | 0.16 | | IGR | island |
| cg02849956 | 0.16 | 0.0019 | 0.56 | 0.72 | 0.16 | | IGR | shore |
| cg11029367 | 0.16 | 0.0016 | 0.64 | 0.80 | 0.16 | HEG1 | Body | opensea |
| cg16429725 | 0.16 | 0.0001 | 0.52 | 0.68 | 0.16 | KIFC3 | Body | shelf |
| cg04760448 | 0.16 | 0.0046 | 0.49 | 0.65 | 0.16 | COL18A1 | Body | island |

| | | | | | | | | |
|---|---|---|---|---|---|---|---|---|
| cg15418499 | 0.16 | 0.0007 | 0.41 | 0.57 | 0.16 | IL18 | 5'UTR | opensea |
| cg23920246 | 0.16 | 0.0011 | 0.73 | 0.89 | 0.16 | | IGR | opensea |
| cg17611046 | 0.16 | 0.0008 | 0.61 | 0.77 | 0.16 | FARS2 | Body | opensea |
| cg16337566 | 0.16 | 0.0001 | 0.25 | 0.41 | 0.16 | PINX1 | Body | opensea |
| cg26361533 | 0.16 | 0.0000 | 0.15 | 0.31 | 0.16 | CACNA1C | Body | opensea |
| cg21818891 | 0.17 | 0.0003 | 0.76 | 0.93 | 0.17 | SLC1A2 | Body | opensea |
| cg00960147 | 0.17 | 0.0011 | 0.67 | 0.84 | 0.17 | | IGR | opensea |
| cg01195564 | 0.17 | 0.0028 | 0.58 | 0.74 | 0.17 | | IGR | opensea |
| cg16911981 | 0.17 | 0.0005 | 0.21 | 0.38 | 0.17 | AUTS2 | Body | island |
| cg05875421 | 0.17 | 0.0015 | 0.69 | 0.86 | 0.17 | GPR68 | 5'UTR | opensea |
| cg04926881 | 0.17 | 0.0000 | 0.19 | 0.36 | 0.17 | | IGR | opensea |
| cg14276379 | 0.17 | 0.0037 | 0.25 | 0.42 | 0.17 | C9orf3 | Body | opensea |
| cg05492387 | 0.17 | 0.0007 | 0.57 | 0.73 | 0.17 | RAP1GDS1 | Body | opensea |
| cg12024811 | 0.17 | 0.0000 | 0.60 | 0.77 | 0.17 | KALRN | Body | opensea |
| cg09548403 | 0.17 | 0.0049 | 0.28 | 0.45 | 0.17 | | IGR | opensea |
| cg22274117 | 0.17 | 0.0279 | 0.41 | 0.58 | 0.17 | ATXN1 | 5'UTR | opensea |
| cg00858840 | 0.17 | 0.0002 | 0.18 | 0.35 | 0.17 | SP5 | Body | island |
| cg26203572 | 0.17 | 0.0001 | 0.64 | 0.81 | 0.17 | | IGR | opensea |
| cg09966895 | 0.17 | 0.0092 | 0.66 | 0.83 | 0.17 | ODZ4 | Body | opensea |
| cg24178897 | 0.17 | 0.0024 | 0.57 | 0.74 | 0.17 | | IGR | opensea |
| cg20651995 | 0.17 | 0.0012 | 0.53 | 0.70 | 0.17 | | IGR | opensea |
| cg11562411 | 0.17 | 0.0001 | 0.24 | 0.41 | 0.17 | SCUBE3 | Body | opensea |
| cg00601450 | 0.17 | 0.0051 | 0.44 | 0.61 | 0.17 | | IGR | shore |
| cg26504263 | 0.17 | 0.0000 | 0.59 | 0.76 | 0.17 | ANKRD6 | 5'UTR | opensea |
| cg24113973 | 0.17 | 0.0086 | 0.20 | 0.37 | 0.17 | | IGR | shelf |
| cg17800426 | 0.17 | 0.0000 | 0.66 | 0.83 | 0.17 | MYOZ3 | TSS1500 | shelf |
| cg18263166 | 0.17 | 0.0104 | 0.57 | 0.74 | 0.17 | | IGR | opensea |
| cg11868461 | 0.17 | 0.0055 | 0.36 | 0.54 | 0.17 | | IGR | opensea |
| cg26893861 | 0.17 | 0.0337 | 0.21 | 0.39 | 0.17 | DUSP3 | 3'UTR | opensea |
| cg06457736 | 0.17 | 0.0031 | 0.49 | 0.66 | 0.17 | HRH1 | TSS200 | opensea |
| cg18700940 | 0.17 | 0.0000 | 0.12 | 0.30 | 0.17 | MAP3K14 | 5'UTR | opensea |
| cg21335012 | 0.17 | 0.0000 | 0.69 | 0.87 | 0.17 | | IGR | opensea |
| cg16683060 | 0.18 | 0.0192 | 0.50 | 0.68 | 0.18 | UBAC2 | Body | opensea |
| cg14708411 | 0.18 | 0.0339 | 0.72 | 0.89 | 0.18 | SLC12A7 | Body | island |
| cg23821329 | 0.18 | 0.0069 | 0.44 | 0.62 | 0.18 | VIM | TSS1500 | shore |
| cg24772753 | 0.18 | 0.0008 | 0.17 | 0.35 | 0.18 | SP5 | Body | island |
| cg00773142 | 0.18 | 0.0036 | 0.38 | 0.55 | 0.18 | PLCG2 | Body | opensea |
| cg07805542 | 0.18 | 0.0015 | 0.48 | 0.66 | 0.18 | PIK3CD | Body | shelf |
| cg02992067 | 0.18 | 0.0000 | 0.68 | 0.86 | 0.18 | FTO | Body | opensea |
| cg24062389 | 0.18 | 0.0001 | 0.61 | 0.79 | 0.18 | BIVM | Body | opensea |
| cg25773259 | 0.18 | 0.0002 | 0.61 | 0.79 | 0.18 | | IGR | opensea |
| cg11118962 | 0.18 | 0.0008 | 0.44 | 0.62 | 0.18 | | IGR | opensea |
| cg08426157 | 0.18 | 0.0004 | 0.64 | 0.82 | 0.18 | HDAC9 | Body | opensea |

| | | | | | | | | |
|---|---|---|---|---|---|---|---|---|
| cg21341586 | 0.18 | 0.0027 | 0.44 | 0.62 | 0.18 | EIF4E | 5'UTR | shore |
| cg16002660 | 0.18 | 0.0001 | 0.42 | 0.60 | 0.18 | LOC284009 | Body | opensea |
| cg07986257 | 0.18 | 0.0000 | 0.63 | 0.81 | 0.18 | | IGR | opensea |
| cg21860675 | 0.18 | 0.0011 | 0.66 | 0.84 | 0.18 | FOXP1 | 5'UTR | opensea |
| cg27640794 | 0.18 | 0.0102 | 0.43 | 0.61 | 0.18 | PALLD | 5'UTR | shelf |
| cg07677157 | 0.18 | 0.0328 | 0.62 | 0.80 | 0.18 | | IGR | opensea |
| cg12686055 | 0.18 | 0.0000 | 0.21 | 0.39 | 0.18 | ANO6 | Body | opensea |
| cg02500300 | 0.18 | 0.0001 | 0.17 | 0.35 | 0.18 | STOX2 | 1stExon | island |
| cg10316899 | 0.18 | 0.0001 | 0.35 | 0.53 | 0.18 | MACF1 | Body | opensea |
| cg27549186 | 0.18 | 0.0015 | 0.62 | 0.81 | 0.18 | TIMP2 | Body | opensea |
| cg24693760 | 0.18 | 0.0309 | 0.35 | 0.53 | 0.18 | | IGR | opensea |
| cg03548415 | 0.18 | 0.0033 | 0.36 | 0.54 | 0.18 | | IGR | opensea |
| cg03839782 | 0.18 | 0.0004 | 0.62 | 0.80 | 0.18 | FAM65B | 5'UTR | opensea |
| cg10862468 | 0.18 | 0.0238 | 0.18 | 0.37 | 0.18 | CYP2E1 | Body | island |
| cg16336556 | 0.18 | 0.0034 | 0.55 | 0.73 | 0.18 | LTBP1 | Body | opensea |
| cg17178175 | 0.19 | 0.0026 | 0.34 | 0.52 | 0.19 | NFE2L2 | Body | opensea |
| cg21727223 | 0.19 | 0.0000 | 0.30 | 0.49 | 0.19 | | IGR | opensea |
| cg06060522 | 0.19 | 0.0029 | 0.40 | 0.59 | 0.19 | | IGR | island |
| cg01413054 | 0.19 | 0.0314 | 0.56 | 0.74 | 0.19 | CAB39 | Body | opensea |
| cg01923775 | 0.19 | 0.0001 | 0.40 | 0.59 | 0.19 | PALLD | Body | opensea |
| cg02515217 | 0.19 | 0.0001 | 0.42 | 0.61 | 0.19 | MIR21 | TSS200 | opensea |
| cg19770281 | 0.19 | 0.0094 | 0.59 | 0.78 | 0.19 | | IGR | opensea |
| cg11809668 | 0.19 | 0.0001 | 0.62 | 0.81 | 0.19 | KIAA0922 | Body | opensea |
| cg07093324 | 0.19 | 0.0134 | 0.63 | 0.82 | 0.19 | ACTR3 | Body | shelf |
| cg11445109 | 0.19 | 0.0184 | 0.10 | 0.29 | 0.19 | CYP2E1 | Body | shore |
| cg25015038 | 0.19 | 0.0033 | 0.38 | 0.57 | 0.19 | | IGR | opensea |
| cg11608150 | 0.19 | 0.0233 | 0.25 | 0.45 | 0.19 | | IGR | shore |
| cg12609785 | 0.19 | 0.0015 | 0.40 | 0.59 | 0.19 | | IGR | shore |
| cg23099839 | 0.20 | 0.0032 | 0.23 | 0.43 | 0.20 | | IGR | opensea |
| cg14986890 | 0.20 | 0.0015 | 0.49 | 0.69 | 0.20 | RARRES1 | Body | opensea |
| cg27149179 | 0.20 | 0.0000 | 0.34 | 0.53 | 0.20 | | IGR | opensea |
| cg21144063 | 0.20 | 0.0007 | 0.29 | 0.49 | 0.20 | | IGR | opensea |
| cg11906781 | 0.20 | 0.0022 | 0.31 | 0.51 | 0.20 | BRE | Body | opensea |
| cg23201812 | 0.20 | 0.0012 | 0.30 | 0.50 | 0.20 | | IGR | opensea |
| cg16233797 | 0.20 | 0.0013 | 0.61 | 0.82 | 0.20 | | IGR | opensea |
| cg24367957 | 0.21 | 0.0066 | 0.33 | 0.54 | 0.21 | | IGR | opensea |
| cg20905796 | 0.21 | 0.0100 | 0.47 | 0.68 | 0.21 | | IGR | opensea |
| cg15694704 | 0.21 | 0.0003 | 0.45 | 0.67 | 0.21 | RPTOR | Body | shelf |
| cg24921221 | 0.21 | 0.0000 | 0.16 | 0.37 | 0.21 | LONRF1 | Body | opensea |
| cg05469819 | 0.21 | 0.0001 | 0.56 | 0.78 | 0.21 | | IGR | opensea |
| cg08085267 | 0.22 | 0.0005 | 0.13 | 0.34 | 0.22 | C17orf57 | 5'UTR | shore |
| cg02782634 | 0.22 | 0.0000 | 0.27 | 0.49 | 0.22 | TMEM49 | Body | opensea |
| cg10587082 | 0.22 | 0.0020 | 0.44 | 0.66 | 0.22 | PLXNA2 | Body | opensea |

| | | | | | | | | |
|---|---|---|---|---|---|---|---|---|
| cg04236915 | 0.23 | 0.0000 | 0.42 | 0.65 | 0.23 | ECE1 | Body | opensea |
| cg12403162 | 0.23 | 0.0000 | 0.26 | 0.49 | 0.23 | ABLIM1 | Body | shelf |
| cg05194426 | 0.23 | 0.0205 | 0.25 | 0.48 | 0.23 | CYP2E1 | Body | shore |
| cg12592365 | 0.23 | 0.0004 | 0.36 | 0.60 | 0.23 | RPTOR | Body | opensea |
| cg04470054 | 0.24 | 0.0031 | 0.42 | 0.66 | 0.24 | RPTOR | Body | shore |
| cg25588844 | 0.24 | 0.0001 | 0.40 | 0.64 | 0.24 | TAF1B | Body | opensea |
| cg27477494 | 0.24 | 0.0002 | 0.33 | 0.57 | 0.24 | | IGR | opensea |
| cg17311132 | 0.24 | 0.0005 | 0.57 | 0.81 | 0.24 | | IGR | opensea |
| cg19683494 | 0.25 | 0.0088 | 0.36 | 0.60 | 0.25 | | IGR | shore |
| cg17749961 | 0.28 | 0.0000 | 0.04 | 0.31 | 0.28 | LCLAT1 | TSS1500 | shore |
| cg12454169 | 0.30 | 0.0000 | 0.12 | 0.42 | 0.30 | LCLAT1 | TSS1500 | shore |
| cg22443212 | 0.36 | 0.0023 | 0.32 | 0.68 | 0.36 | RNF213 | Body | opensea |
| cg15652532 | 0.36 | 0.0001 | 0.16 | 0.53 | 0.36 | LCLAT1 | TSS1500 | shore |

**Supplemental Table 3.** Differentially methylated positions in Non-ischemic HF vs. Non-Failing (adjusted p-value <0.05, abs (delta beta) > 15%)

| CpG Site | log FC | Adj. p-val | NICM | NF | Delta B | Gene | Feature | CGI |
|---|---|---|---|---|---|---|---|---|
| cg05528899 | 0.30 | 0.0188 | 0.65 | 0.35 | -0.30125 | | IGR | island |
| cg26536949 | 0.25 | 0.0134 | 0.75 | 0.51 | -0.24523 | | IGR | island |
| cg10327440 | 0.21 | 0.0281 | 0.63 | 0.42 | -0.21488 | CDC42BPA | 3'UTR | opensea |
| cg22459517 | 0.21 | 0.0110 | 0.50 | 0.29 | -0.21221 | EPS8L1 | TSS200 | opensea |
| cg07167872 | 0.17 | 0.0325 | 0.41 | 0.24 | -0.16924 | PM20D1 | TSS200 | shore |
| cg02928365 | 0.17 | 0.0000 | 0.43 | 0.26 | -0.16878 | HLX | Body | shore |
| cg13283845 | 0.16 | 0.0169 | 0.73 | 0.57 | -0.16269 | | IGR | shore |
| cg02497785 | 0.16 | 0.0004 | 0.66 | 0.51 | -0.15689 | ABCA13 | Body | island |
| cg10507304 | 0.15 | 0.0001 | 0.42 | 0.26 | -0.15134 | | IGR | opensea |
| cg22056094 | 0.15 | 0.0003 | 0.67 | 0.52 | -0.15093 | PCDHGA2 | 1stExon | shore |
| cg10316899 | -0.15 | 0.0001 | 0.38 | 0.53 | 0.150102 | MACF1 | Body | opensea |
| cg04470054 | -0.15 | 0.0018 | 0.51 | 0.66 | 0.150844 | RPTOR | Body | shore |
| cg12403162 | -0.15 | 0.0004 | 0.34 | 0.49 | 0.153153 | ABLIM1 | Body | shelf |
| cg04236915 | -0.15 | 0.0000 | 0.49 | 0.65 | 0.154455 | ECE1 | Body | opensea |
| cg00601450 | -0.16 | 0.0020 | 0.45 | 0.61 | 0.15859 | | IGR | shore |
| cg25588844 | -0.16 | 0.0000 | 0.48 | 0.64 | 0.159595 | TAF1B | Body | opensea |
| cg25755428 | -0.16 | 0.0080 | 0.10 | 0.26 | 0.163686 | MRI1 | TSS1500 | island |
| cg16474696 | -0.17 | 0.0113 | 0.24 | 0.41 | 0.165264 | MRI1 | TSS1500 | shore |
| cg23099839 | -0.17 | 0.0022 | 0.26 | 0.43 | 0.169996 | | IGR | opensea |
| cg12592365 | -0.17 | 0.0005 | 0.42 | 0.60 | 0.174614 | RPTOR | Body | opensea |
| cg08085267 | -0.18 | 0.0011 | 0.16 | 0.34 | 0.179996 | C17orf57 | 5'UTR | shore |
| cg27477494 | -0.18 | 0.0003 | 0.39 | 0.57 | 0.181382 | | IGR | opensea |
| cg24693760 | -0.19 | 0.0105 | 0.34 | 0.53 | 0.192978 | | IGR | opensea |
| cg17749961 | -0.20 | 0.0049 | 0.12 | 0.31 | 0.19739 | LCLAT1 | TSS1500 | shore |
| cg02782634 | -0.20 | 0.0000 | 0.29 | 0.49 | 0.198254 | TMEM49 | Body | opensea |
| cg07147204 | -0.20 | 0.0001 | 0.75 | 0.95 | 0.198727 | | IGR | opensea |
| cg12454169 | -0.20 | 0.0208 | 0.22 | 0.42 | 0.200822 | LCLAT1 | TSS1500 | shore |
| cg15532640 | -0.21 | 0.0058 | 0.20 | 0.41 | 0.21394 | | IGR | opensea |
| cg20905796 | -0.22 | 0.0022 | 0.46 | 0.68 | 0.218515 | | IGR | opensea |
| cg19318364 | -0.23 | 0.0035 | 0.65 | 0.87 | 0.228456 | | IGR | opensea |
| cg19683494 | -0.23 | 0.0029 | 0.37 | 0.60 | 0.234184 | | IGR | shore |
| cg15652532 | -0.24 | 0.0160 | 0.28 | 0.53 | 0.243969 | LCLAT1 | TSS1500 | shore |
| cg22443212 | -0.31 | 0.0005 | 0.37 | 0.68 | 0.314602 | RNF213 | Body | opensea |

**Supplemental Table 4.** Differentially methylated positions common to Ischemic and Non-Ischemic HF (adjusted p-value <0.05, abs (delta beta) > 10%)

| CpG Site | Delta ICM | Delta NICM | Methylation | Gene | Feature | CGI |
|---|---|---|---|---|---|---|
| cg22443212 | -0.36 | -0.31 | Hypomethylated | RNF213 | Body | opensea |
| cg15652532 | -0.36 | -0.24 | Hypomethylated | LCLAT1 | TSS1500 | shore |
| cg12454169 | -0.30 | -0.20 | Hypomethylated | LCLAT1 | TSS1500 | shore |
| cg19683494 | -0.25 | -0.23 | Hypomethylated | | IGR | shore |
| cg17749961 | -0.28 | -0.20 | Hypomethylated | LCLAT1 | TSS1500 | shore |
| cg20905796 | -0.21 | -0.22 | Hypomethylated | | IGR | opensea |
| cg27477494 | -0.24 | -0.18 | Hypomethylated | | IGR | opensea |
| cg02782634 | -0.22 | -0.20 | Hypomethylated | TMEM49 | Body | opensea |
| cg12592365 | -0.23 | -0.17 | Hypomethylated | RPTOR | Body | opensea |
| cg25588844 | -0.24 | -0.16 | Hypomethylated | TAF1B | Body | opensea |
| cg08085267 | -0.22 | -0.18 | Hypomethylated | C17orf57 | 5'UTR | shore |
| cg04470054 | -0.24 | -0.15 | Hypomethylated | RPTOR | Body | shore |
| cg04236915 | -0.23 | -0.15 | Hypomethylated | ECE1 | Body | opensea |
| cg12403162 | -0.23 | -0.15 | Hypomethylated | ABLIM1 | Body | shelf |
| cg24693760 | -0.18 | -0.19 | Hypomethylated | | IGR | opensea |
| cg17311132 | -0.24 | -0.12 | Hypomethylated | | IGR | opensea |
| cg23099839 | -0.20 | -0.17 | Hypomethylated | | IGR | opensea |
| cg10587082 | -0.22 | -0.12 | Hypomethylated | PLXNA2 | Body | opensea |
| cg15694704 | -0.21 | -0.13 | Hypomethylated | RPTOR | Body | shelf |
| cg24921221 | -0.21 | -0.13 | Hypomethylated | LONRF1 | Body | opensea |
| cg21144063 | -0.20 | -0.14 | Hypomethylated | | IGR | opensea |
| cg10316899 | -0.18 | -0.15 | Hypomethylated | MACF1 | Body | opensea |
| cg00601450 | -0.17 | -0.16 | Hypomethylated | | IGR | shore |
| cg12686055 | -0.18 | -0.15 | Hypomethylated | ANO6 | Body | opensea |
| cg27149179 | -0.20 | -0.13 | Hypomethylated | | IGR | opensea |
| cg05469819 | -0.21 | -0.11 | Hypomethylated | | IGR | opensea |
| cg23201812 | -0.20 | -0.13 | Hypomethylated | | IGR | opensea |
| cg24367957 | -0.21 | -0.12 | Hypomethylated | | IGR | opensea |
| cg01923775 | -0.19 | -0.14 | Hypomethylated | PALLD | Body | opensea |
| cg16233797 | -0.20 | -0.12 | Hypomethylated | | IGR | opensea |
| cg03839782 | -0.18 | -0.14 | Hypomethylated | FAM65B | 5'UTR | opensea |
| cg11906781 | -0.20 | -0.11 | Hypomethylated | BRE | Body | opensea |
| cg02992067 | -0.18 | -0.13 | Hypomethylated | FTO | Body | opensea |
| cg07986257 | -0.18 | -0.13 | Hypomethylated | | IGR | opensea |
| cg24113973 | -0.17 | -0.14 | Hypomethylated | | IGR | shelf |
| cg16337566 | -0.16 | -0.14 | Hypomethylated | PINX1 | Body | opensea |
| cg21727223 | -0.19 | -0.12 | Hypomethylated | | IGR | opensea |
| cg02515217 | -0.19 | -0.12 | Hypomethylated | MIR21 | TSS200 | opensea |
| cg11809668 | -0.19 | -0.12 | Hypomethylated | KIAA0922 | Body | opensea |

| | | | | | | |
|---|---|---|---|---|---|---|
| cg17178175 | -0.19 | -0.11 | Hypomethylated | NFE2L2 | Body | opensea |
| cg14986890 | -0.20 | -0.10 | Hypomethylated | RARRES1 | Body | opensea |
| cg09435170 | -0.16 | -0.13 | Hypomethylated | | IGR | opensea |
| cg13001142 | -0.16 | -0.13 | Hypomethylated | STXBP5 | Body | shelf |
| cg04926881 | -0.17 | -0.13 | Hypomethylated | | IGR | opensea |
| cg16002660 | -0.18 | -0.11 | Hypomethylated | LOC284009 | Body | opensea |
| cg11118962 | -0.18 | -0.11 | Hypomethylated | | IGR | opensea |
| cg26504263 | -0.17 | -0.11 | Hypomethylated | ANKRD6 | 5'UTR | opensea |
| cg25773259 | -0.18 | -0.11 | Hypomethylated | | IGR | opensea |
| cg24062389 | -0.18 | -0.11 | Hypomethylated | BIVM | Body | opensea |
| cg01412970 | -0.16 | -0.12 | Hypomethylated | PLD6 | 1stExon | island |
| cg01195564 | -0.17 | -0.12 | Hypomethylated | | IGR | opensea |
| cg24772753 | -0.18 | -0.10 | Hypomethylated | SP5 | Body | island |
| cg02573091 | -0.15 | -0.13 | Hypomethylated | | IGR | shore |
| cg01116477 | -0.16 | -0.12 | Hypomethylated | | IGR | opensea |
| cg12024811 | -0.17 | -0.11 | Hypomethylated | KALRN | Body | opensea |
| cg08426157 | -0.18 | -0.10 | Hypomethylated | HDAC9 | Body | opensea |
| cg21335012 | -0.17 | -0.10 | Hypomethylated | | IGR | opensea |
| cg20848291 | -0.15 | -0.13 | Hypomethylated | ZAN | Body | opensea |
| cg01993169 | -0.16 | -0.12 | Hypomethylated | | IGR | opensea |
| cg11562411 | -0.17 | -0.11 | Hypomethylated | SCUBE3 | Body | opensea |
| cg17800426 | -0.17 | -0.10 | Hypomethylated | MYOZ3 | TSS1500 | shelf |
| cg26203572 | -0.17 | -0.11 | Hypomethylated | | IGR | opensea |
| cg26177041 | -0.15 | -0.12 | Hypomethylated | CAMK2D | Body | opensea |
| cg18700940 | -0.17 | -0.10 | Hypomethylated | MAP3K14 | 5'UTR | opensea |
| cg13173567 | -0.13 | -0.14 | Hypomethylated | DENND5A | Body | opensea |
| cg01409343 | -0.16 | -0.11 | Hypomethylated | TMEM49 | Body | opensea |
| cg15852787 | -0.16 | -0.11 | Hypomethylated | FRMD6 | 5'UTR | opensea |
| cg05227773 | -0.16 | -0.11 | Hypomethylated | ZFHX3 | 3'UTR | shelf |
| cg02351277 | -0.16 | -0.11 | Hypomethylated | | IGR | opensea |
| cg04760448 | -0.16 | -0.11 | Hypomethylated | COL18A1 | Body | island |
| cg25173405 | -0.15 | -0.12 | Hypomethylated | C17orf57 | 5'UTR | shore |
| cg26284735 | -0.16 | -0.11 | Hypomethylated | | IGR | shelf |
| cg07813142 | -0.16 | -0.11 | Hypomethylated | SP5 | Body | island |
| cg24475182 | -0.16 | -0.10 | Hypomethylated | | IGR | opensea |
| cg10557907 | -0.16 | -0.11 | Hypomethylated | PRDM16 | Body | opensea |
| cg10110335 | -0.16 | -0.11 | Hypomethylated | SYN2 | Body | shelf |
| cg24736734 | -0.16 | -0.11 | Hypomethylated | | IGR | opensea |
| cg06474225 | -0.15 | -0.12 | Hypomethylated | HTRA1 | Body | opensea |
| cg07506560 | -0.14 | -0.12 | Hypomethylated | | IGR | opensea |
| cg14977018 | -0.14 | -0.11 | Hypomethylated | TMEM44 | Body | opensea |
| cg13052638 | -0.15 | -0.10 | Hypomethylated | | IGR | shelf |
| cg03052078 | -0.14 | -0.12 | Hypomethylated | STXBP5 | Body | shore |

| | | | | | | |
|---|---|---|---|---|---|---|
| cg16377679 | -0.15 | -0.10 | Hypomethylated | PDE4D | Body | opensea |
| cg04057161 | -0.15 | -0.11 | Hypomethylated | | IGR | opensea |
| cg18842353 | -0.14 | -0.11 | Hypomethylated | | IGR | opensea |
| cg16085649 | -0.14 | -0.11 | Hypomethylated | AKAP13 | Body | opensea |
| cg01412419 | -0.15 | -0.10 | Hypomethylated | | IGR | opensea |
| cg06850787 | -0.11 | -0.14 | Hypomethylated | | IGR | opensea |
| cg05543593 | -0.14 | -0.11 | Hypomethylated | | IGR | opensea |
| cg08889114 | -0.13 | -0.11 | Hypomethylated | | IGR | opensea |
| cg19017553 | -0.14 | -0.10 | Hypomethylated | PARD3 | Body | opensea |
| cg15645888 | -0.14 | -0.11 | Hypomethylated | FBXO16 | 3'UTR | opensea |
| cg13636014 | -0.14 | -0.10 | Hypomethylated | | IGR | opensea |
| cg01589587 | -0.14 | -0.11 | Hypomethylated | BATF | Body | opensea |
| cg11802797 | -0.14 | -0.10 | Hypomethylated | KIF1B | 3'UTR | opensea |
| cg24631102 | -0.11 | -0.13 | Hypomethylated | NPS | Body | opensea |
| cg01351822 | -0.12 | -0.12 | Hypomethylated | UNC45A | 5'UTR | island |
| cg12868173 | -0.13 | -0.10 | Hypomethylated | | IGR | island |
| cg13221458 | -0.13 | -0.10 | Hypomethylated | SOD2 | Body | shore |
| cg11452329 | -0.13 | -0.11 | Hypomethylated | | IGR | opensea |
| cg04228083 | -0.11 | -0.13 | Hypomethylated | LOC100130872-SPON2 | TSS200 | shore |
| cg02564299 | -0.13 | -0.10 | Hypomethylated | ZBTB20 | 5'UTR | opensea |
| cg14111334 | -0.12 | -0.12 | Hypomethylated | | IGR | opensea |
| cg18363918 | -0.13 | -0.11 | Hypomethylated | IGLON5 | Body | shore |
| cg14713217 | -0.13 | -0.11 | Hypomethylated | | IGR | opensea |
| cg04128967 | -0.13 | -0.11 | Hypomethylated | | IGR | opensea |
| cg24611970 | -0.13 | -0.11 | Hypomethylated | | IGR | opensea |
| cg13051700 | -0.13 | -0.10 | Hypomethylated | MAGI2 | Body | opensea |
| cg10082398 | -0.12 | -0.11 | Hypomethylated | CLDN20 | 5'UTR | opensea |
| cg06938601 | -0.12 | -0.11 | Hypomethylated | TCERG1L | Body | opensea |
| cg02784232 | -0.12 | -0.11 | Hypomethylated | PHC3 | Body | shore |
| cg14387312 | -0.12 | -0.10 | Hypomethylated | | IGR | opensea |
| cg07791418 | -0.12 | -0.10 | Hypomethylated | | IGR | opensea |
| cg18942579 | -0.12 | -0.11 | Hypomethylated | TMEM49 | Body | opensea |
| cg00159243 | -0.12 | -0.10 | Hypomethylated | SELPLG | 5'UTR | opensea |
| cg15690347 | -0.12 | -0.10 | Hypomethylated | SPIB | Body | island |
| cg03665360 | -0.12 | -0.10 | Hypomethylated | | IGR | opensea |
| cg21106486 | -0.11 | -0.11 | Hypomethylated | CR1L | Body | island |
| cg20366549 | -0.11 | -0.11 | Hypomethylated | SCNN1A | Body | opensea |
| cg10974479 | -0.11 | -0.10 | Hypomethylated | MLN | TSS1500 | opensea |
| cg26937434 | -0.11 | -0.10 | Hypomethylated | ANKS4B | 1stExon | opensea |
| cg10296718 | -0.11 | -0.10 | Hypomethylated | ARHGEF10 | Body | opensea |
| cg15226275 | -0.10 | -0.11 | Hypomethylated | FRK | TSS200 | opensea |
| cg19590421 | -0.10 | -0.10 | Hypomethylated | | IGR | opensea |

| | | | | | | |
|---|---|---|---|---|---|---|
| cg06234051 | -0.10 | -0.10 | Hypomethylated | SOX9 | 3'UTR | shore |
| cg01404163 | 0.10 | 0.10 | Hypermethylated | TOX3 | TSS200 | shore |
| cg22025206 | 0.10 | 0.10 | Hypermethylated | SLC9A3 | Body | shore |
| cg02793451 | 0.11 | 0.10 | Hypermethylated | TOX3 | TSS1500 | shore |
| cg07741162 | 0.11 | 0.10 | Hypermethylated | PRDM6 | Body | shore |
| cg27353899 | 0.11 | 0.10 | Hypermethylated | MUC4 | Body | island |
| cg14065590 | 0.11 | 0.10 | Hypermethylated | PCDHB11 | TSS200 | shore |
| cg11948367 | 0.11 | 0.10 | Hypermethylated | | IGR | opensea |
| cg03776662 | 0.11 | 0.10 | Hypermethylated | PRDM6 | Body | island |
| cg27067781 | 0.11 | 0.10 | Hypermethylated | PRRT1 | 3'UTR | island |
| cg14141912 | 0.11 | 0.11 | Hypermethylated | ATOH8 | Body | opensea |
| cg16049600 | 0.10 | 0.12 | Hypermethylated | PCDHB11 | TSS200 | shore |
| cg14497054 | 0.12 | 0.10 | Hypermethylated | | IGR | island |
| cg00877329 | 0.12 | 0.11 | Hypermethylated | HPSE2 | TSS1500 | shelf |
| cg05826245 | 0.12 | 0.11 | Hypermethylated | STYK1 | TSS1500 | shore |
| cg04438997 | 0.12 | 0.10 | Hypermethylated | SOX9 | TSS1500 | shore |
| cg24127414 | 0.11 | 0.11 | Hypermethylated | PCDHB11 | 1stExon | shore |
| cg12864721 | 0.12 | 0.10 | Hypermethylated | C10orf41 | Body | island |
| cg10388307 | 0.12 | 0.11 | Hypermethylated | | IGR | opensea |
| cg11430077 | 0.13 | 0.10 | Hypermethylated | GATA3 | Body | shore |
| cg22041228 | 0.13 | 0.10 | Hypermethylated | HLX | Body | shore |
| cg15359163 | 0.12 | 0.11 | Hypermethylated | PRDM6 | Body | shore |
| cg04154653 | 0.13 | 0.11 | Hypermethylated | TTLL10 | 3'UTR | shore |
| cg01310397 | 0.12 | 0.11 | Hypermethylated | MUC4 | Body | shore |
| cg18713687 | 0.12 | 0.11 | Hypermethylated | MUC4 | Body | island |
| cg12211856 | 0.11 | 0.12 | Hypermethylated | SDCCAG8 | Body | island |
| cg15415259 | 0.13 | 0.11 | Hypermethylated | MGC34034 | Body | shore |
| cg22770911 | 0.13 | 0.10 | Hypermethylated | GATA3 | Body | shore |
| cg18463607 | 0.13 | 0.11 | Hypermethylated | EXOC1 | TSS1500 | shore |
| cg23596123 | 0.12 | 0.12 | Hypermethylated | PCDHB6 | 1stExon | shore |
| cg25720795 | 0.13 | 0.11 | Hypermethylated | | IGR | shore |
| cg14774440 | 0.14 | 0.11 | Hypermethylated | RAB11FIP1 | Body | opensea |
| cg01535205 | 0.14 | 0.10 | Hypermethylated | | IGR | opensea |
| cg11350586 | 0.13 | 0.12 | Hypermethylated | SOX9 | TSS1500 | shore |
| cg26884658 | 0.13 | 0.12 | Hypermethylated | | IGR | shelf |
| cg23289079 | 0.14 | 0.11 | Hypermethylated | PRDM6 | Body | shore |
| cg13829104 | 0.14 | 0.10 | Hypermethylated | TBX3 | Body | shore |
| cg16298867 | 0.13 | 0.12 | Hypermethylated | | IGR | opensea |
| cg26680989 | 0.14 | 0.11 | Hypermethylated | | IGR | opensea |
| cg07028950 | 0.12 | 0.13 | Hypermethylated | | IGR | opensea |
| cg19689427 | 0.13 | 0.12 | Hypermethylated | PCDHGA2 | 1stExon | shore |
| cg24441899 | 0.13 | 0.12 | Hypermethylated | SDK1 | Body | opensea |
| cg26648818 | 0.13 | 0.12 | Hypermethylated | TOX3 | TSS200 | shore |

| | | | | | | |
|---|---|---|---|---|---|---|
| cg21283066 | 0.15 | 0.10 | Hypermethylated | | IGR | opensea |
| cg16200531 | 0.16 | 0.10 | Hypermethylated | | IGR | opensea |
| cg02497785 | 0.11 | 0.16 | Hypermethylated | ABCA13 | Body | island |
| cg04392266 | 0.13 | 0.14 | Hypermethylated | | IGR | island |
| cg20618651 | 0.15 | 0.12 | Hypermethylated | EXOC1 | TSS1500 | shore |
| cg01107874 | 0.16 | 0.11 | Hypermethylated | C10orf41 | Body | island |
| cg20454002 | 0.15 | 0.12 | Hypermethylated | HLX | Body | shore |
| cg03329019 | 0.15 | 0.12 | Hypermethylated | | IGR | shore |
| cg11679455 | 0.16 | 0.11 | Hypermethylated | GATA3 | Body | island |
| cg23936410 | 0.16 | 0.11 | Hypermethylated | | IGR | opensea |
| cg10070864 | 0.17 | 0.10 | Hypermethylated | | IGR | shelf |
| cg10507304 | 0.14 | 0.15 | Hypermethylated | | IGR | opensea |
| cg22056094 | 0.14 | 0.15 | Hypermethylated | PCDHGA2 | 1stExon | shore |
| cg13752114 | 0.16 | 0.14 | Hypermethylated | MUC4 | Body | island |
| cg17436134 | 0.17 | 0.14 | Hypermethylated | | IGR | shore |
| cg19657945 | 0.17 | 0.15 | Hypermethylated | | IGR | shore |
| cg12563372 | 0.18 | 0.13 | Hypermethylated | | IGR | shore |
| cg14318858 | 0.17 | 0.14 | Hypermethylated | CPT1C | Body | island |
| cg12777520 | 0.20 | 0.12 | Hypermethylated | LMX1B | Body | island |
| cg02928365 | 0.18 | 0.17 | Hypermethylated | HLX | Body | shore |
| cg13283845 | 0.20 | 0.16 | Hypermethylated | | IGR | shore |
| cg04811114 | 0.26 | 0.14 | Hypermethylated | LGR6 | TSS200 | opensea |
| cg26536949 | 0.25 | 0.25 | Hypermethylated | | IGR | island |
| cg07167872 | 0.34 | 0.17 | Hypermethylated | PM20D1 | TSS200 | shore |
| cg05528899 | 0.33 | 0.30 | Hypermethylated | | IGR | island |

**Supplemental Table 5.** Differentially methylated positions in Pre-LVAD vs. Non-Failing (adjusted p-value <0.05, abs (delta beta) > 15%)

| CpG Site | log FC | Adj. p-val | Pre-LVAD | NF | Delta B | Gene | Feature | CGI |
|---|---|---|---|---|---|---|---|---|
| cg07167872 | 0.23 | 0.0202 | 0.47 | 0.24 | -0.23 | PM20D1 | TSS200 | shore |
| cg14159672 | 0.21 | 0.0486 | 0.43 | 0.23 | -0.21 | PM20D1 | 1stExon | island |
| cg04811114 | 0.20 | 0.0026 | 0.41 | 0.21 | -0.20 | LGR6 | TSS200 | opensea |
| cg10507304 | 0.18 | 0.0003 | 0.44 | 0.26 | -0.18 | | IGR | opensea |
| cg02928365 | 0.18 | 0.0007 | 0.44 | 0.26 | -0.18 | HLX | Body | shore |
| cg12777520 | 0.17 | 0.0013 | 0.52 | 0.35 | -0.17 | LMX1B | Body | island |
| cg17436134 | 0.17 | 0.0024 | 0.47 | 0.30 | -0.17 | | IGR | shore |
| cg14318858 | 0.17 | 0.0003 | 0.77 | 0.60 | -0.17 | CPT1C | Body | island |
| cg11679455 | 0.16 | 0.0001 | 0.68 | 0.51 | -0.16 | GATA3 | Body | island |
| cg12563372 | 0.16 | 0.0004 | 0.48 | 0.32 | -0.16 | | IGR | shore |
| cg24503407 | 0.16 | 0.0323 | 0.56 | 0.40 | -0.16 | PM20D1 | TSS1500 | shore |
| cg03329019 | 0.16 | 0.0001 | 0.62 | 0.47 | -0.16 | | IGR | shore |
| cg09559189 | 0.16 | 0.0012 | 0.37 | 0.22 | -0.16 | EBF2 | Body | shore |
| cg22280475 | 0.16 | 0.0017 | 0.34 | 0.18 | -0.16 | EBF2 | Body | island |
| cg12583076 | 0.15 | 0.0218 | 0.54 | 0.39 | -0.15 | RASSF3 | Body | opensea |
| cg03731740 | 0.15 | 0.0025 | 0.54 | 0.39 | -0.15 | YTHDF2 | TSS1500 | shore |
| cg04492228 | 0.15 | 0.0000 | 0.57 | 0.42 | -0.15 | GATA3 | Body | shore |
| cg25015038 | -0.15 | 0.0177 | 0.42 | 0.57 | 0.15 | | IGR | opensea |
| cg25773259 | -0.15 | 0.0016 | 0.64 | 0.79 | 0.15 | | IGR | opensea |
| cg18263166 | -0.15 | 0.0084 | 0.59 | 0.74 | 0.15 | | IGR | opensea |
| cg26203572 | -0.15 | 0.0007 | 0.66 | 0.81 | 0.15 | | IGR | opensea |
| cg24772753 | -0.15 | 0.0073 | 0.20 | 0.35 | 0.15 | SP5 | Body | island |
| cg10557907 | -0.15 | 0.0080 | 0.44 | 0.59 | 0.15 | PRDM16 | Body | opensea |
| cg11235602 | -0.15 | 0.0007 | 0.24 | 0.39 | 0.15 | MOBP | Body | island |
| cg11809668 | -0.15 | 0.0030 | 0.65 | 0.81 | 0.15 | KIAA0922 | Body | opensea |
| cg16683060 | -0.15 | 0.0191 | 0.52 | 0.68 | 0.15 | UBAC2 | Body | opensea |
| cg16337566 | -0.15 | 0.0002 | 0.26 | 0.41 | 0.15 | PINX1 | Body | opensea |
| cg00117018 | -0.16 | 0.0165 | 0.50 | 0.65 | 0.16 | ZNF251 | Body | island |
| cg03839782 | -0.16 | 0.0021 | 0.64 | 0.80 | 0.16 | FAM65B | 5'UTR | opensea |
| cg13052638 | -0.16 | 0.0060 | 0.30 | 0.46 | 0.16 | | IGR | shelf |
| cg08159989 | -0.16 | 0.0036 | 0.50 | 0.66 | 0.16 | KLHDC8A | TSS200 | opensea |
| cg04760448 | -0.16 | 0.0078 | 0.50 | 0.65 | 0.16 | COL18A1 | Body | island |
| cg07677157 | -0.16 | 0.0323 | 0.64 | 0.80 | 0.16 | | IGR | opensea |
| cg03548415 | -0.16 | 0.0068 | 0.39 | 0.54 | 0.16 | | IGR | opensea |
| cg07506560 | -0.16 | 0.0015 | 0.58 | 0.74 | 0.16 | | IGR | opensea |
| cg24921221 | -0.16 | 0.0014 | 0.21 | 0.37 | 0.16 | LONRF1 | Body | opensea |
| cg12686055 | -0.16 | 0.0001 | 0.23 | 0.39 | 0.16 | ANO6 | Body | opensea |
| cg16233797 | -0.16 | 0.0060 | 0.66 | 0.82 | 0.16 | | IGR | opensea |
| cg02351277 | -0.16 | 0.0031 | 0.28 | 0.43 | 0.16 | | IGR | opensea |

| | | | | | | | | |
|---|---|---|---|---|---|---|---|---|
| cg27149179 | -0.16 | 0.0013 | 0.37 | 0.53 | 0.16 | | IGR | opensea |
| cg06060522 | -0.16 | 0.0288 | 0.43 | 0.59 | 0.16 | | IGR | island |
| cg05469819 | -0.16 | 0.0043 | 0.62 | 0.78 | 0.16 | | IGR | opensea |
| cg04236915 | -0.16 | 0.0014 | 0.48 | 0.65 | 0.16 | ECE1 | Body | opensea |
| cg07147204 | -0.16 | 0.0054 | 0.78 | 0.95 | 0.16 | | IGR | opensea |
| cg11906781 | -0.17 | 0.0067 | 0.34 | 0.51 | 0.17 | BRE | Body | opensea |
| cg02573091 | -0.17 | 0.0158 | 0.25 | 0.42 | 0.17 | | IGR | shore |
| cg24367957 | -0.17 | 0.0122 | 0.37 | 0.54 | 0.17 | | IGR | opensea |
| cg17178175 | -0.17 | 0.0041 | 0.35 | 0.52 | 0.17 | NFE2L2 | Body | opensea |
| cg16336556 | -0.17 | 0.0029 | 0.56 | 0.73 | 0.17 | LTBP1 | Body | opensea |
| cg23201812 | -0.17 | 0.0034 | 0.33 | 0.50 | 0.17 | | IGR | opensea |
| cg02500300 | -0.18 | 0.0002 | 0.18 | 0.35 | 0.18 | STOX2 | 1stExon | island |
| cg04470054 | -0.18 | 0.0185 | 0.48 | 0.66 | 0.18 | RPTOR | Body | shore |
| cg23821329 | -0.18 | 0.0067 | 0.44 | 0.62 | 0.18 | VIM | TSS1500 | shore |
| cg21144063 | -0.18 | 0.0042 | 0.31 | 0.49 | 0.18 | | IGR | opensea |
| cg12403162 | -0.19 | 0.0014 | 0.31 | 0.49 | 0.19 | ABLIM1 | Body | shelf |
| cg01923775 | -0.19 | 0.0002 | 0.40 | 0.59 | 0.19 | PALLD | Body | opensea |
| cg17311132 | -0.19 | 0.0022 | 0.62 | 0.81 | 0.19 | | IGR | opensea |
| cg15694704 | -0.19 | 0.0015 | 0.48 | 0.67 | 0.19 | RPTOR | Body | shelf |
| cg15532640 | -0.19 | 0.0470 | 0.22 | 0.41 | 0.19 | | IGR | opensea |
| cg01412970 | -0.19 | 0.0048 | 0.25 | 0.44 | 0.19 | PLD6 | 1stExon | island |
| cg10587082 | -0.20 | 0.0113 | 0.47 | 0.66 | 0.20 | PLXNA2 | Body | opensea |
| cg00601450 | -0.20 | 0.0059 | 0.41 | 0.61 | 0.20 | | IGR | shore |
| cg23099839 | -0.20 | 0.0092 | 0.23 | 0.43 | 0.20 | | IGR | opensea |
| cg02782634 | -0.20 | 0.0001 | 0.28 | 0.49 | 0.20 | TMEM49 | Body | opensea |
| cg17749961 | -0.20 | 0.0280 | 0.11 | 0.31 | 0.20 | LCLAT1 | TSS1500 | shore |
| cg24693760 | -0.20 | 0.0270 | 0.32 | 0.53 | 0.20 | | IGR | opensea |
| cg27477494 | -0.21 | 0.0017 | 0.36 | 0.57 | 0.21 | | IGR | opensea |
| cg20905796 | -0.21 | 0.0366 | 0.47 | 0.68 | 0.21 | | IGR | opensea |
| cg24113973 | -0.21 | 0.0028 | 0.16 | 0.37 | 0.21 | | IGR | shelf |
| cg25588844 | -0.21 | 0.0004 | 0.43 | 0.64 | 0.21 | TAF1B | Body | opensea |
| cg08085267 | -0.22 | 0.0018 | 0.13 | 0.34 | 0.22 | C17orf57 | 5'UTR | shore |
| cg12592365 | -0.22 | 0.0021 | 0.37 | 0.60 | 0.22 | RPTOR | Body | opensea |
| cg19318364 | -0.24 | 0.0012 | 0.63 | 0.87 | 0.24 | | IGR | opensea |
| cg19683494 | -0.25 | 0.0183 | 0.36 | 0.60 | 0.25 | | IGR | shore |
| cg15652532 | -0.27 | 0.0398 | 0.26 | 0.53 | 0.27 | LCLAT1 | TSS1500 | shore |
| cg22443212 | -0.44 | 0.0002 | 0.24 | 0.68 | 0.44 | RNF213 | Body | opensea |

**Supplemental Table 6.** Differentially methylated positions in Pre-LVAD vs. Post-LVAD (Reverse Remodeling) (adjusted p-value <0.05, abs (delta beta) > 10%)

| CpG Site | log FC | Adj. p-val | Pre-LVAD | Post-LVAD | Delta B | Gene | Feature | CGI |
|---|---|---|---|---|---|---|---|---|
| cg09559189 | 0.17 | 0.0001 | 0.37 | 0.20 | -0.17 | EBF2 | Body | shore |
| cg22280475 | 0.15 | 0.0033 | 0.34 | 0.19 | -0.15 | EBF2 | Body | island |
| cg14855519 | 0.14 | 0.0004 | 0.39 | 0.25 | -0.14 | EBF2 | Body | shore |
| cg18239431 | 0.14 | 0.0005 | 0.28 | 0.14 | -0.14 | EBF2 | Body | shore |
| cg03795776 | 0.14 | 0.0020 | 0.43 | 0.29 | -0.14 | BACH2 | Body | opensea |
| cg16310415 | 0.14 | 0.0001 | 0.33 | 0.19 | -0.14 | EBF2 | Body | shore |
| cg17902947 | 0.13 | 0.0090 | 0.63 | 0.50 | -0.13 | PPAP2B | Body | shelf |
| cg12536527 | 0.13 | 0.0022 | 0.32 | 0.19 | -0.13 | CACNA2D1 | Body | opensea |
| cg05748163 | 0.13 | 0.0030 | 0.32 | 0.19 | -0.13 | EBF2 | Body | shore |
| cg05996789 | 0.12 | 0.0057 | 0.74 | 0.62 | -0.12 | SLC25A4 | TSS1500 | shore |
| cg25892587 | 0.12 | 0.0060 | 0.42 | 0.30 | -0.12 | | IGR | opensea |
| cg04657684 | 0.12 | 0.0011 | 0.54 | 0.42 | -0.12 | ELMOD1 | Body | opensea |
| cg12149795 | 0.12 | 0.0003 | 0.27 | 0.16 | -0.12 | DIP2A | Body | shelf |
| cg16613240 | 0.12 | 0.0044 | 0.38 | 0.27 | -0.12 | | IGR | opensea |
| cg01386185 | 0.12 | 0.0002 | 0.74 | 0.63 | -0.12 | | IGR | opensea |
| cg00066750 | 0.12 | 0.0311 | 0.75 | 0.64 | -0.12 | HEY2 | Body | shore |
| cg24881607 | 0.11 | 0.0289 | 0.79 | 0.68 | -0.11 | HEY2 | Body | island |
| cg02371631 | 0.11 | 0.0010 | 0.55 | 0.44 | -0.11 | CDK15 | Body | opensea |
| cg17936488 | 0.11 | 0.0045 | 0.40 | 0.28 | -0.11 | FAM78A | 1stExon | shore |
| cg00730887 | 0.11 | 0.0008 | 0.27 | 0.16 | -0.11 | | IGR | opensea |
| cg14787880 | 0.11 | 0.0006 | 0.32 | 0.21 | -0.11 | BMPR2 | Body | shore |
| cg13425294 | 0.11 | 0.0058 | 0.31 | 0.19 | -0.11 | | IGR | opensea |
| cg15580052 | 0.11 | 0.0086 | 0.33 | 0.22 | -0.11 | B4GALNT3 | Body | opensea |
| cg09901201 | 0.11 | 0.0201 | 0.74 | 0.63 | -0.11 | SCP2 | 3'UTR | opensea |
| cg10378032 | 0.11 | 0.0043 | 0.59 | 0.48 | -0.11 | RAVER2 | Body | shelf |
| cg26874542 | 0.11 | 0.0031 | 0.55 | 0.44 | -0.11 | FGF18 | Body | shelf |
| cg03731740 | 0.11 | 0.0107 | 0.54 | 0.43 | -0.11 | YTHDF2 | TSS1500 | shore |
| cg03077077 | 0.11 | 0.0304 | 0.77 | 0.66 | -0.11 | TMEM155 | 5'UTR | shore |
| cg04492228 | 0.11 | 0.0000 | 0.57 | 0.47 | -0.11 | GATA3 | Body | shore |
| cg16411101 | 0.11 | 0.0020 | 0.79 | 0.68 | -0.11 | SSBP3 | Body | opensea |
| cg22783664 | 0.11 | 0.0014 | 0.24 | 0.13 | -0.11 | STX11 | 5'UTR | opensea |
| cg19343518 | 0.11 | 0.0047 | 0.39 | 0.28 | -0.11 | ARID1B | Body | opensea |
| cg01900030 | 0.11 | 0.0064 | 0.71 | 0.60 | -0.11 | CDK6 | Body | opensea |
| cg06734510 | 0.11 | 0.0138 | 0.78 | 0.67 | -0.11 | PLCL1 | Body | shelf |
| cg01190989 | 0.11 | 0.0025 | 0.45 | 0.34 | -0.11 | GPR125 | Body | opensea |
| cg06115614 | 0.11 | 0.0200 | 0.55 | 0.45 | -0.11 | GATA2 | Body | shore |
| cg05980083 | 0.11 | 0.0032 | 0.92 | 0.81 | -0.11 | PTPN3 | 5'UTR | opensea |
| cg11577329 | 0.10 | 0.0062 | 0.77 | 0.67 | -0.10 | SLC12A7 | Body | island |
| cg12251895 | 0.10 | 0.0004 | 0.24 | 0.13 | -0.10 | KLF7 | Body | opensea |

| | | | | | | | | |
|---|---|---|---|---|---|---|---|---|
| cg14362113 | 0.10 | 0.0032 | 0.70 | 0.59 | -0.10 | | IGR | opensea |
| cg11421073 | 0.10 | 0.0022 | 0.49 | 0.38 | -0.10 | ARL6IP5 | 3'UTR | opensea |
| cg27078890 | 0.10 | 0.0021 | 0.28 | 0.18 | -0.10 | ETS1 | TSS200 | opensea |
| cg17065262 | 0.10 | 0.0002 | 0.38 | 0.27 | -0.10 | MYO18A | 5'UTR | shore |
| cg08559364 | 0.10 | 0.0072 | 0.32 | 0.22 | -0.10 | VGLL4 | Body | opensea |
| cg05152903 | 0.10 | 0.0179 | 0.36 | 0.25 | -0.10 | CSDA | Body | shelf |
| cg05648629 | 0.10 | 0.0014 | 0.59 | 0.49 | -0.10 | ADCY9 | Body | shelf |
| cg00652274 | 0.10 | 0.0001 | 0.83 | 0.73 | -0.10 | LSAMP | Body | opensea |
| cg05634495 | 0.10 | 0.0044 | 0.36 | 0.26 | -0.10 | | IGR | opensea |
| cg07263393 | 0.10 | 0.0495 | 0.46 | 0.36 | -0.10 | GATA2 | Body | shore |
| cg12962542 | -0.10 | 0.0132 | 0.50 | 0.60 | 0.10 | MECOM | Body | opensea |
| cg09368075 | -0.10 | 0.0019 | 0.47 | 0.57 | 0.10 | | IGR | shore |
| cg02829601 | -0.10 | 0.0037 | 0.34 | 0.44 | 0.10 | SYTL3 | TSS200 | opensea |
| cg11552287 | -0.10 | 0.0167 | 0.37 | 0.47 | 0.10 | LANCL2 | TSS1500 | shore |
| cg12810626 | -0.10 | 0.0048 | 0.79 | 0.89 | 0.10 | LY9 | TSS1500 | opensea |
| cg23404248 | -0.10 | 0.0460 | 0.11 | 0.21 | 0.10 | | IGR | island |
| cg25391092 | -0.10 | 0.0364 | 0.58 | 0.68 | 0.10 | C10orf11 | Body | opensea |
| cg21350115 | -0.10 | 0.0044 | 0.26 | 0.36 | 0.10 | CALCRL | 1stExon | opensea |
| cg02719154 | -0.10 | 0.0400 | 0.17 | 0.27 | 0.10 | | IGR | shore |
| cg14511677 | -0.10 | 0.0157 | 0.39 | 0.49 | 0.10 | ST14 | Body | island |
| cg11036041 | -0.10 | 0.0098 | 0.31 | 0.41 | 0.10 | LIMCH1 | Body | shore |
| cg23557926 | -0.10 | 0.0243 | 0.30 | 0.40 | 0.10 | CFH | TSS200 | opensea |
| cg01552197 | -0.10 | 0.0240 | 0.46 | 0.56 | 0.10 | | IGR | opensea |
| cg22817650 | -0.10 | 0.0013 | 0.47 | 0.57 | 0.10 | CDC73 | Body | opensea |
| cg23044178 | -0.10 | 0.0232 | 0.52 | 0.62 | 0.10 | MICAL2 | 5'UTR | shelf |
| cg09255732 | -0.10 | 0.0107 | 0.37 | 0.47 | 0.10 | COL16A1 | TSS1500 | shore |
| cg25095814 | -0.10 | 0.0038 | 0.37 | 0.47 | 0.10 | CASP8 | TSS200 | opensea |
| cg01245224 | -0.10 | 0.0240 | 0.73 | 0.83 | 0.10 | | IGR | opensea |
| cg00960700 | -0.10 | 0.0075 | 0.16 | 0.26 | 0.10 | TBCD | TSS1500 | island |
| cg15521792 | -0.10 | 0.0000 | 0.53 | 0.63 | 0.10 | NUDT17 | TSS1500 | shore |
| cg08711175 | -0.10 | 0.0108 | 0.23 | 0.33 | 0.10 | NXPH4 | Body | shelf |
| cg11217865 | -0.10 | 0.0040 | 0.39 | 0.49 | 0.10 | NRG2 | 3'UTR | shore |
| cg16336556 | -0.10 | 0.0353 | 0.56 | 0.67 | 0.10 | LTBP1 | Body | opensea |
| cg09501687 | -0.10 | 0.0141 | 0.41 | 0.51 | 0.10 | C5orf62 | Body | opensea |
| cg00067758 | -0.10 | 0.0167 | 0.42 | 0.53 | 0.10 | | IGR | opensea |
| cg07113653 | -0.10 | 0.0204 | 0.74 | 0.85 | 0.10 | GFOD1 | Body | opensea |
| cg23806894 | -0.10 | 0.0174 | 0.13 | 0.23 | 0.10 | | IGR | shelf |
| cg05307752 | -0.10 | 0.0323 | 0.59 | 0.70 | 0.10 | ARHGAP15 | Body | opensea |
| cg26575450 | -0.10 | 0.0127 | 0.55 | 0.66 | 0.10 | | IGR | shore |
| cg21015470 | -0.10 | 0.0087 | 0.60 | 0.70 | 0.10 | OPCML | Body | opensea |
| cg01232969 | -0.10 | 0.0224 | 0.43 | 0.53 | 0.10 | | IGR | opensea |
| cg02500300 | -0.10 | 0.0040 | 0.18 | 0.28 | 0.10 | STOX2 | 1stExon | island |
| cg03926751 | -0.11 | 0.0138 | 0.65 | 0.75 | 0.11 | KLHL8 | 5'UTR | shore |

| | | | | | | | | |
|---|---|---|---|---|---|---|---|---|
| cg12590005 | -0.11 | 0.0044 | 0.18 | 0.28 | 0.11 | MAPK14 | Body | shore |
| cg23618477 | -0.11 | 0.0015 | 0.55 | 0.66 | 0.11 | | IGR | shelf |
| cg15174393 | -0.11 | 0.0256 | 0.37 | 0.48 | 0.11 | | IGR | opensea |
| cg13959207 | -0.11 | 0.0020 | 0.57 | 0.68 | 0.11 | ATP13A3 | TSS200 | opensea |
| cg25838968 | -0.11 | 0.0295 | 0.54 | 0.64 | 0.11 | PLXNA2 | Body | opensea |
| cg09233395 | -0.11 | 0.0162 | 0.43 | 0.53 | 0.11 | TRPS1 | 5'UTR | shore |
| cg01180552 | -0.11 | 0.0384 | 0.62 | 0.73 | 0.11 | | IGR | island |
| cg27520536 | -0.11 | 0.0280 | 0.52 | 0.63 | 0.11 | | IGR | island |
| cg04067806 | -0.11 | 0.0002 | 0.68 | 0.78 | 0.11 | | IGR | shore |
| cg09217350 | -0.11 | 0.0060 | 0.43 | 0.53 | 0.11 | | IGR | opensea |
| cg15931471 | -0.11 | 0.0097 | 0.54 | 0.65 | 0.11 | SMAD6 | Body | opensea |
| cg07034563 | -0.11 | 0.0091 | 0.69 | 0.80 | 0.11 | PDLIM7 | Body | shore |
| cg26877720 | -0.11 | 0.0017 | 0.64 | 0.75 | 0.11 | FAM107B | Body | shore |
| cg08750534 | -0.11 | 0.0007 | 0.24 | 0.35 | 0.11 | SPRED2 | Body | opensea |
| cg20765408 | -0.11 | 0.0088 | 0.61 | 0.71 | 0.11 | PARP4 | 5'UTR | shore |
| cg07777652 | -0.11 | 0.0039 | 0.71 | 0.82 | 0.11 | WRB | Body | shore |
| cg08141395 | -0.11 | 0.0060 | 0.18 | 0.29 | 0.11 | MAML2 | Body | opensea |
| cg09275704 | -0.11 | 0.0331 | 0.46 | 0.57 | 0.11 | | IGR | island |
| cg11509088 | -0.11 | 0.0000 | 0.53 | 0.63 | 0.11 | WDFY4 | Body | opensea |
| cg22184990 | -0.11 | 0.0041 | 0.39 | 0.50 | 0.11 | ARHGAP26 | Body | opensea |
| cg19541865 | -0.11 | 0.0107 | 0.69 | 0.80 | 0.11 | ACPP | TSS1500 | opensea |
| cg10229594 | -0.11 | 0.0051 | 0.25 | 0.36 | 0.11 | | IGR | opensea |
| cg24166450 | -0.11 | 0.0027 | 0.34 | 0.45 | 0.11 | | IGR | opensea |
| cg16867680 | -0.11 | 0.0047 | 0.30 | 0.41 | 0.11 | | IGR | opensea |
| cg21994818 | -0.11 | 0.0002 | 0.49 | 0.60 | 0.11 | | IGR | opensea |
| cg21594702 | -0.11 | 0.0235 | 0.36 | 0.47 | 0.11 | VCAN | 5'UTR | shore |
| cg23821329 | -0.11 | 0.0331 | 0.44 | 0.55 | 0.11 | VIM | TSS1500 | shore |
| cg27285720 | -0.11 | 0.0002 | 0.36 | 0.48 | 0.11 | GBP4 | TSS200 | opensea |
| cg14553740 | -0.11 | 0.0190 | 0.45 | 0.56 | 0.11 | FAM154A | Body | opensea |
| cg16496024 | -0.11 | 0.0023 | 0.35 | 0.46 | 0.11 | PCP4 | TSS1500 | opensea |
| cg17941330 | -0.11 | 0.0161 | 0.23 | 0.34 | 0.11 | GJA5 | TSS200 | opensea |
| cg09251959 | -0.11 | 0.0051 | 0.42 | 0.53 | 0.11 | COL16A1 | TSS1500 | shore |
| cg10699857 | -0.11 | 0.0376 | 0.11 | 0.22 | 0.11 | | IGR | island |
| cg11074047 | -0.12 | 0.0013 | 0.52 | 0.64 | 0.12 | NFATC2 | Body | shore |
| cg05033239 | -0.12 | 0.0079 | 0.36 | 0.47 | 0.12 | GPR98 | Body | opensea |
| cg04456219 | -0.12 | 0.0292 | 0.34 | 0.46 | 0.12 | | IGR | opensea |
| cg22305268 | -0.12 | 0.0008 | 0.54 | 0.66 | 0.12 | | IGR | shore |
| cg10308253 | -0.12 | 0.0050 | 0.30 | 0.42 | 0.12 | ZC3H12D | 5'UTR | opensea |
| cg01973456 | -0.12 | 0.0386 | 0.53 | 0.65 | 0.12 | | IGR | shelf |
| cg06088745 | -0.12 | 0.0298 | 0.47 | 0.60 | 0.12 | IRX3 | 3'UTR | shore |
| cg18142262 | -0.12 | 0.0116 | 0.18 | 0.31 | 0.12 | | IGR | opensea |
| cg26639076 | -0.13 | 0.0047 | 0.46 | 0.59 | 0.13 | RIF1 | 3'UTR | opensea |
| cg18693345 | -0.13 | 0.0144 | 0.25 | 0.38 | 0.13 | C5orf38 | Body | island |

| | | | | | | | | |
|---|---|---|---|---|---|---|---|---|
| cg03044684 | -0.13 | 0.0214 | 0.48 | 0.61 | 0.13 | HUNK | Body | shore |
| cg13064658 | -0.14 | 0.0057 | 0.07 | 0.21 | 0.14 | LPGAT1 | 5'UTR | island |
| cg15945235 | -0.15 | 0.0008 | 0.49 | 0.64 | 0.15 | ANKRD22 | 1stExon | opensea |
| cg07904452 | -0.15 | 0.0008 | 0.36 | 0.51 | 0.15 | | IGR | opensea |
| cg19611616 | -0.22 | 0.0293 | 0.03 | 0.25 | 0.22 | STK38L | 5'UTR | shore |

**Supplemental Table 7.** LVAD Responsive HF associated differentially methylated positions (adjusted p-value <0.05, abs (delta beta) > 10%)

| CpG Site | Non-Failing | Pre-LVAD | Post-LVAD | Delta HF | Delta LVAD | Gene | Feature | CGI |
|---|---|---|---|---|---|---|---|---|
| cg23821329 | 0.62 | 0.44 | 0.55 | -0.18 | 0.11 | VIM | TSS1500 | shore |
| cg02500300 | 0.35 | 0.18 | 0.28 | -0.18 | 0.10 | STOX2 | 1stExon | island |
| cg16336556 | 0.73 | 0.56 | 0.67 | -0.17 | 0.10 | LTBP1 | Body | opensea |
| cg22305268 | 0.69 | 0.54 | 0.66 | -0.15 | 0.12 | | IGR | shore |
| cg09233395 | 0.56 | 0.43 | 0.53 | -0.14 | 0.11 | TRPS1 | 5'UTR | shore |
| cg03044684 | 0.61 | 0.48 | 0.61 | -0.14 | 0.13 | HUNK | Body | shore |
| cg26877720 | 0.77 | 0.64 | 0.75 | -0.13 | 0.11 | FAM107B | Body | shore |
| cg27520536 | 0.65 | 0.52 | 0.63 | -0.13 | 0.11 | | IGR | island |
| cg09251959 | 0.55 | 0.42 | 0.53 | -0.13 | 0.11 | COL16A1 | TSS1500 | shore |
| cg26575450 | 0.68 | 0.55 | 0.66 | -0.13 | 0.10 | | IGR | shore |
| cg14553740 | 0.58 | 0.45 | 0.56 | -0.13 | 0.11 | FAM154A | Body | opensea |
| cg15174393 | 0.49 | 0.37 | 0.48 | -0.12 | 0.11 | | IGR | opensea |
| cg25838968 | 0.65 | 0.54 | 0.64 | -0.12 | 0.11 | PLXNA2 | Body | opensea |
| cg04456219 | 0.46 | 0.34 | 0.46 | -0.12 | 0.12 | | IGR | opensea |
| cg15931471 | 0.66 | 0.54 | 0.65 | -0.12 | 0.11 | SMAD6 | Body | opensea |
| cg07034563 | 0.80 | 0.69 | 0.80 | -0.11 | 0.11 | PDLIM7 | Body | shore |
| cg21994818 | 0.60 | 0.49 | 0.60 | -0.11 | 0.11 | | IGR | opensea |
| cg15945235 | 0.60 | 0.49 | 0.64 | -0.11 | 0.15 | ANKRD22 | 1stExon | opensea |
| cg24166450 | 0.45 | 0.34 | 0.45 | -0.11 | 0.11 | | IGR | opensea |
| cg20765408 | 0.71 | 0.61 | 0.71 | -0.10 | 0.11 | PARP4 | 5'UTR | shore |
| cg03926751 | 0.75 | 0.65 | 0.75 | -0.10 | 0.11 | KLHL8 | 5'UTR | shore |
| cg17936488 | 0.29 | 0.40 | 0.28 | 0.11 | -0.11 | FAM78A | 1stExon | shore |
| cg04657684 | 0.43 | 0.54 | 0.42 | 0.11 | -0.12 | ELMOD1 | Body | opensea |
| cg15580052 | 0.21 | 0.33 | 0.22 | 0.12 | -0.11 | B4GALNT3 | Body | opensea |
| cg08559364 | 0.20 | 0.32 | 0.22 | 0.12 | -0.10 | VGLL4 | Body | opensea |
| cg16310415 | 0.21 | 0.33 | 0.19 | 0.12 | -0.14 | EBF2 | Body | shore |
| cg14855519 | 0.26 | 0.39 | 0.25 | 0.12 | -0.14 | EBF2 | Body | shore |
| cg18239431 | 0.16 | 0.28 | 0.14 | 0.13 | -0.14 | EBF2 | Body | shore |
| cg05748163 | 0.18 | 0.32 | 0.19 | 0.14 | -0.13 | EBF2 | Body | shore |
| cg01900030 | 0.57 | 0.71 | 0.60 | 0.14 | -0.11 | CDK6 | Body | opensea |
| cg01190989 | 0.30 | 0.45 | 0.34 | 0.15 | -0.11 | GPR125 | Body | opensea |
| cg04492228 | 0.42 | 0.57 | 0.47 | 0.15 | -0.11 | GATA3 | Body | shore |
| cg03731740 | 0.39 | 0.54 | 0.43 | 0.15 | -0.11 | YTHDF2 | TSS1500 | shore |
| cg22280475 | 0.18 | 0.34 | 0.19 | 0.16 | -0.15 | EBF2 | Body | island |
| cg09559189 | 0.22 | 0.37 | 0.20 | 0.16 | -0.17 | EBF2 | Body | shore |

**Supplemental Table 8.** DNA Methylation vs. Gene Expression Correlation of Common HF Differentially Methylated Position in ICM and NICM

| DNA | mRNA | Gene | Log FC | AveExpr | t | P value | Adj. p | B | Delta B |
|---|---|---|---|---|---|---|---|---|---|
| Hypo-methylated | Up-regulated | HTRA1 | 1.014 | 6.316 | 7.864 | 0.000 | 0.000 | 14.911 | -0.132 |
| Hypo-methylated | Up-regulated | FAM65B | 0.822 | 2.262 | 4.827 | 0.000 | 0.000 | 3.198 | -0.161 |
| Hypo-methylated | Up-regulated | FBXO16 | 0.726 | 2.274 | 5.119 | 0.000 | 0.000 | 4.248 | -0.122 |
| Hypo-methylated | Up-regulated | EFCAB13 | 0.483 | 1.259 | 3.059 | 0.003 | 0.010 | -2.385 | -0.198 |
| Hypo-methylated | Up-regulated | COL18A1 | 0.467 | 5.685 | 1.783 | 0.079 | 0.142 | -5.207 | -0.136 |
| Hypo-methylated | Up-regulated | UNC45A | 0.460 | 4.032 | 5.407 | 0.000 | 0.000 | 5.305 | -0.119 |
| Hypo-methylated | Up-regulated | KALRN | 0.439 | 5.233 | 3.638 | 0.001 | 0.002 | -0.732 | -0.140 |
| Hypo-methylated | Up-regulated | AKAP13 | 0.430 | 7.326 | 4.383 | 0.000 | 0.000 | 1.659 | -0.124 |
| Hypo-methylated | Up-regulated | RPTOR | 0.419 | 3.359 | 4.728 | 0.000 | 0.000 | 2.849 | -0.205 |
| Hypo-methylated | Up-regulated | KIAA0922 | 0.349 | 3.258 | 3.184 | 0.002 | 0.007 | -2.045 | -0.152 |
| Hypo-methylated | Up-regulated | HDAC9 | 0.347 | 3.486 | 2.322 | 0.023 | 0.052 | -4.168 | -0.140 |
| Hypo-methylated | Up-regulated | PLXNA2 | 0.345 | 3.962 | 2.960 | 0.004 | 0.013 | -2.646 | -0.174 |
| Hypo-methylated | Up-regulated | BRE | 0.343 | 5.701 | 5.139 | 0.000 | 0.000 | 4.319 | -0.156 |
| Hypo-methylated | Up-regulated | MAP3K14 | 0.261 | 3.396 | 2.397 | 0.019 | 0.045 | -4.005 | -0.138 |
| Hypo-methylated | Down-regulated | PRDM16 | -0.285 | 2.458 | -2.704 | 0.009 | 0.023 | -3.292 | -0.133 |
| Hypo-methylated | Down-regulated | STXBP5 | -0.286 | 2.405 | -2.703 | 0.009 | 0.023 | -3.294 | -0.147 |
| Hypo-methylated | Down-regulated | BATF | -0.291 | 1.178 | -1.255 | 0.214 | 0.316 | -5.983 | -0.121 |
| Hypo-methylated | Down-regulated | PLD6 | -0.330 | 2.603 | -2.744 | 0.008 | 0.021 | -3.195 | -0.141 |
| Hypo-methylated | Down-regulated | SYN2 | -0.339 | 1.276 | -1.494 | 0.140 | 0.225 | -5.663 | -0.133 |
| Hypo-methylated | Down-regulated | PDE4D | -0.352 | 3.379 | -2.235 | 0.029 | 0.062 | -4.352 | -0.127 |
| Hypo-methylated | Down-regulated | ANKRD6 | -0.372 | 3.635 | -2.908 | 0.005 | 0.014 | -2.781 | -0.142 |
| Hypo-methylated | Down-regulated | FRK | -0.417 | 1.906 | -3.230 | 0.002 | 0.007 | -1.919 | -0.106 |
| Hypo-methylated | Down-regulated | ABLIM1 | -0.427 | 8.671 | -5.452 | 0.000 | 0.000 | 5.474 | -0.191 |
| Hypo-methylated | Down-regulated | SOD2 | -0.578 | 9.177 | -5.893 | 0.000 | 0.000 | 7.141 | -0.118 |
| Hypo-methylated | Down-regulated | PARD3 | -0.699 | 4.697 | -7.768 | 0.000 | 0.000 | 14.526 | -0.123 |
| Hypo-methylated | Down-regulated | PALLD | -0.998 | 8.631 | -9.168 | 0.000 | 0.000 | 20.098 | -0.162 |
| Hypo-methylated | Down-regulated | RARRES1 | -1.755 | 4.029 | -4.402 | 0.000 | 0.000 | 1.721 | -0.149 |
| Hyper-methylated | Up-regulated | PCDHGA2 | 0.740 | 2.127 | 6.295 | 0.000 | 0.000 | 8.695 | 0.148 |
| Hyper-methylated | Up-regulated | ATOH8 | 0.701 | 4.693 | 4.155 | 0.000 | 0.001 | 0.897 | 0.109 |
| Hyper-methylated | Up-regulated | SOX9 | 0.617 | 3.453 | 3.977 | 0.000 | 0.001 | 0.324 | 0.123 |
| Hyper-methylated | Up-regulated | CPT1C | 0.495 | 2.080 | 3.135 | 0.003 | 0.008 | -2.180 | 0.156 |
| Hyper-methylated | Up-regulated | RAB11FIP1 | 0.417 | 3.402 | 2.219 | 0.030 | 0.064 | -4.386 | 0.122 |
| Hyper-methylated | Up-regulated | PRRT1 | 0.270 | 2.535 | 2.233 | 0.029 | 0.062 | -4.355 | 0.108 |
| Hyper-methylated | Down-regulated | TBX3 | -0.537 | 3.790 | -3.840 | 0.000 | 0.001 | -0.111 | 0.123 |

**Supplemental Table 9.** DNA Methylated CpG Sites that are located within or near (<10 kb) human long-non-coding RNA (lncRNA) genes

| Heart Failure DMPs | lncRNA Name | lncRNA Type | DNA Strand | Genomic Location | Cardiac Expression GTEX (TPM) |
|---|---|---|---|---|---|
| **Ischemic and Non-Ischemic Cardiomyopathy DMPs** | | | | | |
| cg23099839 | GS1-57L11.1 | Intergenic | + | chr8:2584858-2680004 | 0.000 |
| cg23099839 | RP11-134O21.1 | Intergenic | - | chr8:2523591-2585991 | 0.000 |
| cg27149179 | CTD-2005H7.2 | Intergenic | + | chr11:86438397-86476307 | 0.078 |
| cg07986257 | RP3-404K8.2 | Antisense | + | chr6:22260653-22318027 | - |
| cg21727223 | LINC01482 | Intergenic | + | chr17:66587980-66746693 | 0.001 |
| cg21727223 | RP11-118B18.2 | Intergenic | - | chr17:66789690-66793963 | 0.000 |
| cg04926881 | CTB-113D17.1 | Antisense | + | chr7:29019583-29052983 | 0.198 |
| cg04926881 | AC005162.5 | Antisense | + | chr7:29026644-29028515 | 0.000 |
| cg01195564 | LINC01331 | Intergenic | - | chr5:73407515-73832649 | 0.000 |
| cg26203572 | LINC00525 | Intergenic | + | chr7:47801074-47806370 | 0.168 |
| cg02351277 | LINC00578 | Intergenic | + | chr3:177159709-177469882 | 0.000 |
| cg02351277 | RP11-114M1.1 | Intergenic | + | chr3:177401415-177409038 | 0.000 |
| cg24475182 | RP11-134O21.1 | Intergenic | - | chr8:2523591-2585991 | 0.000 |
| cg24475182 | GS1-57L11.1 | Intergenic | + | chr8:2584858-2680004 | 0.000 |
| cg07506560 | AC093802.1 | Intergenic | + | chr2:240684554-240724577 | 0.000 |
| cg13052638 | RP11-534L20.5 | Intergenic | + | chr1:206677281-206677789 | 0.000 |
| cg04057161 | RP11-317M11.1 | Sense | + | chr4:54525375-54603323 | 0.000 |
| cg01412419 | RP4-536B24.3 | Intergenic | - | chr16:87813740-87840082 | 0.000 |
| cg01412419 | RP4-536B24.2 | Antisense | + | chr16:87870138-87871269 | 0.000 |
| cg08889114 | LMCD1-AS1 | Antisense | - | chr2:21444047-22193831 | 0.054 |
| cg08889114 | AC034187.2 | Intergenic | - | chr3:8615412-8634810 | 0.000 |
| cg14111334 | RP11-626H12.1 | Intergenic | + | chr11:69831982-69861921 | 0.070 |
| cg14111334 | RP11-626H12.2 | Intergenic | - | chr11:69860964-69867165 | 0.113 |
| cg14387312 | LINC01618 | Intergenic | + | chr4:53578561-53732988 | 0.000 |
| cg19590421 | RP11-218E20.2 | Antisense | + | chr14:51314840-51332377 | 0.000 |
| cg25720795 | HOTTIP | Antisense | + | chr7:27241461-27246878 | 0.000 |
| cg01535205 | LINC00880 | Intergenic | - | chr3:156799456-156840793 | 0.031 |
| cg01535205 | LINC00881 | Intergenic | + | chr3:156807670-156818924 | 35.17 |
| cg16200531 | CCNT2-AS1 | Antisense | - | chr2:135493034-135676240 | 0.444 |
| cg03329019 | HLX-AS1 | Antisense | - | chr1:221006105-221053482 | 0.053 |
| cg19657945 | MINCR | Antisense | - | chr8:144362336-144363830 | 2.42 |
| cg13283845 | MINCR | Antisense | - | chr8:144362336-144363830 | 2.42 |
| cg26536949 | AC108004.2 | Intergenic | - | chr17:33615-41378 | 0.068 |
| cg05528899 | AC108004.2 | Intergenic | - | chr17:33615-41378 | 0.068 |
| **LVAD Responsive HF DMPs** | | | | | |
| cg26575450 | LINC02579 | Intergenic | + | chr2:64834446-64843616 | 0.000 |
| cg21994818 | C8orf37-AS1 | Antisense | + | chr8:96281064-96822371 | 0.000 |

**Supplemental Table 10.** Top 100 mRNAs that are regulated by LINC00881 plasmid overexpression in the beating human iPS cell derived cardiomyocytes by Deseq2

| Gene ID | Base | Log2 FC | lfc SE | stat | p value | p adj |
|---|---|---|---|---|---|---|
| LINC00881 | 3323.2 | 2.07 | 0.18 | 11.38 | 0.0000 | 0.0000 |
| MYH6 | 58976.6 | 0.35 | 0.07 | 4.98 | 0.0000 | 0.0105 |
| FXYD6 | 608.8 | 0.41 | 0.08 | 4.84 | 0.0000 | 0.0143 |
| DNTTIP2 | 1686.7 | -0.33 | 0.07 | -4.66 | 0.0000 | 0.0253 |
| ARL6IP1 | 1427.0 | -0.29 | 0.06 | -4.55 | 0.0000 | 0.0350 |
| EIF2S2 | 3515.4 | -0.24 | 0.05 | -4.47 | 0.0000 | 0.0428 |
| CACNA1C | 1375.7 | 0.36 | 0.08 | 4.39 | 0.0000 | 0.0461 |
| HSPG2 | 1642.4 | 0.29 | 0.07 | 4.36 | 0.0000 | 0.0461 |
| SYF2 | 532.0 | -0.36 | 0.08 | -4.36 | 0.0000 | 0.0461 |
| POLR1F | 592.9 | -0.38 | 0.09 | -4.21 | 0.0000 | 0.0797 |
| RIOK2 | 1058.1 | -0.30 | 0.07 | -4.19 | 0.0000 | 0.0797 |
| ZNF146 | 1964.0 | -0.25 | 0.06 | -4.16 | 0.0000 | 0.0797 |
| MT-RNR2 | 101974.6 | -0.29 | 0.07 | -4.16 | 0.0000 | 0.0797 |
| NFE2L2 | 1280.6 | -0.26 | 0.06 | -4.14 | 0.0000 | 0.0808 |
| MYBPC3 | 12012.0 | 0.35 | 0.09 | 4.08 | 0.0000 | 0.0966 |
| CEBPZ | 1638.6 | -0.23 | 0.06 | -4.01 | 0.0001 | 0.1251 |
| CALR | 3952.5 | 0.31 | 0.08 | 3.99 | 0.0001 | 0.1276 |
| NID2 | 3748.1 | 0.30 | 0.07 | 3.97 | 0.0001 | 0.1276 |
| LDLR | 1381.0 | 0.32 | 0.08 | 3.95 | 0.0001 | 0.1310 |
| A2M | 8017.2 | 0.36 | 0.09 | 3.94 | 0.0001 | 0.1310 |
| DYSF | 1200.8 | 0.28 | 0.07 | 3.92 | 0.0001 | 0.1386 |
| CNN2 | 1355.3 | 0.30 | 0.08 | 3.88 | 0.0001 | 0.1474 |
| UNC5B | 1389.8 | 0.26 | 0.07 | 3.88 | 0.0001 | 0.1474 |
| RGS4 | 696.6 | -0.54 | 0.14 | -3.87 | 0.0001 | 0.1476 |
| ACACB | 981.4 | 0.26 | 0.07 | 3.85 | 0.0001 | 0.1476 |
| EMC2 | 728.0 | -0.30 | 0.08 | -3.85 | 0.0001 | 0.1476 |
| TMEM167A | 4189.5 | -0.25 | 0.07 | -3.77 | 0.0002 | 0.1808 |
| CACNA1D | 591.3 | 0.35 | 0.09 | 3.76 | 0.0002 | 0.1808 |
| ZFAND5 | 3454.4 | -0.23 | 0.06 | -3.75 | 0.0002 | 0.1808 |
| VCAN | 4920.5 | 0.21 | 0.06 | 3.75 | 0.0002 | 0.1808 |
| PHAX | 600.5 | -0.28 | 0.08 | -3.75 | 0.0002 | 0.1808 |
| SVIL | 9311.4 | 0.19 | 0.05 | 3.74 | 0.0002 | 0.1808 |
| SEC62 | 2171.8 | -0.31 | 0.08 | -3.74 | 0.0002 | 0.1808 |
| TM4SF1 | 118.4 | -0.54 | 0.15 | -3.72 | 0.0002 | 0.1888 |
| CDC42EP3 | 3137.1 | -0.31 | 0.08 | -3.71 | 0.0002 | 0.1902 |
| RPL14 | 3635.7 | -0.25 | 0.07 | -3.70 | 0.0002 | 0.1954 |
| COL6A1 | 751.1 | 0.44 | 0.12 | 3.68 | 0.0002 | 0.2014 |
| SYNE2 | 814.5 | 0.25 | 0.07 | 3.67 | 0.0002 | 0.2014 |
| OARD1 | 618.8 | -0.30 | 0.08 | -3.67 | 0.0002 | 0.2014 |

| | | | | | | |
|---|---|---|---|---|---|---|
| CIR1 | 624.8 | -0.29 | 0.08 | -3.66 | 0.0002 | 0.2016 |
| ESF1 | 641.2 | -0.32 | 0.09 | -3.65 | 0.0003 | 0.2083 |
| GNAI3 | 1217.0 | -0.24 | 0.07 | -3.63 | 0.0003 | 0.2146 |
| MT-CO2 | 63460.8 | -0.22 | 0.06 | -3.63 | 0.0003 | 0.2146 |
| MT-RNR1 | 13798.2 | -0.24 | 0.07 | -3.62 | 0.0003 | 0.2146 |
| C16orf72 | 1744.5 | -0.25 | 0.07 | -3.61 | 0.0003 | 0.2199 |
| PYROXD2 | 293.7 | 0.42 | 0.12 | 3.60 | 0.0003 | 0.2199 |
| ZNF800 | 265.3 | -0.44 | 0.12 | -3.60 | 0.0003 | 0.2199 |
| BNIP2 | 1981.3 | -0.24 | 0.07 | -3.59 | 0.0003 | 0.2249 |
| CFAP97 | 1046.6 | -0.28 | 0.08 | -3.57 | 0.0004 | 0.2249 |
| GAA | 288.3 | 0.39 | 0.11 | 3.57 | 0.0004 | 0.2249 |
| ACTA2 | 4114.2 | 0.33 | 0.09 | 3.56 | 0.0004 | 0.2249 |
| SLK | 3830.9 | -0.24 | 0.07 | -3.56 | 0.0004 | 0.2249 |
| PXDN | 1313.2 | 0.22 | 0.06 | 3.56 | 0.0004 | 0.2249 |
| PDIA4 | 1199.4 | 0.31 | 0.09 | 3.55 | 0.0004 | 0.2249 |
| TAF1D | 747.3 | -0.33 | 0.09 | -3.54 | 0.0004 | 0.2249 |
| ZNF281 | 1022.4 | -0.28 | 0.08 | -3.53 | 0.0004 | 0.2249 |
| LSS | 897.1 | 0.37 | 0.11 | 3.53 | 0.0004 | 0.2249 |
| EIF4A2 | 7267.6 | -0.18 | 0.05 | -3.52 | 0.0004 | 0.2249 |
| PIK3CA | 1201.6 | -0.30 | 0.08 | -3.52 | 0.0004 | 0.2249 |
| TMEM167B | 994.4 | -0.25 | 0.07 | -3.52 | 0.0004 | 0.2249 |
| PLEKHA6 | 575.9 | 0.32 | 0.09 | 3.52 | 0.0004 | 0.2249 |
| DNAJC2 | 660.2 | -0.30 | 0.09 | -3.51 | 0.0004 | 0.2249 |
| PPP4R2 | 2391.2 | -0.28 | 0.08 | -3.51 | 0.0004 | 0.2249 |
| RAD17 | 1171.7 | -0.24 | 0.07 | -3.51 | 0.0005 | 0.2249 |
| CAMK2A | 243.5 | 0.37 | 0.11 | 3.50 | 0.0005 | 0.2249 |
| ADGRB2 | 727.1 | 0.39 | 0.11 | 3.50 | 0.0005 | 0.2249 |
| AFTPH | 449.7 | -0.31 | 0.09 | -3.50 | 0.0005 | 0.2249 |
| SH3GLB2 | 417.9 | 0.34 | 0.10 | 3.49 | 0.0005 | 0.2339 |
| RAB22A | 499.2 | -0.29 | 0.08 | -3.48 | 0.0005 | 0.2339 |
| KCNH7 | 131.9 | 0.50 | 0.14 | 3.48 | 0.0005 | 0.2339 |
| SLU7 | 1103.9 | -0.26 | 0.08 | -3.46 | 0.0005 | 0.2339 |
| RPL23A | 3334.9 | -0.25 | 0.07 | -3.46 | 0.0005 | 0.2339 |
| GREB1 | 1164.5 | 0.22 | 0.06 | 3.46 | 0.0005 | 0.2339 |
| ADAM11 | 321.3 | 0.40 | 0.12 | 3.46 | 0.0005 | 0.2339 |
| KLHL41 | 1289.4 | -0.24 | 0.07 | -3.46 | 0.0005 | 0.2339 |
| SCUBE3 | 1507.6 | 0.24 | 0.07 | 3.46 | 0.0005 | 0.2339 |
| NMD3 | 1603.0 | -0.24 | 0.07 | -3.44 | 0.0006 | 0.2361 |
| UBE3A | 3478.2 | -0.21 | 0.06 | -3.44 | 0.0006 | 0.2361 |
| KLHL9 | 1370.4 | -0.24 | 0.07 | -3.44 | 0.0006 | 0.2361 |
| POT1 | 467.7 | -0.28 | 0.08 | -3.44 | 0.0006 | 0.2361 |
| DMAP1 | 306.1 | 0.32 | 0.09 | 3.43 | 0.0006 | 0.2361 |
| MYH7B | 782.1 | 0.32 | 0.09 | 3.43 | 0.0006 | 0.2361 |

| | | | | | | |
|---|---|---|---|---|---|---|
| UBE2V2 | 1054.1 | -0.27 | 0.08 | -3.42 | 0.0006 | 0.2430 |
| OBI1 | 1128.3 | -0.29 | 0.08 | -3.42 | 0.0006 | 0.2430 |
| CCT8 | 4009.7 | -0.22 | 0.07 | -3.42 | 0.0006 | 0.2430 |
| RAD23B | 3510.6 | -0.23 | 0.07 | -3.41 | 0.0006 | 0.2434 |
| LAMB2 | 3760.3 | 0.31 | 0.09 | 3.41 | 0.0007 | 0.2456 |
| AP5M1 | 1967.0 | -0.24 | 0.07 | -3.39 | 0.0007 | 0.2504 |
| SERP1 | 620.1 | -0.28 | 0.08 | -3.39 | 0.0007 | 0.2504 |
| TENM4 | 687.6 | 0.27 | 0.08 | 3.39 | 0.0007 | 0.2504 |
| FAM107B | 271.4 | -0.33 | 0.10 | -3.39 | 0.0007 | 0.2513 |
| ZNF639 | 787.2 | -0.26 | 0.08 | -3.38 | 0.0007 | 0.2526 |
| RBAK | 821.8 | -0.31 | 0.09 | -3.38 | 0.0007 | 0.2554 |
| COL5A1 | 352.3 | 0.33 | 0.10 | 3.37 | 0.0007 | 0.2566 |
| CHD1 | 990.2 | -0.25 | 0.07 | -3.35 | 0.0008 | 0.2802 |
| MSANTD4 | 1268.7 | -0.26 | 0.08 | -3.34 | 0.0009 | 0.2837 |
| LRP1 | 1043.1 | 0.30 | 0.09 | 3.33 | 0.0009 | 0.2837 |
| MYOM2 | 107.3 | 0.51 | 0.15 | 3.33 | 0.0009 | 0.2837 |
| EMILIN2 | 3342.9 | 0.20 | 0.06 | 3.33 | 0.0009 | 0.2837 |
| ATP2B4 | 3356.0 | 0.23 | 0.07 | 3.33 | 0.0009 | 0.2837 |

**Supplemental Table 11.** Top 100 mRNAs that are regulated by LINC00881 GapmeR knockdown in the beating human iPS cell derived cardiomyocytes by Deseq2

| Gene ID | Base | log2 FC | lfc SE | stat | p value | p adj |
|---|---|---|---|---|---|---|
| DTNA | 1940.6 | -1.08 | 0.17 | -6.19 | 0.0000 | 0.0000 |
| LINC00881 | 1907.5 | -2.27 | 0.39 | -5.84 | 0.0000 | 0.0000 |
| ERRFI1 | 605.7 | 1.21 | 0.23 | 5.29 | 0.0000 | 0.0007 |
| THADA | 523.8 | -1.32 | 0.25 | -5.26 | 0.0000 | 0.0007 |
| FOSL1 | 75.1 | 2.10 | 0.41 | 5.11 | 0.0000 | 0.0012 |
| F2RL2 | 17.3 | 6.88 | 1.36 | 5.07 | 0.0000 | 0.0012 |
| FAM53C | 3717.1 | 0.71 | 0.14 | 5.03 | 0.0000 | 0.0013 |
| VPS45 | 2903.3 | -0.89 | 0.18 | -4.98 | 0.0000 | 0.0015 |
| RP11-419I17.1 | 167.8 | -1.17 | 0.24 | -4.80 | 0.0000 | 0.0032 |
| CYB561 | 598.2 | 1.38 | 0.29 | 4.78 | 0.0000 | 0.0032 |
| CCNG1 | 5904.2 | 0.62 | 0.13 | 4.76 | 0.0000 | 0.0032 |
| NUS1 | 513.9 | 0.76 | 0.16 | 4.71 | 0.0000 | 0.0034 |
| FARP1 | 859.5 | -0.85 | 0.18 | -4.70 | 0.0000 | 0.0034 |
| GALNT11 | 1152.6 | -0.61 | 0.13 | -4.70 | 0.0000 | 0.0034 |
| RSU1 | 3123.8 | -0.82 | 0.18 | -4.65 | 0.0000 | 0.0041 |
| TBCD | 875.3 | -1.04 | 0.22 | -4.62 | 0.0000 | 0.0042 |
| NOCT | 172.8 | 1.04 | 0.23 | 4.62 | 0.0000 | 0.0042 |
| KLF12 | 667.5 | -1.04 | 0.23 | -4.60 | 0.0000 | 0.0043 |
| MYLIP | 1137.7 | 1.04 | 0.23 | 4.58 | 0.0000 | 0.0045 |
| KIF26B | 1749.9 | -1.01 | 0.22 | -4.53 | 0.0000 | 0.0048 |
| PMAIP1 | 386.9 | 1.34 | 0.30 | 4.53 | 0.0000 | 0.0048 |
| CDH4 | 136.8 | -1.51 | 0.33 | -4.53 | 0.0000 | 0.0048 |
| PLCXD2 | 63.3 | 2.00 | 0.44 | 4.53 | 0.0000 | 0.0048 |
| SLC16A14 | 186.9 | 1.84 | 0.41 | 4.50 | 0.0000 | 0.0051 |
| RP11-556O5.7 | 50.1 | -2.55 | 0.57 | -4.47 | 0.0000 | 0.0058 |
| TRAPPC9 | 349.1 | -1.08 | 0.24 | -4.44 | 0.0000 | 0.0064 |
| CDC42EP1 | 325.5 | 1.53 | 0.35 | 4.43 | 0.0000 | 0.0064 |
| ATF6 | 1325.9 | -0.81 | 0.18 | -4.42 | 0.0000 | 0.0064 |
| ADAM23 | 553.6 | -1.76 | 0.40 | -4.40 | 0.0000 | 0.0070 |
| FBXW8 | 845.3 | -0.71 | 0.16 | -4.39 | 0.0000 | 0.0070 |
| SLC12A7 | 1981.0 | -0.81 | 0.19 | -4.38 | 0.0000 | 0.0070 |
| DOCK1 | 490.8 | -1.15 | 0.26 | -4.37 | 0.0000 | 0.0070 |
| NDUFV2 | 56.8 | 1.68 | 0.38 | 4.37 | 0.0000 | 0.0070 |
| KLF10 | 842.4 | 2.39 | 0.55 | 4.36 | 0.0000 | 0.0072 |
| SIX4 | 328.9 | 0.98 | 0.22 | 4.35 | 0.0000 | 0.0072 |
| WDR59 | 471.3 | -1.43 | 0.33 | -4.34 | 0.0000 | 0.0072 |
| DUSP1 | 1488.4 | 0.74 | 0.17 | 4.33 | 0.0000 | 0.0073 |
| PRKCI | 1235.1 | 0.63 | 0.14 | 4.33 | 0.0000 | 0.0073 |
| SLC3A2 | 6297.2 | 0.79 | 0.18 | 4.32 | 0.0000 | 0.0073 |

| | | | | | | |
|---|---|---|---|---|---|---|
| CRADD | 661.9 | -1.27 | 0.30 | -4.29 | 0.0000 | 0.0082 |
| ACTBL2 | 15.4 | 7.44 | 1.74 | 4.29 | 0.0000 | 0.0082 |
| BCLAF1 | 5568.3 | 0.73 | 0.17 | 4.28 | 0.0000 | 0.0083 |
| FOXH1 | 175.8 | 2.99 | 0.70 | 4.27 | 0.0000 | 0.0084 |
| LZTS3 | 352.8 | 1.01 | 0.24 | 4.26 | 0.0000 | 0.0084 |
| GPR3 | 126.3 | 1.63 | 0.38 | 4.26 | 0.0000 | 0.0084 |
| PIK3C3 | 937.4 | -0.96 | 0.23 | -4.24 | 0.0000 | 0.0089 |
| FBXO2 | 1219.5 | 0.61 | 0.14 | 4.24 | 0.0000 | 0.0089 |
| CYB5RL | 300.8 | -0.88 | 0.21 | -4.23 | 0.0000 | 0.0089 |
| CAMLG | 1914.9 | 0.59 | 0.14 | 4.22 | 0.0000 | 0.0094 |
| FSTL3 | 573.8 | 1.07 | 0.25 | 4.21 | 0.0000 | 0.0096 |
| RCC2 | 1164.4 | 0.80 | 0.19 | 4.19 | 0.0000 | 0.0100 |
| EDA | 24.8 | -3.68 | 0.88 | -4.19 | 0.0000 | 0.0100 |
| RALYL | 99.1 | -1.78 | 0.43 | -4.18 | 0.0000 | 0.0101 |
| CHST9 | 18.1 | -4.99 | 1.20 | -4.16 | 0.0000 | 0.0110 |
| MARCHF11 | 387.0 | -1.65 | 0.40 | -4.15 | 0.0000 | 0.0113 |
| ACVR2B | 1213.5 | 0.65 | 0.16 | 4.14 | 0.0000 | 0.0115 |
| RP11-110G21.1 | 254.9 | -0.82 | 0.20 | -4.12 | 0.0000 | 0.0124 |
| SERP1 | 2751.2 | 0.62 | 0.15 | 4.11 | 0.0000 | 0.0125 |
| JUND | 235.7 | 1.16 | 0.28 | 4.09 | 0.0000 | 0.0135 |
| GPER1 | 55.9 | 1.72 | 0.42 | 4.09 | 0.0000 | 0.0135 |
| MAFF | 84.7 | 1.28 | 0.31 | 4.07 | 0.0000 | 0.0139 |
| DISP1 | 336.4 | -0.88 | 0.22 | -4.06 | 0.0000 | 0.0139 |
| TSPYL2 | 1481.3 | 1.42 | 0.35 | 4.06 | 0.0000 | 0.0139 |
| MAPKAP1 | 2286.2 | -0.72 | 0.18 | -4.06 | 0.0000 | 0.0139 |
| RP11-74J13.8 | 39.7 | -1.97 | 0.49 | -4.06 | 0.0000 | 0.0139 |
| DYM | 1056.3 | -0.68 | 0.17 | -4.06 | 0.0000 | 0.0139 |
| AP4S1 | 266.8 | -0.91 | 0.23 | -4.04 | 0.0001 | 0.0148 |
| YOD1 | 947.7 | 0.59 | 0.15 | 4.04 | 0.0001 | 0.0148 |
| CACNA1C | 898.0 | -1.17 | 0.29 | -4.03 | 0.0001 | 0.0148 |
| ERI3 | 2031.7 | -0.70 | 0.17 | -4.02 | 0.0001 | 0.0155 |
| PDLIM3 | 955.6 | 1.14 | 0.28 | 4.02 | 0.0001 | 0.0155 |
| LINC00638 | 47.5 | -2.24 | 0.56 | -4.01 | 0.0001 | 0.0159 |
| IMPG2 | 44.7 | -5.04 | 1.26 | -3.99 | 0.0001 | 0.0164 |
| FTL | 74039.0 | 1.26 | 0.31 | 3.99 | 0.0001 | 0.0165 |
| P3H1 | 749.0 | 0.92 | 0.23 | 3.99 | 0.0001 | 0.0166 |
| PHKB | 4104.6 | -0.60 | 0.15 | -3.98 | 0.0001 | 0.0171 |
| SOX9 | 141.1 | 1.44 | 0.36 | 3.97 | 0.0001 | 0.0173 |
| CIB1 | 1466.3 | 0.57 | 0.14 | 3.97 | 0.0001 | 0.0174 |
| RP11-274B21.4 | 13.4 | -3.86 | 0.98 | -3.96 | 0.0001 | 0.0178 |
| NR4A1 | 237.8 | 1.14 | 0.29 | 3.95 | 0.0001 | 0.0179 |
| ALB | 73.1 | -2.48 | 0.63 | -3.94 | 0.0001 | 0.0181 |
| CIDEC | 10.7 | -4.38 | 1.11 | -3.93 | 0.0001 | 0.0181 |

| | | | | | | |
|---|---|---|---|---|---|---|
| AD000864.6 | 14.5 | 3.63 | 0.92 | 3.93 | 0.0001 | 0.0181 |
| NAPG | 1015.4 | 0.69 | 0.17 | 3.93 | 0.0001 | 0.0181 |
| FTLP3 | 511.8 | 1.42 | 0.36 | 3.93 | 0.0001 | 0.0181 |
| COBLL1 | 372.4 | -0.92 | 0.23 | -3.93 | 0.0001 | 0.0181 |
| HECW2 | 157.8 | -1.08 | 0.28 | -3.93 | 0.0001 | 0.0181 |
| DUSP3 | 8306.9 | 0.60 | 0.15 | 3.92 | 0.0001 | 0.0181 |
| CSGALNACT2 | 685.9 | 0.60 | 0.15 | 3.92 | 0.0001 | 0.0181 |
| LONRF1 | 991.1 | 0.83 | 0.21 | 3.92 | 0.0001 | 0.0181 |
| HPR | 9.8 | 5.29 | 1.35 | 3.92 | 0.0001 | 0.0181 |
| CCDC117 | 2648.7 | 0.65 | 0.16 | 3.91 | 0.0001 | 0.0183 |
| STIM2 | 736.4 | -0.71 | 0.18 | -3.90 | 0.0001 | 0.0189 |
| MARK3 | 3086.1 | -0.60 | 0.16 | -3.89 | 0.0001 | 0.0197 |
| UGT2B7 | 7.8 | -4.92 | 1.27 | -3.88 | 0.0001 | NA |
| LL0XNC01-7P3.1 | 38.1 | 1.47 | 0.38 | 3.88 | 0.0001 | 0.0204 |
| ARRDC4 | 2419.3 | 1.03 | 0.27 | 3.87 | 0.0001 | 0.0207 |
| GHR | 121.1 | -1.16 | 0.30 | -3.87 | 0.0001 | 0.0210 |
| ITPKB | 42.5 | -1.92 | 0.50 | -3.86 | 0.0001 | 0.0213 |

**Supplemental Table 12.** Genes significantly regulated by both LINC00881 overexpression and LINC0881 knockdown in human iPS cell derived cardiomyocytes

| Gene ID | log2FC (OE) | p val (OE) | log2FC (KD) | p val (KD) | Direction |
|---|---|---|---|---|---|
| LINC00881 | 2.07 | 0.0000 | -2.27 | 0.0000 | Up-Down |
| SLC16A13 | 1.07 | 0.0177 | -1.76 | 0.0244 | Up-Down |
| DOC2A | 1.34 | 0.0183 | -1.67 | 0.0131 | Up-Down |
| FAM189A1 | 0.29 | 0.0494 | -1.66 | 0.0171 | Up-Down |
| ITGB3 | 0.36 | 0.0332 | -1.41 | 0.0350 | Up-Down |
| GLB1L | 0.43 | 0.0441 | -1.40 | 0.0142 | Up-Down |
| MYH7B | 0.32 | 0.0006 | -1.25 | 0.0227 | Up-Down |
| CACNA1C | 0.36 | 0.0000 | -1.17 | 0.0001 | Up-Down |
| LARGE1 | 0.24 | 0.0130 | -1.16 | 0.0016 | Up-Down |
| FRAS1 | 0.21 | 0.0041 | -1.13 | 0.0014 | Up-Down |
| TENM4 | 0.27 | 0.0007 | -1.11 | 0.0002 | Up-Down |
| RYR2 | 0.17 | 0.0168 | -1.09 | 0.0124 | Up-Down |
| MAPK4 | 0.29 | 0.0405 | -1.08 | 0.0002 | Up-Down |
| TRAPPC9 | 0.23 | 0.0152 | -1.08 | 0.0000 | Up-Down |
| TPCN1 | 0.34 | 0.0030 | -1.06 | 0.0106 | Up-Down |
| SLIT3 | 0.26 | 0.0012 | -1.04 | 0.0002 | Up-Down |
| CACNA1D | 0.35 | 0.0002 | -1.03 | 0.0032 | Up-Down |
| RP3-412A9.16 | 1.08 | 0.0282 | -1.03 | 0.0167 | Up-Down |
| OBSCN | 0.20 | 0.0214 | -0.99 | 0.0236 | Up-Down |
| SDK2 | 0.22 | 0.0055 | -0.99 | 0.0064 | Up-Down |
| EPHB2 | 0.29 | 0.0062 | -0.98 | 0.0275 | Up-Down |
| FHOD3 | 0.13 | 0.0156 | -0.94 | 0.0008 | Up-Down |
| PLXNA4 | 0.20 | 0.0027 | -0.93 | 0.0006 | Up-Down |
| FOXN3 | 0.15 | 0.0249 | -0.91 | 0.0293 | Up-Down |
| TAP1 | 0.30 | 0.0270 | -0.90 | 0.0212 | Up-Down |
| TSPAN18 | 0.23 | 0.0394 | -0.89 | 0.0296 | Up-Down |
| MYOM2 | 0.51 | 0.0009 | -0.89 | 0.0011 | Up-Down |
| MYO18B | 0.22 | 0.0013 | -0.87 | 0.0005 | Up-Down |
| ACAD10 | 0.20 | 0.0405 | -0.85 | 0.0009 | Up-Down |
| SH3RF2 | 0.22 | 0.0118 | -0.85 | 0.0269 | Up-Down |
| ACACB | 0.26 | 0.0001 | -0.84 | 0.0145 | Up-Down |
| SH3PXD2A | 0.16 | 0.0213 | -0.83 | 0.0155 | Up-Down |
| PI4KA | 0.18 | 0.0120 | -0.79 | 0.0136 | Up-Down |
| CLYBL | 0.40 | 0.0030 | -0.77 | 0.0102 | Up-Down |
| GREB1 | 0.22 | 0.0005 | -0.77 | 0.0056 | Up-Down |
| TANC2 | 0.14 | 0.0454 | -0.75 | 0.0245 | Up-Down |
| PPARD | 0.29 | 0.0045 | -0.72 | 0.0040 | Up-Down |
| IGSF9B | 0.22 | 0.0036 | -0.70 | 0.0477 | Up-Down |
| PACS2 | 0.23 | 0.0112 | -0.70 | 0.0392 | Up-Down |

| | | | | | |
|---|---|---|---|---|---|
| FREM1 | 0.26 | 0.0011 | -0.69 | 0.0156 | Up-Down |
| ZNF76 | 0.27 | 0.0061 | -0.67 | 0.0406 | Up-Down |
| PATJ | 0.16 | 0.0479 | -0.67 | 0.0012 | Up-Down |
| POLE | 0.17 | 0.0495 | -0.67 | 0.0006 | Up-Down |
| MYOM1 | 0.14 | 0.0159 | -0.66 | 0.0064 | Up-Down |
| KCNH7 | 0.50 | 0.0005 | -0.65 | 0.0398 | Up-Down |
| PHKA2 | 0.20 | 0.0111 | -0.64 | 0.0059 | Up-Down |
| PLEKHA7 | 0.17 | 0.0193 | -0.64 | 0.0006 | Up-Down |
| FAM189A2 | 0.18 | 0.0211 | -0.63 | 0.0091 | Up-Down |
| FGFR2 | 0.17 | 0.0319 | -0.61 | 0.0274 | Up-Down |
| ANXA6 | 0.24 | 0.0049 | -0.60 | 0.0351 | Up-Down |
| CAPZB | 0.13 | 0.0417 | -0.59 | 0.0073 | Up-Down |
| MYH6 | 0.35 | 0.0000 | -0.59 | 0.0310 | Up-Down |
| TLN2 | 0.19 | 0.0104 | -0.58 | 0.0040 | Up-Down |
| TSPAN9 | 0.18 | 0.0107 | -0.57 | 0.0142 | Up-Down |
| DGLUCY | 0.18 | 0.0122 | -0.56 | 0.0067 | Up-Down |
| ITGA3 | 0.19 | 0.0460 | -0.53 | 0.0458 | Up-Down |
| MATN2 | 0.22 | 0.0279 | -0.53 | 0.0200 | Up-Down |
| LDB3 | 0.16 | 0.0045 | -0.52 | 0.0048 | Up-Down |
| MECR | 0.27 | 0.0221 | -0.52 | 0.0403 | Up-Down |
| PPP6R2 | 0.22 | 0.0206 | -0.52 | 0.0171 | Up-Down |
| LSS | 0.37 | 0.0004 | -0.51 | 0.0240 | Up-Down |
| KALRN | 0.19 | 0.0282 | -0.51 | 0.0094 | Up-Down |
| ITGB1BP2 | 0.25 | 0.0139 | -0.51 | 0.0342 | Up-Down |
| DROSHA | 0.15 | 0.0339 | -0.51 | 0.0009 | Up-Down |
| CARS2 | 0.24 | 0.0267 | -0.47 | 0.0275 | Up-Down |
| PRMT7 | 0.25 | 0.0085 | -0.46 | 0.0167 | Up-Down |
| RUSF1 | 0.30 | 0.0159 | -0.45 | 0.0447 | Up-Down |
| TNNI1 | 0.18 | 0.0499 | -0.43 | 0.0277 | Up-Down |
| PYROXD2 | 0.42 | 0.0003 | -0.42 | 0.0260 | Up-Down |
| CAMK2A | 0.37 | 0.0005 | -0.38 | 0.0165 | Up-Down |
| PREP | 0.17 | 0.0138 | -0.36 | 0.0131 | Up-Down |
| PWWP3A | 0.23 | 0.0128 | -0.33 | 0.0464 | Up-Down |
| SELENOW | 0.18 | 0.0140 | -0.28 | 0.0388 | Up-Down |
| SH3GLB1 | -0.14 | 0.0258 | 0.27 | 0.0386 | Down-Up |
| RNF2 | -0.24 | 0.0014 | 0.27 | 0.0478 | Down-Up |
| FAM222B | -0.22 | 0.0166 | 0.33 | 0.0424 | Down-Up |
| SSB | -0.20 | 0.0119 | 0.33 | 0.0237 | Down-Up |
| CHORDC1 | -0.20 | 0.0211 | 0.33 | 0.0447 | Down-Up |
| PRR13 | -0.21 | 0.0386 | 0.34 | 0.0408 | Down-Up |
| PAFAH1B2 | -0.17 | 0.0161 | 0.35 | 0.0138 | Down-Up |
| CNOT8 | -0.16 | 0.0401 | 0.35 | 0.0325 | Down-Up |
| TOMM20 | -0.13 | 0.0341 | 0.36 | 0.0010 | Down-Up |

| | | | | | |
|---|---|---|---|---|---|
| ZNF229 | -0.21 | 0.0357 | 0.36 | 0.0461 | Down-Up |
| RAB5A | -0.20 | 0.0068 | 0.36 | 0.0188 | Down-Up |
| GASK1B | -0.20 | 0.0237 | 0.37 | 0.0468 | Down-Up |
| BZW1 | -0.20 | 0.0015 | 0.37 | 0.0053 | Down-Up |
| YWHAG | -0.12 | 0.0322 | 0.38 | 0.0167 | Down-Up |
| RIT1 | -0.14 | 0.0166 | 0.38 | 0.0491 | Down-Up |
| DNAJA1 | -0.12 | 0.0431 | 0.40 | 0.0029 | Down-Up |
| PHF10 | -0.18 | 0.0156 | 0.40 | 0.0431 | Down-Up |
| EIF5 | -0.15 | 0.0035 | 0.41 | 0.0126 | Down-Up |
| ZNF697 | -0.18 | 0.0188 | 0.42 | 0.0251 | Down-Up |
| CCT6A | -0.16 | 0.0049 | 0.43 | 0.0457 | Down-Up |
| PPP4R2 | -0.28 | 0.0004 | 0.43 | 0.0459 | Down-Up |
| RMND5A | -0.18 | 0.0451 | 0.44 | 0.0009 | Down-Up |
| CGGBP1 | -0.21 | 0.0070 | 0.44 | 0.0423 | Down-Up |
| MAPK6 | -0.17 | 0.0148 | 0.44 | 0.0481 | Down-Up |
| RAB14 | -0.17 | 0.0087 | 0.45 | 0.0342 | Down-Up |
| SMAD5 | -0.18 | 0.0260 | 0.45 | 0.0279 | Down-Up |
| TERF2IP | -0.21 | 0.0016 | 0.45 | 0.0097 | Down-Up |
| MCL1 | -0.17 | 0.0015 | 0.45 | 0.0106 | Down-Up |
| SIRT1 | -0.20 | 0.0089 | 0.46 | 0.0466 | Down-Up |
| PSMD12 | -0.14 | 0.0403 | 0.46 | 0.0005 | Down-Up |
| ARF4 | -0.17 | 0.0293 | 0.47 | 0.0117 | Down-Up |
| RAD21 | -0.11 | 0.0484 | 0.47 | 0.0290 | Down-Up |
| PLEKHA3 | -0.26 | 0.0009 | 0.48 | 0.0185 | Down-Up |
| PNRC2 | -0.16 | 0.0279 | 0.48 | 0.0217 | Down-Up |
| RRM2B | -0.28 | 0.0128 | 0.48 | 0.0456 | Down-Up |
| GMFB | -0.18 | 0.0208 | 0.49 | 0.0227 | Down-Up |
| GXYLT1 | -0.24 | 0.0132 | 0.49 | 0.0130 | Down-Up |
| ZNF639 | -0.26 | 0.0007 | 0.49 | 0.0115 | Down-Up |
| SOCS4 | -0.20 | 0.0392 | 0.50 | 0.0472 | Down-Up |
| MTHFD2 | -0.19 | 0.0103 | 0.50 | 0.0405 | Down-Up |
| NAA50 | -0.22 | 0.0021 | 0.50 | 0.0047 | Down-Up |
| KIF5B | -0.19 | 0.0068 | 0.50 | 0.0026 | Down-Up |
| CCN2 | -0.18 | 0.0179 | 0.51 | 0.0329 | Down-Up |
| KLHL28 | -0.33 | 0.0013 | 0.51 | 0.0405 | Down-Up |
| SLU7 | -0.26 | 0.0005 | 0.52 | 0.0058 | Down-Up |
| SYNPO2L | -0.23 | 0.0064 | 0.52 | 0.0062 | Down-Up |
| SAMD8 | -0.31 | 0.0010 | 0.52 | 0.0211 | Down-Up |
| ZBTB10 | -0.34 | 0.0025 | 0.54 | 0.0193 | Down-Up |
| CNIH1 | -0.17 | 0.0397 | 0.54 | 0.0003 | Down-Up |
| MAP1LC3B | -0.18 | 0.0128 | 0.54 | 0.0049 | Down-Up |
| CD2AP | -0.19 | 0.0230 | 0.55 | 0.0036 | Down-Up |
| RLF | -0.20 | 0.0052 | 0.55 | 0.0165 | Down-Up |

| | | | | | |
|---|---|---|---|---|---|
| ARL5B | -0.24 | 0.0198 | 0.55 | 0.0192 | Down-Up |
| ZBTB6 | -0.30 | 0.0119 | 0.56 | 0.0322 | Down-Up |
| TOR1AIP1 | -0.19 | 0.0220 | 0.57 | 0.0114 | Down-Up |
| GNA13 | -0.19 | 0.0235 | 0.57 | 0.0124 | Down-Up |
| TWF1 | -0.21 | 0.0162 | 0.57 | 0.0035 | Down-Up |
| PNN | -0.19 | 0.0063 | 0.59 | 0.0040 | Down-Up |
| CSGALNACT2 | -0.21 | 0.0441 | 0.60 | 0.0001 | Down-Up |
| RSBN1 | -0.30 | 0.0063 | 0.61 | 0.0100 | Down-Up |
| TSC22D2 | -0.20 | 0.0244 | 0.61 | 0.0036 | Down-Up |
| TENT4B | -0.21 | 0.0173 | 0.61 | 0.0403 | Down-Up |
| SERP1 | -0.28 | 0.0007 | 0.62 | 0.0000 | Down-Up |
| CDKN2AIP | -0.26 | 0.0016 | 0.62 | 0.0430 | Down-Up |
| ATF1 | -0.17 | 0.0419 | 0.63 | 0.0358 | Down-Up |
| SLC38A2 | -0.26 | 0.0050 | 0.63 | 0.0415 | Down-Up |
| FBXO28 | -0.19 | 0.0248 | 0.64 | 0.0005 | Down-Up |
| RNF6 | -0.28 | 0.0013 | 0.65 | 0.0024 | Down-Up |
| STX3 | -0.19 | 0.0485 | 0.66 | 0.0108 | Down-Up |
| ZNF281 | -0.28 | 0.0004 | 0.66 | 0.0289 | Down-Up |
| RSL1D1 | -0.14 | 0.0283 | 0.66 | 0.0266 | Down-Up |
| ZNF24 | -0.17 | 0.0233 | 0.67 | 0.0160 | Down-Up |
| MORC3 | -0.16 | 0.0213 | 0.67 | 0.0237 | Down-Up |
| DBF4 | -0.21 | 0.0225 | 0.69 | 0.0172 | Down-Up |
| RCN1 | -0.18 | 0.0253 | 0.70 | 0.0042 | Down-Up |
| DNAJB4 | -0.24 | 0.0084 | 0.70 | 0.0185 | Down-Up |
| ZXDB | -0.20 | 0.0445 | 0.72 | 0.0480 | Down-Up |
| HBEGF | -0.26 | 0.0324 | 0.73 | 0.0032 | Down-Up |
| BCLAF1 | -0.18 | 0.0245 | 0.73 | 0.0000 | Down-Up |
| ANKRD1 | -0.22 | 0.0107 | 0.74 | 0.0141 | Down-Up |
| NUS1 | -0.20 | 0.0196 | 0.76 | 0.0000 | Down-Up |
| IRS2 | -0.35 | 0.0281 | 0.78 | 0.0466 | Down-Up |
| CEBPG | -0.17 | 0.0325 | 0.80 | 0.0257 | Down-Up |
| CCSAP | -0.29 | 0.0126 | 0.80 | 0.0304 | Down-Up |
| FOSL2 | -0.35 | 0.0218 | 0.81 | 0.0067 | Down-Up |
| EIF5A2 | -0.35 | 0.0112 | 0.82 | 0.0027 | Down-Up |
| SPTY2D1 | -0.18 | 0.0439 | 0.83 | 0.0002 | Down-Up |
| KBTBD8 | -0.28 | 0.0054 | 0.85 | 0.0027 | Down-Up |
| ARL6IP1 | -0.29 | 0.0000 | 0.86 | 0.0002 | Down-Up |
| ELOVL4 | -0.32 | 0.0028 | 0.92 | 0.0008 | Down-Up |
| RND3 | -0.25 | 0.0195 | 0.93 | 0.0043 | Down-Up |
| PHLDA1 | -0.21 | 0.0058 | 0.95 | 0.0037 | Down-Up |
| CTD-3157E16.2 | -0.38 | 0.0499 | 0.98 | 0.0178 | Down-Up |
| HEXIM1 | -0.16 | 0.0434 | 1.03 | 0.0001 | Down-Up |

| | | | | | |
|---|---|---|---|---|---|
| ARID5B | -0.43 | 0.0039 | 1.14 | 0.0015 | Down-Up |
| ATF3 | -0.32 | 0.0324 | 1.15 | 0.0033 | Down-Up |
| KLF5 | -0.70 | 0.0340 | 1.51 | 0.0009 | Down-Up |
| ANXA1 | -0.31 | 0.0127 | 1.74 | 0.0023 | Down-Up |
| ZNF252P-AS1 | -1.25 | 0.0429 | 1.81 | 0.0082 | Down-Up |
| RP11-135F9.4 | -1.04 | 0.0377 | 2.08 | 0.0113 | Down-Up |
| KLF10 | -0.49 | 0.0048 | 2.39 | 0.0000 | Down-Up |

**Supplemental Table 13.** List of qPCR primers

| Gene | Primers 5'-3' |
|---|---|
| LINC00881 | ACAGTCACGGTACTCGTTTCC |
| | TTCCCTGTCATGCCAGATCC |
| AKAP13 | ACCGGAGTTCAATGCGAGTT |
| | CACCAGCTCCTCCTGTCAAG |
| HTRA1 | AACTTTATCGCGGACGTGGT |
| | CCGGCACCTCTCGTTTAGAA |
| RPTOR | GGACCTCGTGAAGGACAACG |
| | TGACGATCACGGCGAGAATG |
| HDAC9 | AGTAAGGATGGTGGCTGTGC |
| | CGGTCTCTGTCTCCTCTTGC |
| EFCAB13 | TGGACAAGGACCTTCATACAGC |
| | CCTTGCCACTTTCATGTTCAGG |
| FBXO16 | AGCACCTGGACACCCCTAAA |
| | TGTCAAACCATTTGCCAAGCA |
| TBX3 | CGCTGTGACTGCATACCAGA |
| | GTGTCCCGGAAACCTTTTGC |
| GATA4 | GTCCTCGCCAGTCTACGTG |
| | CGCCCTGGAGGTAGGACA |
| HAND2 | ACTTCCATGGCTGGCTCATC |
| | ATACTCGGGGCTGTAGGACA |
| TBX5 | AGAATATCCCGTGGTCCCCA |
| | GACTCGCTGCTGAAAGGACT |
| TNNT2 | AGAGGAGGAGGAGCTCGTTT |
| | CTCCTTCTCCCGCTCATTCC |
| MYH6 | AGATAGAGAGACTCCTGCGGC |
| | TCGGTCATCTTGGTGCTTCC |
| MYH7 | CTCGCTTCGGCAGCACA |
| | AACGCTTCACGAATTTGCGT |
| FHL2 | GGAGTTGGGGAGACTGGTTG |
| | GATTCGTTGCAATGGTGGCA |
| SYNPO2L | ACCTGGATGAAAAGCCTCGG |
| | GTCTTGTAACTCCTGGCCCC |
| CACNA1C | GCTTATGGGGCTTTCTTGCAC |
| | ACTGGACTGGATGCCAAAGG |
| RYR2 | GAGCCAGTGTCATCCACCAA |
| | AGGTGGCTGAAAGAATGAGCA |
| 18S | GTAACCCGTTGAACCCCATT |
| | CCATCCAATCGGTAGTAGCG |
| U6 | CTCGCTTCGGCAGCACA |
| | AACGCTTCACGAATTTGCGT |

**Supplemental Table 14.** List of LNA GapmeR sequences

| Name | Sequence 5'-3' |
|---|---|
| LINC00881 | A*G*A*A*C*A*G*G*C*A*G*G*A*G*G*T |
| Negative Control | A*A*C*A*C*G*T*C*T*A*T*A*C*G*C |
